# NW-flex: flexible-block sequence alignment for short tandem repeats

**DOI:** 10.64898/2025.12.22.695990

**Authors:** Zhezhen Yu, Dan Levy

## Abstract

Aligning sequencing reads to short tandem repeats (STRs) is challenging: the number of repeat copies in a read often differs from the reference, and small changes inside and around the repeat can lead to many competing alignments. We introduce NW-flex, a simple extension of classical sequence alignment that addresses this problem.

NW-flex takes a reference with a designated internal block and, while allowing that block to contract to any shorter substring, identifies the best-scoring alignment. Apart from this substring choice, the alignment obeys the usual substitution scoring and gap constraints. For STRs, we design the flexible block with enough repeat copies to accommodate any expected repeat count. The alignment then contracts the reference to match the read while the unique flanks remain fully constrained.

NW-flex requires a small modification to the Needleman–Wunsch/Gotoh algorithm, adds minimal computational overhead, and preserves optimality with respect to classical scoring. We provide open-source Python and Cython implementations, a set of worked examples, and notebooks that reproduce the figures and validate correctness against baseline alignments.

## 1. Introduction

**Short tandem repeats** (**STRs**, also called microsatellites) are genomic sequences built from motifs of 1–6 base pairs repeated in tandem and collectively account for 3% of the human genome [3]. STRs are mutation hotspots: repeat copy-number changes are frequent and substitutions within tracts are common [11], resulting in either **pure** (only motif bases), **interrupted** (tracts that include single-nucleotide variants, SNVs), or **compound** (adjacent tracts with different motifs) repeats. This variability makes STRs useful in forensics, population genetics, and disease studies [12], but it complicates alignment by creating many equally plausible choices.

Standard dynamic-programming alignment, from Needleman–Wunsch [7] to affine-gap Gotoh [4] and including local/semi-global variants [2, 10], treats every sequence position equivalently. At STR loci this uniformity blurs the boundary between fixed flanks and the repeat tract: expansions, contractions, and internal mismatches can produce multiple alignments that trade a gap for a shifted boundary, especially when the repeat is imperfect or the flanks carry variants.

Some STR callers avoid full alignment by using short, unique signatures from the left and right flanks. They search each read for both signatures and infer repeat length from the distance between the two matches. Tools such as lobSTR and GangSTR augment this approach with motif-aware scoring and locus-specific priors on allele lengths [5, 6]. This strategy works well when the flanks remain distinctive and well conserved, but polymorphism, sequencing errors, and adjacent repeats can make these matches weak or ambiguous. Graph-based mappers offer an alternative by encoding repeat structures in a variation graph and aligning reads by path search [9]. This approach handles length variation naturally, but requires a predefined graph and more infrastructure than a single pass over a linear reference.

We take a different approach. Consider a reference *X* = *A* · *Z* · *B*, where *A* and *B* are flanking sequences surrounding a designated internal block *Z*. Given a read *Y*, we want an alignment that treats the flanks normally but allows *Z* to contract to any shorter contiguous substring *Z*^∗^, with the choice of *Z*^∗^ ⊆ *Z* determined by the alignment itself. It is true that STRs both expand and contract relative to the reference; however, we only need contraction. By choosing *Z* longer than any expected repeat length, every allele appears as a substring of *Z*.

A naive solution — enumerate all substrings, align each, and take the best — requires *O*(|*Z*|^2^) separate alignments. **NW-flex** achieves the same result in a single pass. Conceptually, NW-flex extends semi-global alignment (which allows prefix/suffix skipping at the sequence ends) to operate on *internal* blocks. It accomplishes this by adding “shortcut” connections that let the alignment skip any prefix and any suffix of *Z*, effectively exploring all possible contractions simultaneously.

The scoring scheme is unchanged and the computational cost is the same as classical alignment with a constant-factor overhead.

### Guarantee

*NW-flex returns* max_*Z*∗⊆*Z*_ NWG(*A* · *Z*^∗^ · *B, Y*), *where NWG denotes the standard Needleman–Wunsch/Gotoh alignment score [4, 7]*.

Full algorithmic details are given in Supplement S1 and a proof of the guarantee in Supplement S2; here we focus on the conceptual framework and its application to STRs.

## 2. The NW-flex algorithm

We now describe how NW-flex works. Readers familiar with dynamic programming (DP) for sequence alignment will recognize the standard Needleman–Wunsch/Gotoh (NWG) framework; those less familiar can follow the intuition here, with Notebook 1 (`Alignment_Basics`) providing a self-contained primer. We have included a static, printable version of the notebooks as an additional supplement.

### Setup and terminology

We write the reference as *X* = *A* · *Z* · *B*, where *A* and *B* are fixed flanking sequences and *Z* is the designated flexible block (Figure 1A). Standard alignment fills in a grid where each cell (*i, j*) represents aligning the first *i* bases of *X* to the first *j* bases of the read *Y*. Cells are filled row by row, with each row depending only on the row immediately above. We call the last row of flank *A* the **leader row** and the first row of flank *B* the **closer row**. The flexible block *Z* occupies the rows between them.

**Figure 1.**
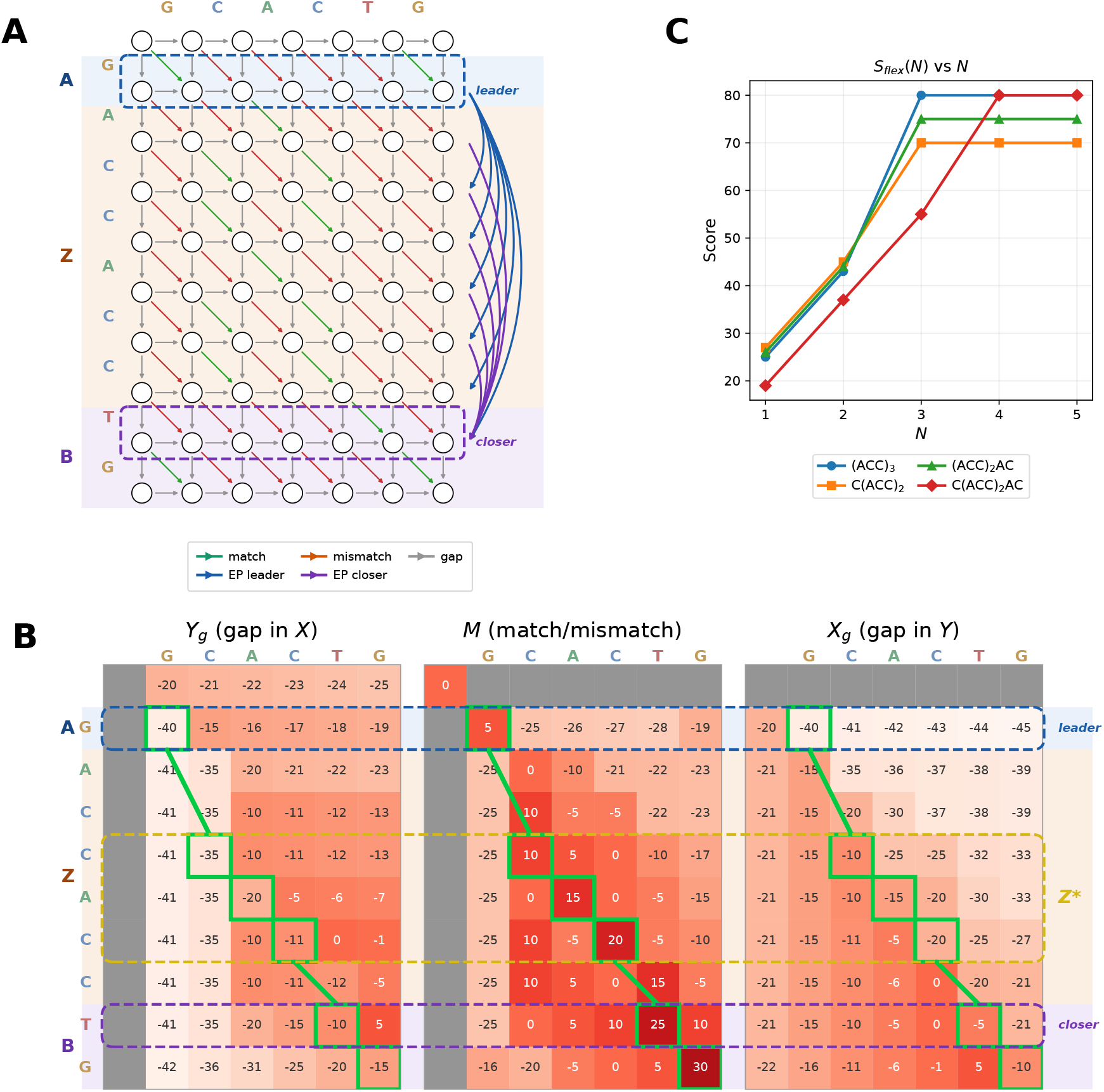
NW-flex alignment for a small example. The reference is *X* = G · ACCACC · TG (*X* = *A* · *Z* · *B*) and the read is *Y* = `GCACTG`. Colored bands indicate the flanks (*A, B*) and flexible block (*Z*). **(A) Alignment graph.** Standard algorithms connect only adjacent rows. NW-flex adds *extra predecessors* that allow the alignment to skip portions of *Z*: blue edges connect the leader row to rows in *Z*; purple edges connect rows in *Z* to the closer row. **(B) DP matrices**. The green path shows the optimal alignment. Where the path jumps over rows (via shortcut edges), portions of *Z* are skipped. The rows traversed define the contracted substring *Z*^∗^, giving the effective reference *A Z*^∗^ · *B*. **(C) STR plateau. (C) STR application**. For STR alignment, we design the flexible repeat block *Z* = *R*^*N*^ to contain many copies of the repeat motif. If *N* is too small, the best alignment score is suboptimal (e.g., *N* = 1, 2), but once *N* is large enough, the best alignment score is attained and levels off as *N* increases further.

### Extra predecessors

In standard alignment, each row depends only on its immediate predecessor. NW-flex adds **extra predecessor** (EP) relationships such that:

- Every row inside *Z* has one EP: the leader row.
- The closer row has many EPs: the leader row *and* every row inside *Z*.

These extra predecessors let the alignment skip any prefix of *Z* (by jumping from the leader to a row inside *Z*) and skip any suffix (by jumping from anywhere inside *Z* directly to the closer). The effect is that all possible contractions *Z*^∗^ ⊆ *Z* are explored in a single pass. Figure 1A shows these connections as colored edges: blue from the leader into *Z*, purple from *Z* to the closer. Figure 1B shows the resulting alignment matrices, with the optimal path (green) jumping over unused portions of *Z*.

### Traceback

Traceback proceeds as usual, except that some steps may jump across multiple rows via the extra connections. Where the path enters *Z* from the leader marks the start of the chosen substring *Z*^∗^, and where it exits to the closer marks the end. The rows traversed define *Z*^∗^, which together with the flanks gives the optimal effective reference *A* · *Z*^∗^ · *B*. In Figure 1B, the best flex alignment has *Z*^∗^ = `CAC`.

### Complexity

NW-flex has the same time complexity as standard alignment, *O*(|*X*||*Y* |), with a small constant-factor overhead from checking extra predecessors. In principle, memory can be reduced to *O*(|*Y* |) by retaining the previous row, the current row, and one row of memory for each leader and closer. Our current implementations store the full matrix to simplify traceback. Full recurrences, boundary conditions, and pseudocode are given in Supplement S1.

## 3. Aligning tandem repeats

For STR loci, we take the flexible block to be a repeated motif: *Z* = *R*^*N*^, where *R* is the repeat unit (e.g., `ACC`) and *N* is chosen larger than any expected repeat length in the read. The flanks *A* and *B* anchor the locus, and NW-flex contracts *Z* to whatever substring best matches the read. Because we select *N* large enough to cover all expected alleles, this effectively explores all plausible repeat copy numbers.

### Avoiding redundant alignments

A subtlety arises with tandem repeats: because the motif repeats, the same alignment can be achieved at multiple positions within *Z*. For example, if the read contains three copies of `ACC`, that match could be placed at positions 1–3, or 2–4, or 3–5 of a longer reference and each alignment would have the same score.

To avoid this redundancy, we restrict where alignments can exit the repeat block: specifically, only within the last motif-length portion of *Z*. Any alignment that would have exited earlier can slide forward by whole motif copies without changing its score, so no optimal solutions are lost. Details are in Supplement S3.

### Score plateau

As the reference copy number *N* increases, the best alignment score is non-decreasing and eventually plateaus. This follows from a simple observation: every substring of *R*^*N*^ is also a substring of *R*^*N*+1^, so adding copies to the reference can only help or make no difference. Once *N* is large enough to contain whatever repeat content the read has, further increases in *N* will not improve the score. This is shown in Figure 1C for four example reads with different partial-repeat content.

Within the DP matrices, this property also creates a banded pattern of non-decreasing scores over each motif-length segment of rows. As *N* grows, these bands eventually stabilize across all columns, indicating that extending the read block further cannot improve the best path.

This observation suggests a practical optimization: by monitoring score changes while filling out the DP matrices, we can stop our alignment over *Z* early once the scores stabilize across a full motif-length segment, and advance directly to the closer row. This avoids unnecessary computation and offsets our use of a long reference to accommodate large expansions. This is not currently implemented in our package, but it is a natural direction for future work. Supplement S3 illustrates these properties with examples.

### Compound and interrupted repeats

More complex loci, such as compound repeats (adjacent blocks with different motifs) or interrupted repeats (motifs containing variants), are handled by combining multiple flexible blocks. Example configurations appear in Notebook 4.

## 4. Implementation & availability

An open-source implementation of NW-flex, including code, tests, and notebooks, is available at https://github.com/nwflex/nwflex.

The package provides two backends with a shared interface. The pure Python implementation mirrors the algorithm described above and is intended for readability and experimentation. A compiled Cython/NumPy backend targets performance-sensitive use and is up to 200 − 500*×* faster on typical inputs.

High-level functions expose three modes: standard Needleman–Wunsch/Gotoh (no flexible block), single-block alignment, and STR alignment with the closer row restriction. These building blocks can be composed to create references with multiple flexible blocks. The current implementation supports global and semi-global boundary conditions; local alignment may be added in a future release. The repository also includes plotting utilities, STR helper functions, and notebooks that illustrate the concepts and reproduce the paper’s examples.

## 5. Validation

We validated NW-flex against two independent baselines. First, when no flexible block is designated, NW-flex reduces to standard Needleman–Wunsch/Gotoh; we confirmed exact score agreement with an independent implementation. Second, for references with a flexible block, we compared NW-flex to the naive method that enumerates all substrings *Z*^∗^ ⊆ *Z*, aligns each separately, and takes the maximum. NW-flex matched the naive optimum in all cases.

We tested hand-constructed edge cases, thousands of randomized sequences, multiple scoring schemes, and STR-specific configurations. The Python and Cython backends produce identical results. All validation code and a `pytest` suite are included in the repository; details are in Supplement S4.

## 6. Discussion

NW-flex extends classical sequence alignment by optimizing over all contractions of a designated internal block while preserving standard substitution and gap scoring. We designed this alignment method for STR loci, where flanks often carry locus identity but repeat copy-number varies. In this setting, we place an overlong repeat block between the flanks and NW-flex selects the contraction that best matches the read.

This approach has practical limitations. First, it assumes a reference model for the locus: we must identify repeat elements, choose boundaries, and decide how to represent imperfect repeats. In our workflow we use Tandem Repeat Finder [1] to locate candidate repeat tracts, then choose a maximal region and a simplest-repeat unit to form an idealized motif reference. We score the reference against its idealized version to bound expected alignment scores from reads. Second, the overlong block increases compute; however this computational cost is linear with repeat length and the monotone-plateau behavior in *N* suggests an approach to mitigate this cost by monitoring scores for early termination. Finally, NW-flex does not directly encode priors or penalties on repeat length. While not currently implemented, such preferences can be included by assigning costs to extra predecessor connections.

Our immediate motivation came from aligning partially bisulfite-converted reads at C-rich STR loci [13]. In that assay, half of the cytosines are randomly converted to uracil before amplification, and are later read as thymine. The interruption of perfect repeats preserves repeat copy-number during amplification, but the mutations also reduce sequence complexity and introduce dense, structured mismatches across both the repeat and its flanks. We wanted a simple scoring scheme that treats C-to-T as a match, while still enforcing that reads anchor to the locus-defining flanks.

In our first analyses of this assay, we relied on flank-signature matching. We searched for short sequences in the flanks that remained unique at the locus after C-to-T conversion. As we expanded to population data, this strategy began to break down. Polymorphism and reference-to-sample differences can mutate or erase candidate signatures, and conversion further compresses sequence complexity, making the remaining signatures less distinctive. These effects turn locus identification into a brittle thresholding problem. NW-flex resolves this tension. It supports permissive scoring while still anchoring alignments to the flanks, yielding results that degrade smoothly as signal is lost rather than failing at a hard threshold.

While STRs motivate this work, the extra-predecessor mechanism applies more generally. By choosing where additional predecessor relationships are allowed, NW-flex can optimize over a structured family of related references while retaining standard substitution and gap scoring. Multiple flexible blocks can model compound or interrupted repeats, and similar constructions can represent other bounded internal variation, such as optional exons in isoform-aware transcript alignment.

Some settings may require mutually exclusive alternatives, where taking one path should preclude taking another. Conceptually, this can be expressed by *removing* selected standard predecessor links so that incompatible alternatives cannot be combined within a single alignment path. Taken together, these extensions make NW-flex a composable modification to classical alignment for exploring structured variation with standard scoring and traceback.

## Supporting information

Supplementary Notebooks

## 7. Acknowledgments

## Use of large language models

We used LLMs to assist with manuscript formatting, code documentation, and code development, as detailed in Supplement S5.

## 8. Supplementary Material

### S1 Full NW-flex dynamic programming details

We record the boundary conditions and recurrences underlying NW-flex and summarize them in Algorithm S1.

#### S1.1 Notation

Let *X* = *X*_1..*n*_ and *Y* = *Y*_1..*m*_ denote the reference and read. The reference decomposes as *X* = *A* · *Z* · *B*, with flanks *A, B* and flexible block *Z*. We set *s* = |*A*| and *e* = *s* + |*Z*|, so that *A* = *X*_1..*s*_, *Z* = *X*_*s*+1..*e*_, and *B* = *X*_*e*+1..*n*_. In the DP grid, row *s* is the **leader row** (last row of *A*) and row *e*+1 is the **closer row** (first row of *B*).

We use the standard three-layer Gotoh formulation for affine-gap alignment: *Y*_*g*_(*i, j*) tracks alignments ending with a gap in *X, M* (*i, j*) tracks alignments ending with a match or mismatch, and *X*_*g*_(*i, j*) tracks alignments ending with a gap in *Y*. We write σ(·, ·) for the substitution score matrix, *g*_*s*_ for gap-start (gap-open), and *g*_*e*_ for gap-extend.

NW-flex augments the standard recurrences using **extra predecessor sets** *E*(*i*) ⊆ {0, …, *i*−1} for each row *i*. In the single-block case:

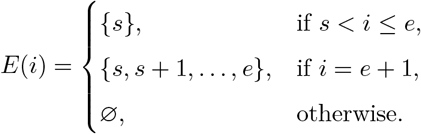

With this notation established, we now give the full recurrences.

#### S1.2 Initialization and boundary conditions

We use the standard global-alignment initialization, with the modification that extra predecessors only affect gaps in *Y* (the *X*_*g*_ layer). For 0 ≤ *i* ≤ *n* and 0 ≤ *j* ≤ *m* we set

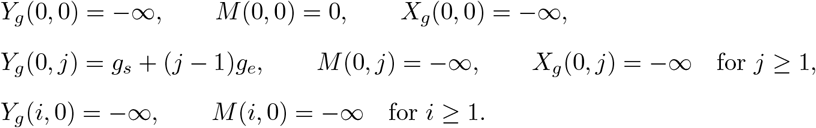

The first column of *X*_*g*_ is initialized by a baseline NWG update from the previous row *i* − 1, followed by an optional refinement using the extra predecessor set *E*(*i*) ⊆ {0, …, *i* − 1}:

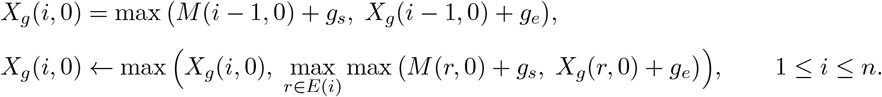

If *E*(*i*) = ∅, this reduces to the standard Gotoh initialization in the first column.

##### Algorithm S1

NW-flex DP via per-row extra predecessors

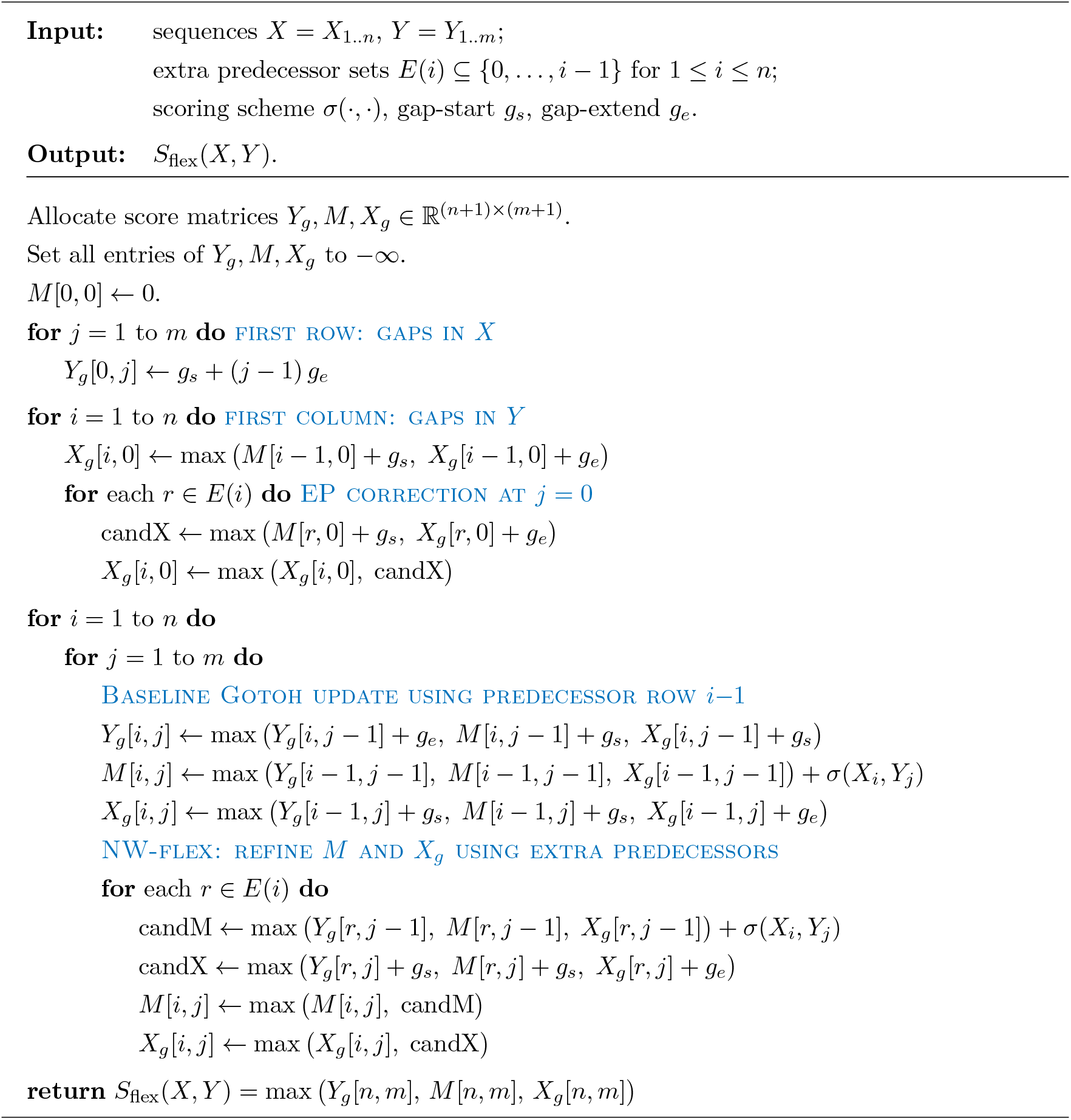

Other boundary conventions (local or semi-global) are obtained in the usual way by changing the initialization of the first row and column and by modifying which cells are eligible as final states. For example, a one-sided semi-global alignment that does not penalize leading gaps in *X* sets the *X*_*g*_ layer of column 0 to 0 for all rows; in the NW-flex view this corresponds to treating row 0 as a universal leader for the first column.

Trailing gaps in *X* can be handled by allowing the alignment to terminate in any row *r* of the last column and taking max_*r*_ max(*Y*_*g*_(*r, m*), *M* (*r, m*), *X*_*g*_(*r, m*)) as the final score. Equivalently, we may think of a virtual terminal row *n*+1 with a terminal predecessor set *E*_term_ ⊆ {0, …, *n*}; standard global alignment uses *E*_term_ = {*n*}, while a local-in-*X* semi-global scheme uses *E*_term_ = {0, …, *n*}. These boundary choices are orthogonal to the internal flex-block structure: once the first row/column and the terminal rows are fixed, NW-flex simply augments the interior recurrences via the sets *E*(*i*), independently of whether the surrounding scheme is global, semi-global, or local.

#### S1.3 NW-flex recurrence updates

##### Baseline Gotoh update

For 1 ≤ *i* ≤ *n* and 1 ≤ *j* ≤ *m* we first perform the usual three-state Gotoh update using the immediate predecessor row *i* − 1:

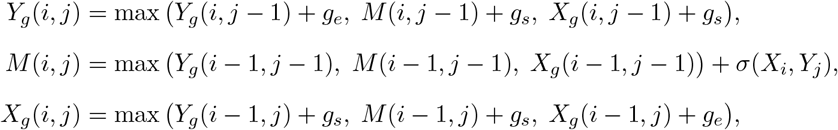

##### Extra-predecessor refinement

The effect of NW-flex is entirely captured by augmenting the predecessor set for *M* and *X*_*g*_. For each row *i* and each extra predecessor *r* ∈ *E*(*i*), we compute alternative candidates candM(*i, j*; *r*) and candX(*i, j*; *r*) using row *r* in place of row *i* − 1:

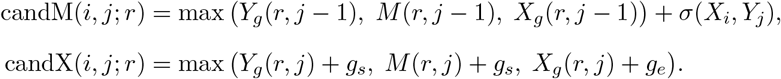

We then refine the baseline values by

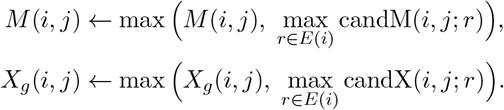

#### S1.4 Traceback

In the default global setting we take

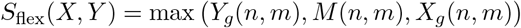

and start traceback from the maximizing state at (*n, m*). In semi-global variants we instead select the terminal row from the prescribed set *E*_term_ ⊆ {0, …, *n*} (as described above) and start traceback from the maximizing state at (*r*^∗^, *m*).

In our reference implementation we additionally record *row jumps* during traceback: whenever an *M* or *X*_*g*_ step uses a predecessor row *r i* − 1, we log the transition *r* → *i* at column *j* as a jump induced by an extra predecessor. Along any best path, these jumps mark the entry into and exit from the flexible block *Z*; the corresponding row indices give the interval of rows in *Z* that are actually used in the alignment. This provides a direct way to reconstruct the realized substring *Z*^∗^ (and its location inside *Z*) from the traceback, in addition to the aligned sequences themselves.

### S1.5 Graphical alignment DAGs

Figure S1 gives a graphical view of the recurrences described above, following the three-state gadget representation of affine-gap alignment in Pachter and Sturmfels [8]. Each grid position (*i, j*) is expanded into a local three-state gadget with in-ports and out-ports corresponding to the *Y*_*g*_, *M*, and *X*_*g*_ layers; internal edges encode gap-open versus gap-extend penalties, and horizontal, vertical, and diagonal edges connect gadgets between neighboring cells. In the Gotoh panel the only inter-cell edges are between adjacent rows and columns, realizing the baseline three-layer DP.

**Figure S1.**
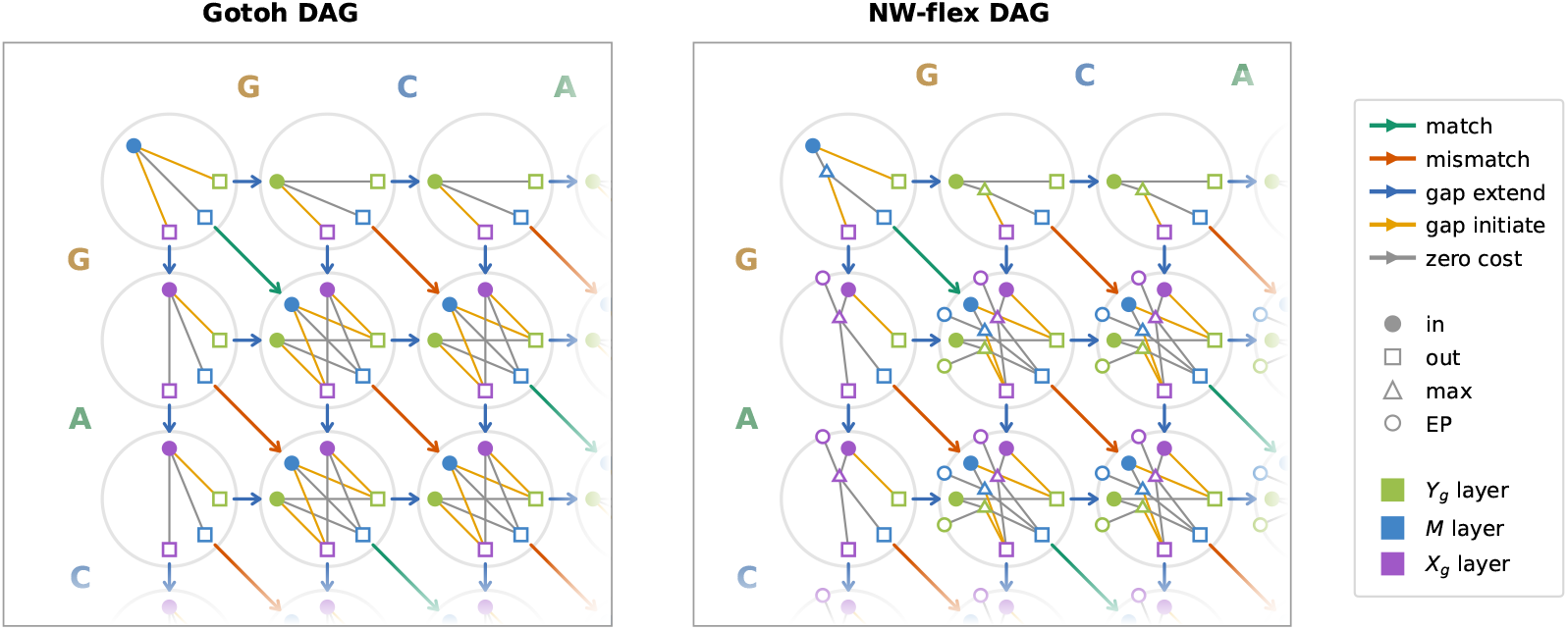
Graphical alignment DAGs for affine-gap Gotoh and NW-flex. Left: Gotoh DAG for a small example, with each grid cell expanded into a local three-state gadget. Colored edges correspond to match/mismatch, gap-initiate, and gap-extend transitions; gray edges denote zero-cost internal wiring, and node shapes indicate in-ports, out-ports, and max (DP) states. Right: the corresponding NW-flex DAG for the same example. The local gadget is augmented with an extra-predecessor (EP) in-port on each row. This EP port represents the max over all logical edges from predecessor rows *r* ∈ E(i) in the same column, so the diagram remains readable while capturing how NW-flex adds row-to-row shortcuts on top of the standard affine-gap structure.

In the NW-flex panel, the local gadget is extended by additional *EP input ports* on each row. These blank circles stand in for the set of all extra predecessor row input ports in the *same* column and layer. The triangular *max-port* takes as input both the usual predecessor port (from row *i*−1) and the EP ports and passes the maximum value to the *M* and *Xg* layers. The *Y g* layer is unchanged, as extra predecessors do not affect gaps in *Y*. This graphical representation captures the essence of NW-flex while keeping the diagram readable, by summarizing the potentially many extra-predecessor edges into a single edge from an EP port per row.

## S2 Flex guarantee: NW-flex as a union over substrings

We briefly justify the guarantee

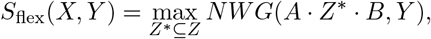

for the single-block pattern *X* = *A* · *Z* · *B* with extra-predecessor sets

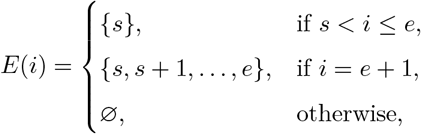

where *s* = |*A*| and *e* = *s* + |*Z*| as in Supplement S1.

### View as a union of NWG DAGs

For each contiguous substring *Z*^∗^ = *X*_*u*+1..*v*_ ⊆ *Z*, consider the standard NWG alignment DAG for the pair (*X*^*′*^, *Y*) = (*A* · *Z*^∗^ · *B, Y*). Its vertices can be identified with a subset of the NW-flex grid: rows 0, …, *s* align *A*, rows *u* + 1, …, *v* align the chosen substring of *Z*, and rows *e* + 1, …, *n* align *B*. The prefix *X*_*s*+1..*u*_ and suffix *X*_*v*+1..*e*_ of *Z* are absent in *X*^*′*^ and must be skipped.

The extra-predecessor edges in NW-flex do exactly this skipping at zero cost in *X*: the edge pattern *E*(*i*) = {*s*} for *s < i* ≤ *e* allows us to jump from the leader row *s* down to an arbitrary row *u* + 1 inside *Z*, thereby skipping *X*_*s*+1..*u*_. Similarly, the pattern *E*(*e* + 1) = {*s*, …, *e*} allows us to jump from any row *v* inside *Z* to the closer row *e* + 1, skipping *X*_*v*+1..*e*_. All other edges coincide with the standard NWG transitions.

Thus, for each choice of substring *Z*^∗^ there is a one-to-one correspondence between paths in the NWG DAG for *A* · *Z*^∗^ · *B* and paths in the NW-flex DAG that (i) enter *Z* at row *u* + 1 via an edge from row *s*, (ii) stay within rows *u* + 1, …, *v*, and (iii) leave *Z* at row *v* via an edge to row *e* + 1. The substitution and gap penalties are unchanged, so each path has exactly the same score in both graphs.

### Optimality over all substrings

Conversely, any path in the NW-flex DAG from (0, 0) to (*n, m*) that uses extra predecessors can enter the block *Z* only once and leave it only once, because row indices are non-decreasing along any path. The entry and exit rows determine a contiguous interval *X*_*u*+1..*v*_ ⊆ *Z*, and restricting the path to this interval yields a valid path in the corresponding NWG DAG for *A* · *X*_*u*+1..*v*_ · *B* with the same score.

Taken together, the NW-flex DAG is the union of the NWG DAGs for all effective references *A* · *Z*^∗^ · *B*. Running the usual max-weight dynamic program on this union therefore computes

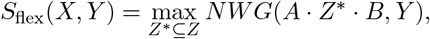

as claimed.

## S3 Aligning STRs with NW-flex

We provide formal details for the STR specialization introduced in Section 3. Recall that *Z* = *R*^*N*^ is a tandem repeat of a motif *R* of length *k*. We choose *N* large enough to cover the expected range of repeat lengths in the read, so that any plausible repeat segment can be realized as a contraction of *Z*. We modify the extra-predecessor set for the closer row to

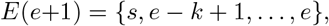

while keeping *E*(*i*) = {*s*} for *s < i* ≤ *e* as in Supplement S1.

### S3.1 Phase classes and phased repeats

#### Equivalent positions in a tandem repeat

Because *Z* = *R*^*N*^ consists of identical copies of *R*, every *k*-th row of the DP grid looks the same: row *s* + 1 aligns *R*_1_, row *s* + 1 + *k* also aligns *R*_1_, and so on. We say two rows have the same *phase* if they correspond to the same position within a motif copy.

This situation creates potential redundancy: an alignment using some substring of *Z* will have the same score as one using a shifted copy of that substring (offset by w hole motifs), p rovided the reference is long enough to contain both. Since we choose *N* larger than any expected repeat length, we can restrict exits to the last *k* rows of the block (plus the leader, for alignments that skip *Z* entirely) without losing any optimal solutions.

#### Describing a substring of *R*^*N*^

A contiguous substring *Z*^∗^ ⊆ *R*^*N*^ may start or end mid-motif. We parameterize it by (*a, b, M*) where:

1. *a* ∈ {0, …, *k*−1}: entry phase (a suffix of *R*),
2. *M* ≥ 0: number of complete motif copies,
3. *b* ∈ {0, …, *k*−1}: exit phase (a prefix of *R*).

Writing suf(*R, a*) for the last *a* bases of *R* and pre(*R, b*) for the first *b* bases:

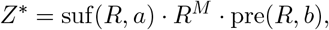

with total length *a* + *Mk* + *b*. For example, with *R* = ACT: the string T(ACT)_2_AC has *a* = 1, *M* = 2, *b* = 2.

#### Repeat content

For *Z*^∗^ to appear as a substring of *R*^*N*^ requires a sufficiently la rge va lue fo r *N*. We define the *repeat content* of *Z*^∗^ as

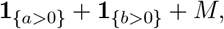

counting one for each partial motif (entry and/or exit) plus *M* complete copies. The condition *N* ≥ repeat content ensures *Z*^∗^ fits w ithin *R*^*N*^.

### S3.2 Monotonicity in copy number and phase classes

#### Monotone in copy number

Write *S*_flex_(*N*) for the best flex score when the reference contains *N* copies of the motif. Every substring of *R*^*N*^ is also a substring of *R*^*N*+1^, so the candidate contractions grow with *N*. More candidates means the best score cannot decrease:

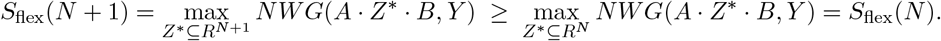

Once *N* reaches the repeat content of the read’s repeat segment, the optimal substring fits entirely in *R*^*N*^ and further increases in *N* add only redundant substrings. The score plateaus.

Figure 1C illustrates this for four example reads with different phase parameters (*a, b, M*). Each curve increases and then flattens when *N* reaches the repeat content. This behavior is precisely why we select *N* larger than the longest expected repeat content of the reads.

#### Phase-wise monotonicity in the DP matrix

A fine grained version of this pattern appears within the DP matrix itself. Group the rows of *Z* by phase: rows at the same offset modulo *k* correspond to the same position within a motif copy. Fix a column *j* and move down through the rows of the same phase (i.e., row *s* + *i, s* + *i* + *k, s* + *i* + 2*k*, …). The score in each Gotoh layer is non-decreasing along this sequence and (given the global boundary conditions) eventually plateaus. Figure S2 illustrate this for the match/mismatch layer *M* (*i, j*). For each phase class and each column, the scores rise and then level off.

**Figure S2.**
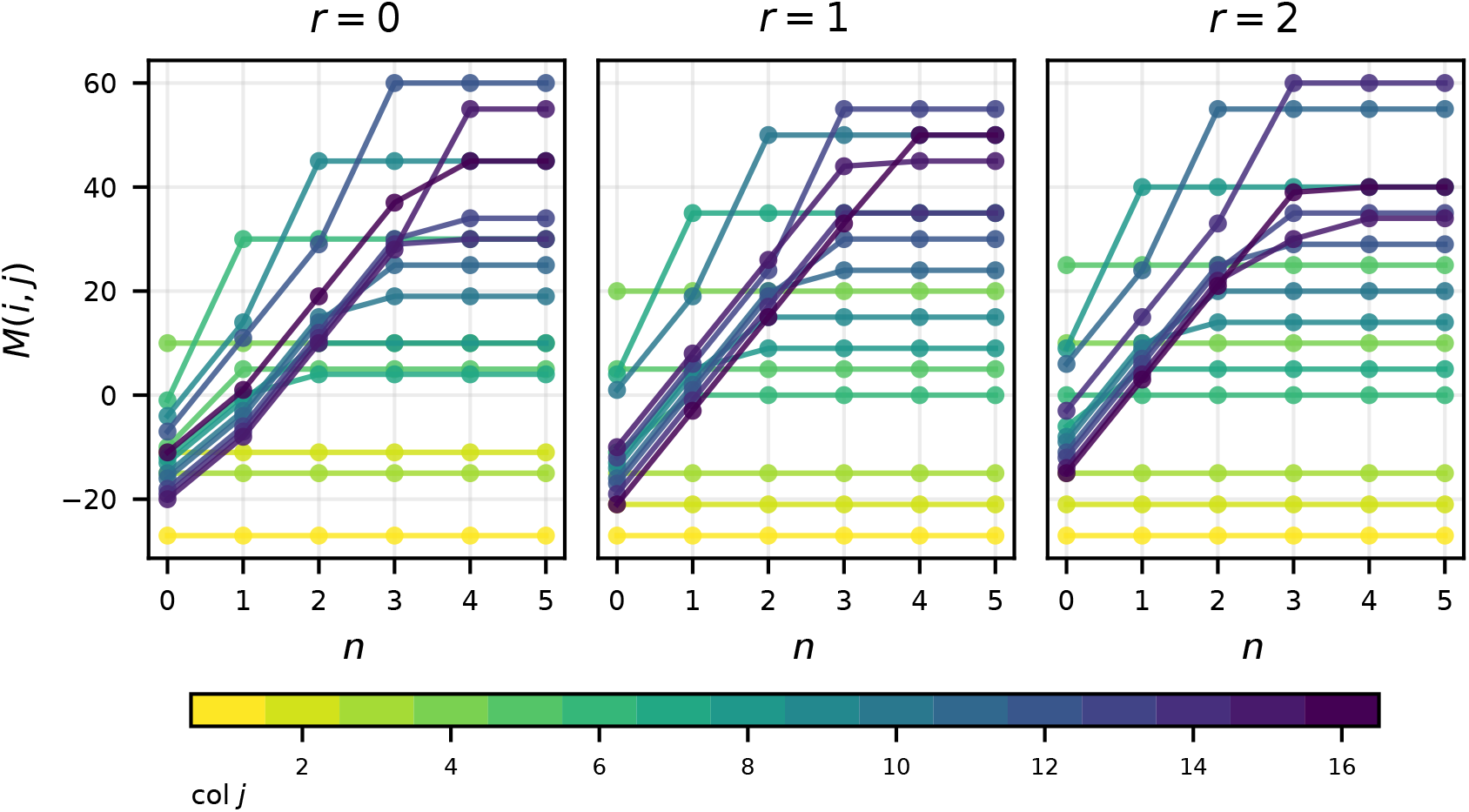
DP monotonicity. Phase-wise scores in the match/mismatch layer *M* (*i, j*) for an STR locus with motif length *k* = 3. For each motif phase *r* ∈ {0, 1, 2} (separate panels) and column *j* (color scale), we plot *M* (*i*_*n*_, *j*) against the repeat index *n* along rows *i*_*n*_ = *s* + 1 + *r* + *nk*. Within each phase class and column, these trajectories are non-decreasing and approach a plateau as *n* increases.

#### Possible optimization

This plateau behavior suggests an early-stopping criterion: once scores have stabilized across all phase classes at a given column, further rows cannot improve the result. We do not pursue this optimization here, but it may be useful for performance-critical implementations. In particular, we have selected a large value for *N* to cover all expected repeat lengths; early stopping could offset the cost of this choice by avoiding unnecessary computation in the long tail of the DP matrix.

### S4 Implementation and validation details

The reference implementations follow the recurrences in Section S1 directly. The pure Python version stores the three Gotoh layers (*Y*_*g*_, *M, X*_*g*_) and the corresponding traceback information in NumPy arrays and iterates over rows *i* and columns *j*, performing a baseline update from row *i* − 1 followed by the per-row extra-predecessor refinement over *r* ∈ *E*(*i*). The C/NumPy implementation uses the same layout but pushes the DP initialization and recurrence updates into C for efficiency; both implementations share the same interface and test suite.

We validated the NW-flex implementation through three complementary approaches: optimality verification against a naive baseline, STR phase correctness, and Cython implementation equivalence.

#### S4.1 Optimality verification

To confirm that NW-flex computes the flex optimum

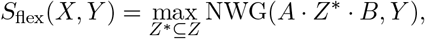

we compared scores against a naive baseline that explicitly enumerates all contiguous substrings *Z*^∗^ ⊆ *Z* and computes the standard Needleman–Wunsch/Gotoh score for each candidate *A* · *Z*^∗^ · *B*.

Validation proceeded in stages:

1. **Base case**. With *E*(*i*) = ∅ for all rows, NW-flex reduces to standard Gotoh alignment. We verified score equality against an independent NWG implementation on test cases covering identical sequences, single deletions, substitutions, and mixed indels.
2. **Single-block flex**. For the *A* · *Z* · *B* configuration, we verified that NW-flex achieves the naive maximum on (i) six hand-crafted test cases exercising contractions, complete block skipping, insertions within *Z*, and partial repeat matching; (ii) 6000 mutated variants, with 15% substitution rate and 8% indel rate; and (iii) 1000 fully randomized *A* · *Z* · *B* inputs.
3. **Alignment validity**. Beyond score equality, we verified that aligned sequences have equal length and that removing gap characters recovers the original sequences.
4. **Scoring robustness**. We repeated the randomized tests under four different scoring schemes varying gap-open and gap-extend penalties, with 1000 random tests per scheme. All tests passed.

#### S4.2 STR phase validation

For the STR-specific EP pattern, we verified correctness across all valid phase combinations (*a, b, M*) satisfying the constraint **1**_{*a>*0}_ + **1**_{*b>*0}_ + *M* ≤ *N*. For each combination, we constructed the read *Y* = *A* · *Z*^∗^ · *B* where *Z*^∗^ = suf(*R, a*) · *R*^*M*^ · pre(*R, b*), and verified that NW-flex achieves the perfect-match score |*Y* | *×* match_score.

We also verified phase inference from traceback: the recovered (*a, b, M*) values computed from the entry and exit row jumps matched the ground-truth parameters for all test cases. Additionally, we confirmed the phase-wise monotonicity property by checking that *L*(*i, j*) ≤ *L*(*i* + *k, j*) holds for all phases *r* ∈ {0, …, *k*−1}, columns *j*, and Gotoh layers *L* ∈ {*Y*_*g*_, *M, X*_*g*_}.

#### S4.3 Cython implementation equivalence

The Cython implementation was validated against the reference Python implementation. For both standard NW (empty EP) and single-block configurations, we verified identical alignment scores, identical aligned sequences (*X*_aln_ and *Y*_aln_), identical row jumps recorded during traceback, and identical DP matrices (*Y*_*g*_, *M, X*_*g*_) at all reachable cells. The Cython implementation achieves speedups of up to 200–500*×* over pure Python while maintaining exact numerical agreement.

### S5 Use of large language models

We used large language models (LLMs) to assist with manuscript formatting, code documentation, and code development. Primary uses were for editing prose, generating well-formatted LaTeX, and writing docstrings. Copilot exposed plotting parameters while developing figures. The algorithm, proofs, and imaginations are our own.

