## Supplementary Notebooks for "NW-flex: flexible-block sequence alignment for short tandem repeats"

December 22, 2025

### Supplementary Notebooks for NW-flex

```
# PDF build configuration - vector graphics
%config InlineBackend.figure_formats = ['pdf', 'png']
import matplotlib
matplotlib.rcParams['figure.dpi'] = 150
```

import sys print(sys.version) print(sys.executable)

### 1 Notebook 1: Introduction to Alignment

This notebook reviews the classical Needleman–Wunsch algorithm in a form that matches the notation used in the NW-flex paper. We work with two sequences

$$X = X_1 \dots X_n, \qquad Y = Y_1 \dots Y_m,$$

and a scoring scheme that assigns values to matches, mismatches, and gaps.

#### 1.1 Overview

The global alignment problem is:

> Given sequences $X$ and $Y$, find an alignment (with gap symbols "–") that maximizes a prescribed scoring function.

The Needleman–Wunsch algorithm solves this by:

1. Representing prefixes $(X_{1..i}, Y_{1..j})$ as nodes $(i, j)$ in a grid.
2. Defining a recurrence for the optimal score $F(i, j)$ in terms of its three predecessors $(i-1, j-1)$, $(i-1, j)$, and $(i, j-1)$.
3. Filling the DP matrix $F$ and tracing back from $(n, m)$ to $(0, 0)$ to recover one optimal alignment.

This notebook is organized as follows:

1. **Sequences and scoring.** Introduce the notation for $X$, $Y$, and the match/mismatch/gap scoring scheme.
2. **The alignment DAG.** Represent alignments as paths in a directed acyclic graph.
3. **Simple Needleman–Wunsch dynamic program.** Derive and visualize the DP recurrence and traceback for linear gaps.
4. **Gotoh: affine gap penalties (short recap).** Introduce the three-state DP $(Y_g, M, X_g)$ and the $K_{3,3}$ gadget.

We begin with the simpler case where every gap has a single fixed cost (a linear gap penalty). This lets us focus on the alignment graph and how dynamic programming works step by step. At the end of the notebook, we return to gaps and allow them to be more expensive to start than to extend (often called affine gap penalties). We show how Gotoh's method handles this by keeping three separate score tables that record whether the alignment currently ends in a match, a gap in the reference, or a gap in the read.

##### 1.3 Setup and imports

```
#  Notebook magic: autoreload modules
%load_ext autoreload
%autoreload 2

import matplotlib.pyplot as plt
from nwflex.nw_basics_plot import (
    NWDagPlotter,
    plot_example_alignments,
    plot_dp_initialization,
    run_dp_fill_demo,
    plot_filled_dag,
    plot_optimal_path
)

from nwflex.nw_basics import nw_gotoh_matrix
from nwflex.plot import (
    draw_affine_k33_dag,
    plot_gotoh_matrices
)
```

##### 1.4 Sequences and scoring

We start with two short sequences and a simple scoring scheme:

- match $= +5$
- mismatch $= -5$
- gap $= -3$

This keeps the numbers small and makes it easy to follow by hand.

**Note:** If 2*gap > mismatch, the optimal alignments will not use mismatches since they could instead use two gaps.

###### 1.4.1 Example sequences

```
X = "GATT"
Y = "GACT"

print("X:", X)
print("Y:", Y)
```

```
X: GATT
Y: GACT
```

##### 1.4.2 Match, mismatch, and gap penalties

```
MATCH = 5
MISMATCH = -5
GAP = -3

def pair_score(a: str, b: str) -> int:
    """Return the match/mismatch score for a pair of bases."""
    return MATCH if a == b else MISMATCH

print("MATCH   =", MATCH)
print("MISMATCH=", MISMATCH)
print("GAP     =", GAP)
```

```
MATCH   = 5
MISMATCH= -5
GAP     = -3
```

#### 1.5 The alignment DAG

We view global alignment as finding a path in a directed acyclic graph (DAG). Let

$$X = X_1 \dots X_n, \qquad Y = Y_1 \dots Y_m.$$

We construct a grid of nodes indexed by integer pairs $(i, j)$ with $0 \le i \le n$ and $0 \le j \le m$, where node $(i, j)$ represents the prefixes

$$X_{1..i} = X_1 \dots X_i, \qquad Y_{1..j} = Y_1 \dots Y_j.$$

From each node $(i, j)$ we add three types of directed edges:

| | | | |
|---|---|---|---|
| **diagonal:** | $(i, j) \to (i+1, j+1)$ | align $X_{i+1}$ with $Y_{j+1}$ | (match or mismatch), |
| **vertical:** | $(i, j) \to (i+1, j)$ | align $X_{i+1}$ with a gap in $Y$ | (insertion in $X$), |
| **horizontal:** | $(i, j) \to (i, j+1)$ | align a gap in $X$ with $Y_{j+1}$ | (insertion in $Y$). |

A global alignment corresponds to a path from the source node $(0, 0)$ to the sink node $(n, m)$.

##### 1.5.1 Drawing the DAG

```
# Create a plotter
plotter = NWDagPlotter(X, Y, match=MATCH, mismatch=MISMATCH, gap=GAP)
fig, ax = plt.subplots(1, 1, figsize=(6, 5.5))
grid = plotter.draw_grid(ax, edge_alpha=0.9)
fig.suptitle(f"Alignment DAG for X={X}, Y={Y}")
```

```
# Add legend with arrows at the bottom
from matplotlib.lines import Line2D
legend_handles = [
    Line2D([0], [0], color='green', lw=1, marker='>', markersize=4,␣
 ↪label='match'),
    Line2D([0], [0], color='red', lw=1, marker='>', markersize=4,␣
 ↪label='mismatch'),
    Line2D([0], [0], color='blue', lw=1, marker='>', markersize=4, label='gap'),
]
legend = ax.legend(
    handles=legend_handles,
    loc='upper center',
    bbox_to_anchor=(0.5, -0.02),
    ncol=3,
    fontsize=12,
    frameon=True,
    edgecolor='black',
    fancybox=False,
    borderpad=0.6,
)
legend.get_frame().set_linewidth(1.0)
plt.tight_layout()
plt.show()
```

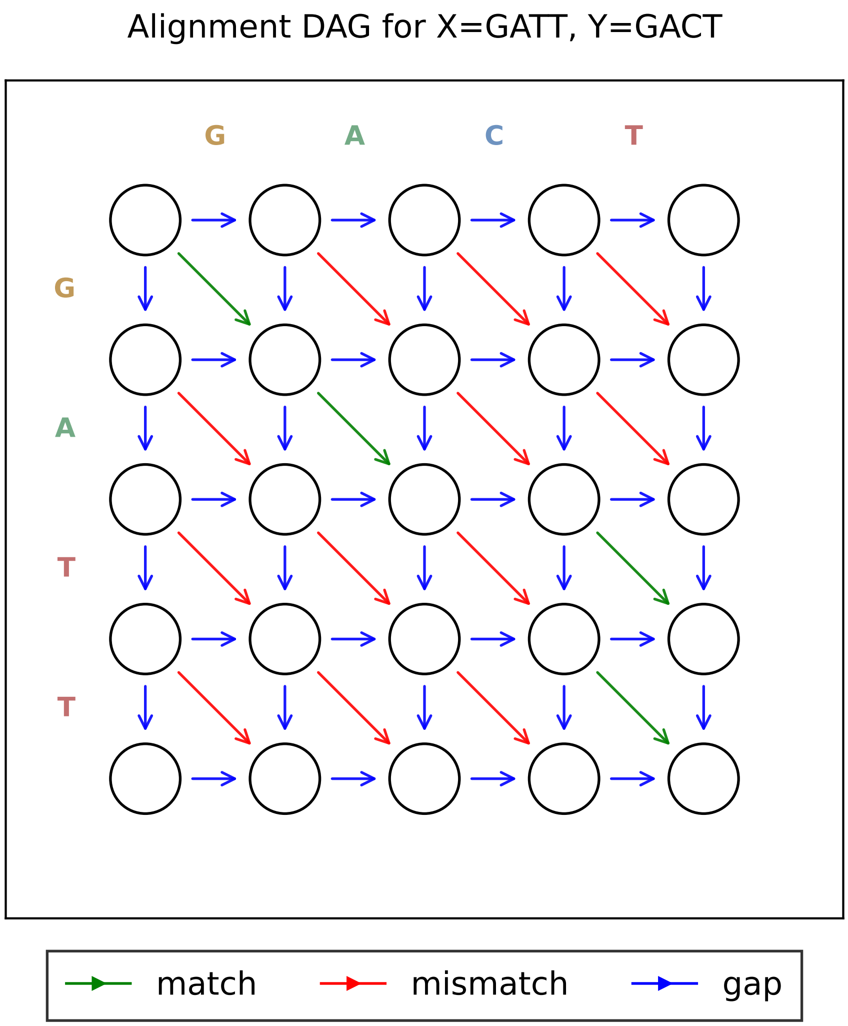

The DAG visually encodes the same three move types: diagonal matches/mismatches, vertical gaps in $Y$, and horizontal gaps in $X$. The score of a path is the sum of edge weights.

##### 1.5.2 Example alignments as paths

Before we introduce the NW algorithm, we first look at a few concrete examples of alignments and their paths in the DAG. The alignment DAG is built so that every possible global alignment between $X$ and $Y$ appears as some path from $(0,0)$ to $(n,m)$. By drawing a few example alignments and then tracing their paths in the DAG, we make this correspondence explicit: each step in the path (diagonal, vertical, horizontal) matches exactly one step in the gapped alignment (match/mismatch, gap in $Y$, gap in $X$).

```
# Define three different paths (alignments) through the DAG
# X = "GATT", Y = "GACT"

# Path 1: All diagonal (direct alignment, one mismatch T vs C)
```

```
path1 = [(0, 0), (1, 1), (2, 2), (3, 3), (4, 4)]

# Path 2: Insert gap in Y early, shift alignment
path2 = [(0, 0), (1, 0), (2, 1), (3, 2), (4, 3), (4, 4)]

# Path 3: Insert gap in X to skip the T-C mismatch
path3 = [(0, 0), (1, 1), (2, 2), (2, 3), (3, 4), (4, 4)]

# Plot all three paths using the helper function
fig, axes = plot_example_alignments(
    X, Y,
    paths=[path1, path2, path3],
    titles=["All diagonal (1 mismatch)", "Gaps at ends", "Gap the mismatch"],
    match=MATCH, mismatch=MISMATCH, gap=GAP,
)
```

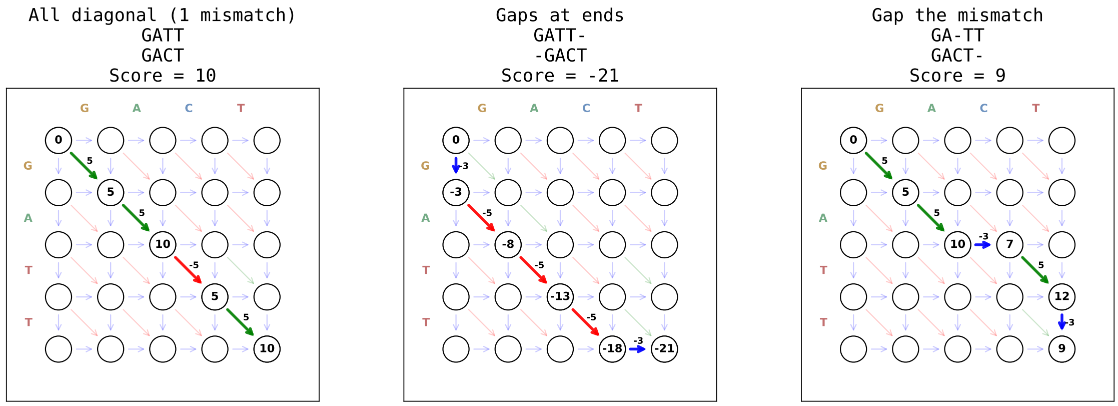

The figure above shows three different paths through the DAG, each corresponding to a different alignment of $X =$ `GATT` and $Y =$ `GACT`:

- **Left panel (score=10)**: All diagonal moves. This aligns every position directly: G–G (match, +5), A–A (match, +5), T–C (mismatch, -5), T–T (match, +5). Total: $5+5-5+5 = 10$.
- **Center panel (score=-21)**: This path takes a vertical move first (gap in Y), then proceeds diagonally before a final horizontal move. The shifted alignment causes more mismatches and gaps, resulting in a poor score.
- **Right panel (score=9)**: This path inserts a gap in X (horizontal move) to skip the C in Y, avoiding the T–C mismatch. The alignment is GA-TT / GACT- with score $5+5-3+5-3 = 9$.

Next, we will see how the Needleman–Wunsch algorithm systematically identifies the best path by computing optimal scores for all prefix pairs $(X_{1..i}, Y_{1..j})$ using dynamic programming.

##### 1.6 Simple Needleman–Wunsch dynamic program

We now implement a basic global alignment DP with a single penalty per gap.

Let $F(i, j)$ denote the best score for aligning $X_{1..i}$ with $Y_{1..j}$. The recurrence is

$$F(i,j) = \max\Big\{F(i-1,j-1) + s(X_i, Y_j),\ F(i-1,j) + \text{gap},\ F(i,j-1) + \text{gap}\Big\},$$

where

- $s(X_i, Y_j)$ is the match/mismatch score for aligning $X_i$ with $Y_j$,
- `gap` is the penalty for inserting a gap in either sequence.

###### 1.6.1 Initialization (boundary conditions)

Before filling interior cells, we must initialize the boundaries:

$$F(0,0) = 0$$

$$F(i,0) = i \cdot \text{gap} \quad \text{for } 1 \le i \le n,$$

$$F(0,j) = j \cdot \text{gap} \quad \text{for } 1 \le j \le m.$$

These correspond to aligning a prefix of one sequence against an all-gap prefix of the other, with each gap incurring the same cost.

```
# Visualize the initialization steps
fig_init, axes = plot_dp_initialization(X, Y, match=MATCH, mismatch=MISMATCH,
    gap=GAP)
```

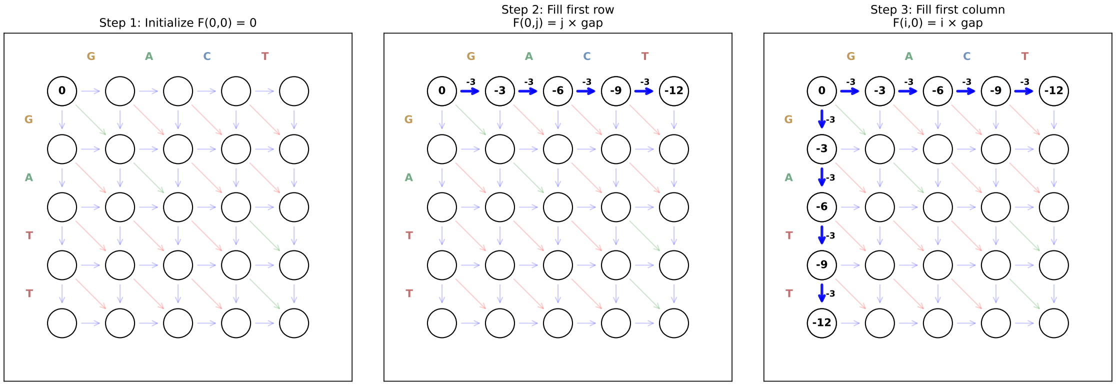

The figure above shows how we initialize the first row and column of the DAG. Once these boundary cases are filled in, every interior node has exactly three possible incoming edges, so the same update rule applies uniformly to all remaining entries. With boundaries initialized, we can now fill the interior cells using the recurrence.

##### 1.6.2 Filling an interior cell

We compare the three possible ways to reach $(1, 1)$:

$$
\begin{array}{rlllll}
\textbf{from left:} & (1,0) \to (1,1) & F(1,0) + \text{gap} & = -3 + (-3) & = -6 & \text{gap } Y : (\texttt{G}, \texttt{-}), \\
\textbf{from diagonal:} & (0,0) \to (1,1) & F(0,0) + s(X_1, Y_1) & = 0 + 5 & = +5 & \text{match} : (\texttt{G}, \texttt{G}), \\
\textbf{from above:} & (0,1) \to (1,1) & F(0,1) + \text{gap} & = -3 + (-3) & = -6 & \text{gap } X : (\texttt{-}, \texttt{G}).
\end{array}
$$

By definition, $F(i, j)$ is the best score over all paths from $(0, 0)$ to $(i, j)$. Any such path must arrive at $(i, j)$ from one of its three predecessors, so its score can be no better than the best of these three candidate scores. Taking the maximum of the three therefore enforces the "best over all paths" definition of $F(i, j)$.

The maximum is 5 (diagonal), so $F(1, 1) = 5$.

The function `run_dp_fill_demo` wraps this logic for all cells: it iterates over the DP grid, computes the three candidates at each $(i, j)$, writes the best score into `F[i, j]`, records which move was chosen, and produces the panel-style visualization.

**Note:** Change `show_first_n` and `show_last_m` in the call to `run_dp_fill_demo` to visualize different portions of the DP fill.

```
# Run and visualize the DP fill process
dp_result = run_dp_fill_demo(X, Y, match=MATCH, mismatch=MISMATCH, gap=GAP,
                             show_first_n=2, show_last_m=2)
```

```
DP fill: visualizing first 2 and last 2 of 16 interior cells
```

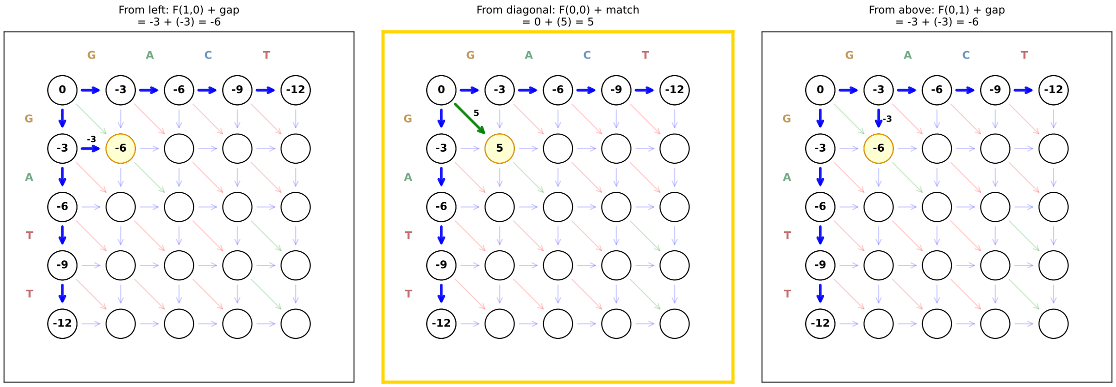

```
Cell (1, 1): best_move='diag', best_score=5
```

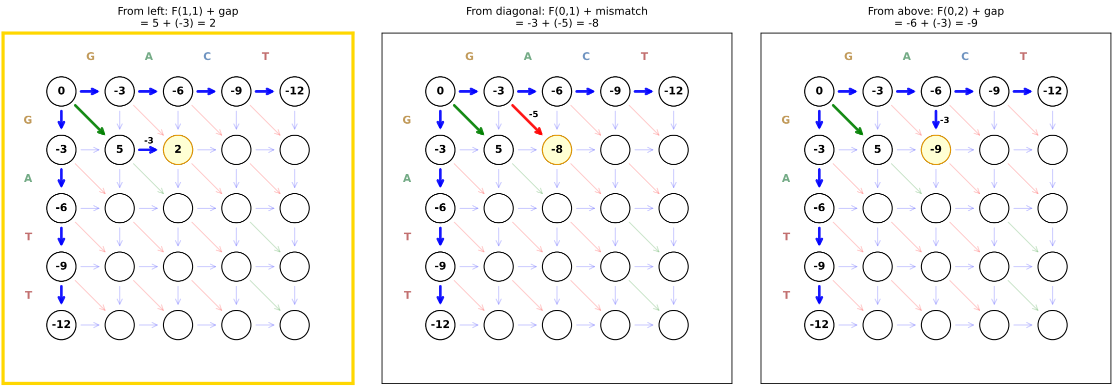

Cell (1, 2): best_move='left', best_score=2

... (silently filling 12 intermediate cells) ...

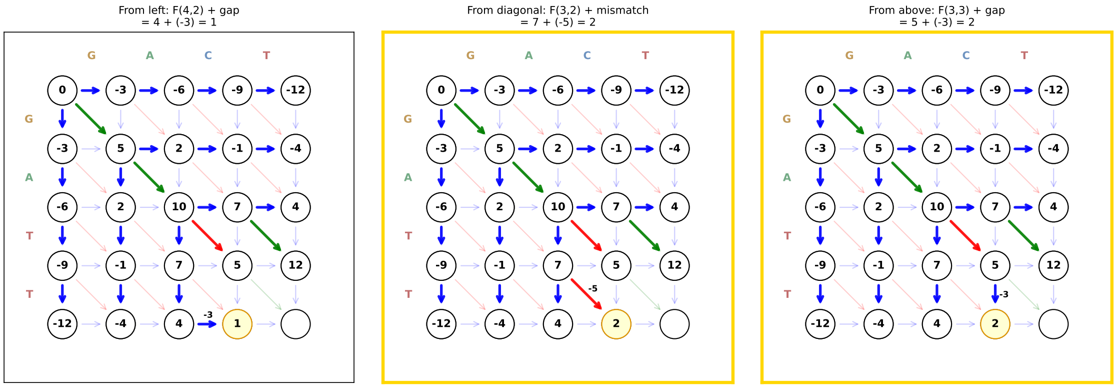

Cell (4, 3): best_move='diag', best_score=2

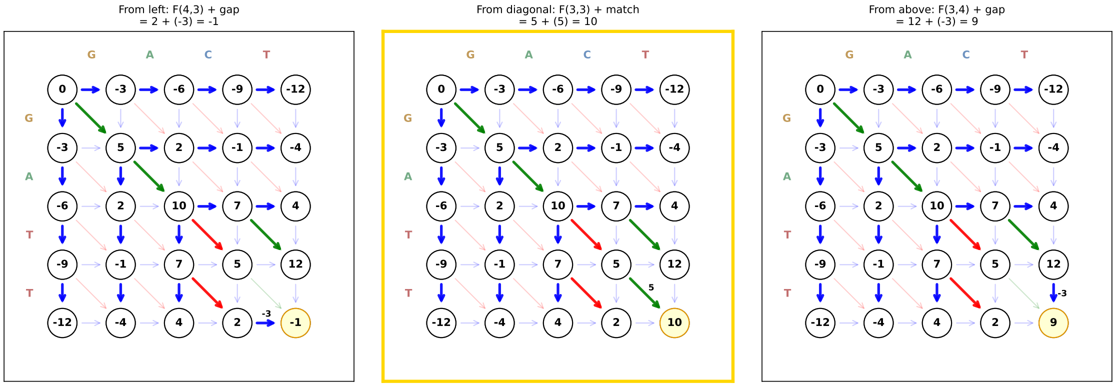

```
Cell (4, 4): best_move='diag', best_score=10

*** Final alignment score: F[4,4] = 10 ***
```

##### 1.6.3 Multiple optimal paths

Sometimes more than one incoming edge gives the same best score at a cell. In the panel above, cell $(4, 3)$ can be reached with score 2 both from the diagonal and from above. The DP table only needs the *value* of $F(i, j)$, so ties do not change the scores; they only affect which path we recover during traceback. In this demo we record a single predecessor at each cell, so the visualization shows one optimal path, not all of them. A different tie-breaking rule would produce a different, but equally good, optimal alignment.

```python
# Plot the fully filled DAG with all traceback edges
fig, ax = plot_filled_dag(
    X, Y, dp_result['F'], dp_result['traceback_edges'],
    match=MATCH, mismatch=MISMATCH, gap=GAP,
    plotter=dp_result['plotter'], figsize=(6, 5)
)
```

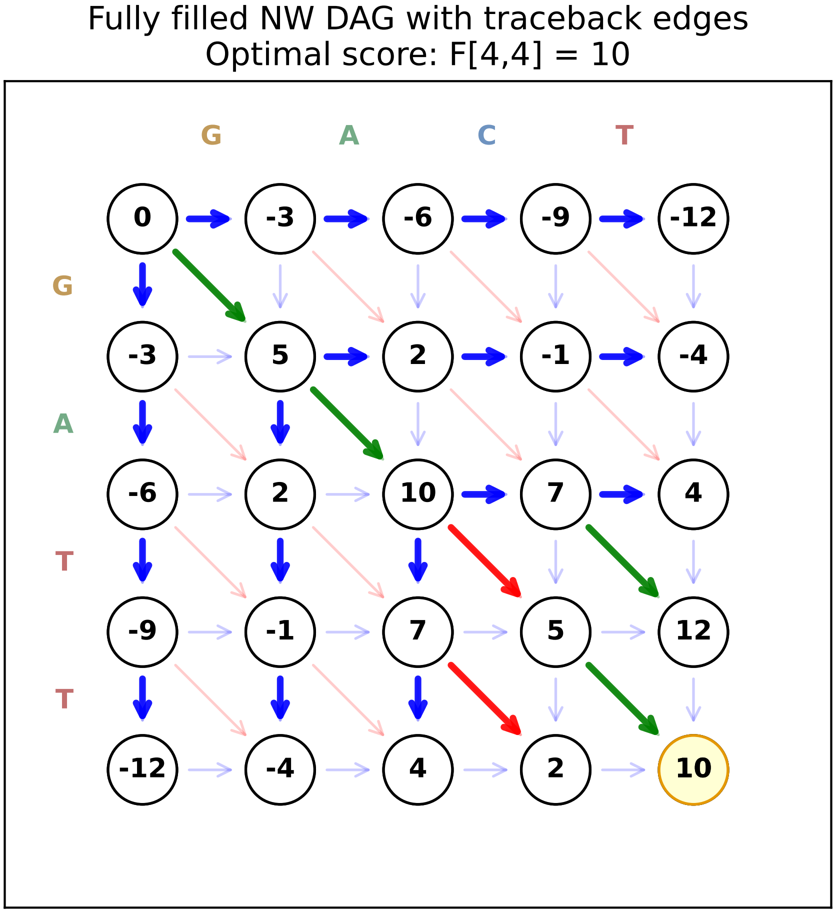

```
Final F matrix:
[[  0  -3  -6  -9 -12]
 [ -3   5   2  -1  -4]
 [ -6   2  10   7   4]
 [ -9  -1   7   5  12]
 [-12  -4   4   2  10]]
```

##### 1.6.4 Traceback: recovering the optimal alignment

The traceback edges recorded during the DP fill represent a best predecessors for each cell. To find *one* optimal alignment, we trace back from $(n, m)$ to $(0, 0)$ following these edges. This gives us a single path through the DAG corresponding to one optimal alignment.

The figure below highlights this **optimal path** in red, showing the exact sequence of moves that yields the best alignment score.

```python
# Plot the optimal path and recover the alignment
fig, ax, alignment_info = plot_optimal_path(
    X, Y, dp_result['F'], dp_result['traceback_edges'],
    match=MATCH, mismatch=MISMATCH, gap=GAP,
    plotter=dp_result['plotter']
)
```

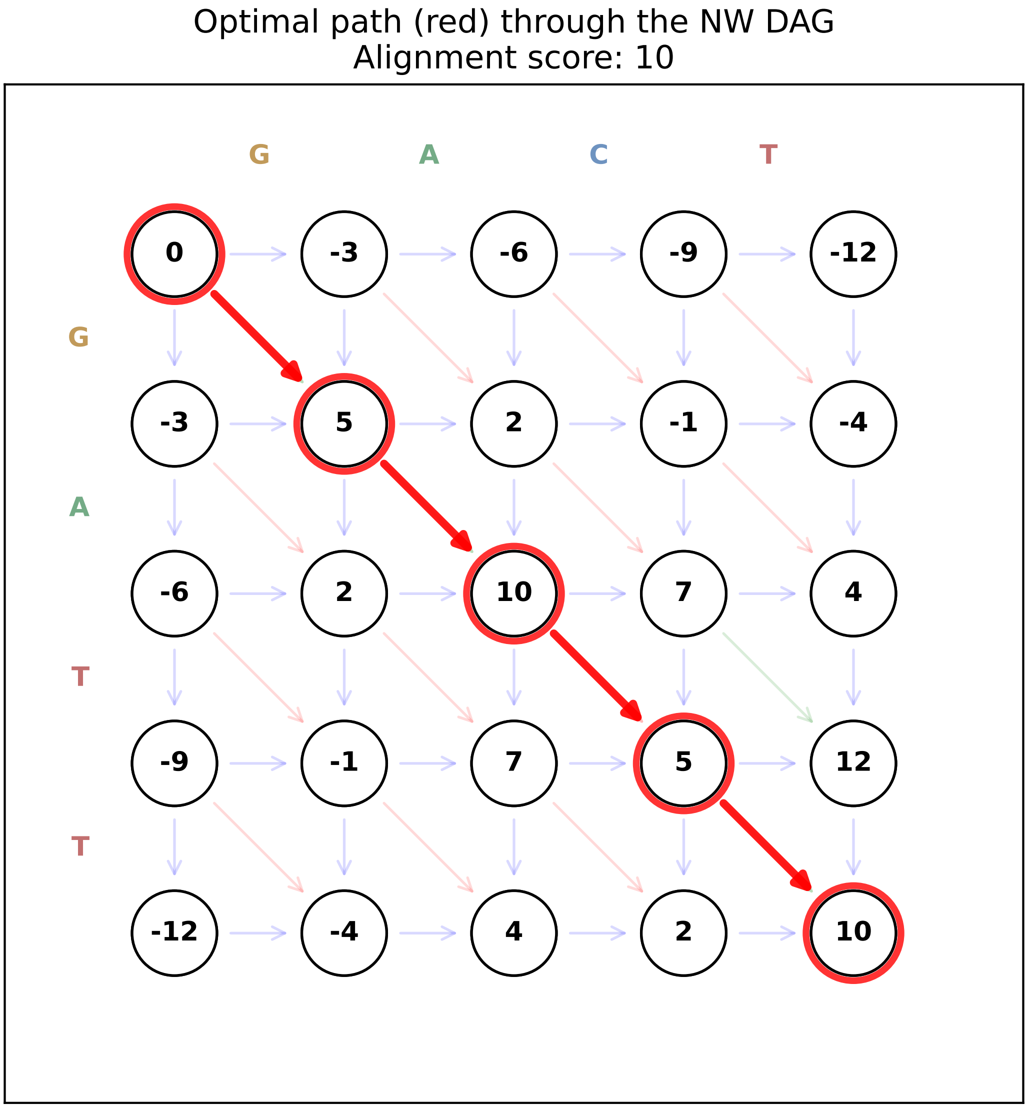

```
Optimal alignment recovered from traceback:
  X: GATT
  Y: GACT
     ||x|
     (|=match, x=mismatch, space=gap)
```

##### 1.7 Gotoh: NW with affine gap penalties

So far we used a single DP matrix $F$ with a **linear gap penalty**, where each gap character costs the same amount. This means ten separate 1-bp gaps have the same total cost as one 10-bp deletion, even though in real sequences long contiguous insertions/deletions are common while starting a new gap is rarer. To reflect this, it is more realistic to use an **affine gap penalty**, which makes opening a gap expensive but extending an existing gap cheaper.

Gotoh's affine-gap formulation keeps the global-alignment setup but tracks three DP layers at each cell $(i, j)$:

- $Y_g(i, j)$: best score ending with a gap in $X$ (horizontal move),
- $M(i, j)$: best score ending with a match or mismatch (diagonal move),
- $X_g(i, j)$: best score ending with a gap in $Y$ (vertical move).

In this section we will:

1. run the Gotoh algorithm on an example pair of sequences,
2. briefly explain why these three layers are needed for affine gaps, and
3. summarize the three-state recurrences and their $K_{3,3}$ gadget view.

##### 1.7.1 Example sequences

For the affine-gap example we use longer sequences that will reappear in the next NW-flex notebook:

- $X =$ `GATTACA`
- $Y =$ `GTTCA`

```
# Example sequences for affine-gap (Gotoh) alignment
X = "GATTACA"
Y = "GTTCA"

print("Affine-gap example:")
print("X:", X)
print("Y:", Y)
```

```
Affine-gap example:
X: GATTACA
Y: GTTCA
```

##### 1.7.2 Scoring and affine gap penalty

For this affine-gap example we use the same basic scoring scheme as in the NW-flex notebooks:

- match: $+5$
- mismatch: $-5$
- gap start $g_s$: $-20$
- gap extend $g_e$: $-1$

A contiguous gap of length $k$ then has total cost

$$\text{gap\_cost}(k) = g_s + (k-1)g_e.$$

These values make a single long gap cheaper (per base) than breaking it into many short gaps, mirroring the idea that one long indel event is more likely than many independent small ones.

```
# Scoring scheme
from nwflex.validation import get_default_scoring
from nwflex.plot import plot_score_system
```

```
score_matrix, gap_open, gap_extend, alphabet_to_index = get_default_scoring()

# Display the default score scheme:
clean_map = {str(k): v for k, v in alphabet_to_index.items()}
fig = plot_score_system(
    score_matrix, gap_open, gap_extend, alphabet_to_index,
    figsize=(5, 3.5)
)
plt.show()
```

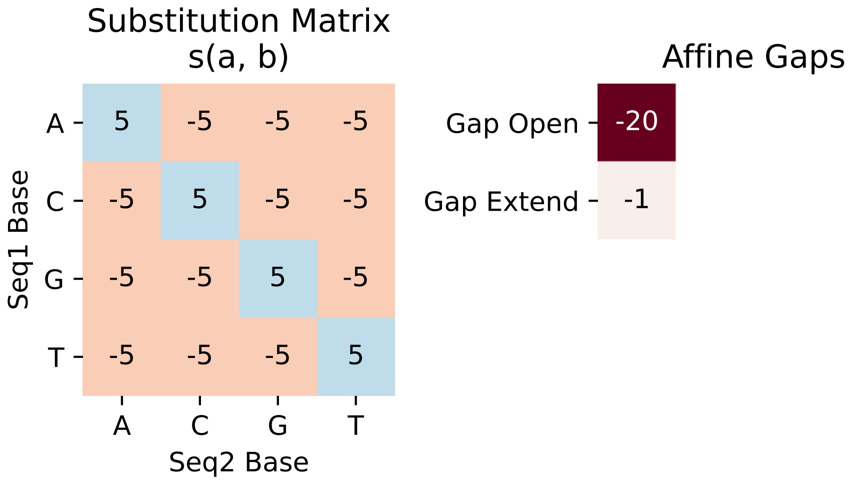

##### 1.7.3 Why affine gaps require three layers

In the linear-gap DP above, each horizontal or vertical move adds the same gap cost, so a single matrix $F(i, j)$ is enough.

With an affine penalty

$$\text{gap\_cost}(k) = g_s + (k-1)g_e,$$

we must distinguish between opening a gap (pay $g_s$) and extending an existing gap (pay $g_e$). To decide which cost applies at $(i, j)$ we need to know whether the previous step already had a gap or not, which a single value $F(i, j)$ does not record.

Gotoh's idea is to keep three scores at each cell $(i, j)$, one for each possible last move:

| State | Meaning | Last move type |
|---|---|---|
| $Y_g$ | gap in $X$ (horizontal move) | horizontal |
| $M$ | match/mismatch (diagonal) | diagonal |
| $X_g$ | gap in $Y$ (vertical move) | vertical |

This three-layer view makes the affine-gap bookkeeping local: each state only looks at the appropriate predecessors from the previous row/column.

##### 1.7.4 Gotoh recurrences

Let $\sigma(a, b)$ be the substitution score for aligning $a$ with $b$, and let $g_s$ and $g_e$ be the gap-start and gap-extend penalties.

For global alignment we first set the boundary conditions:

$$M(0,0) = 0, \qquad Y_g(0,0) = X_g(0,0) = -\infty,$$

$$Y_g(0,j) = g_s + (j-1)\,g_e \quad \text{for } j \geq 1,$$
$$X_g(i,0) = g_s + (i-1)\,g_e \quad \text{for } i \geq 1,$$

and all other entries with $i = 0$ or $j = 0$ are set to $-\infty$.

For $1 \leq i \leq n$ and $1 \leq j \leq m$ the three-layer DP updates are

$$Y_g(i,j) = \max\left\{Y_g(i,j-1) + g_e,\ M(i,j-1) + g_s,\ X_g(i,j-1) + g_s\right\},$$
$$M(i,j) = \sigma(X_i, Y_j) + \max\left\{Y_g(i-1,j-1),\ M(i-1,j-1),\ X_g(i-1,j-1)\right\},$$
$$X_g(i,j) = \max\left\{X_g(i-1,j) + g_e,\ M(i-1,j) + g_s,\ Y_g(i-1,j) + g_s\right\}.$$

Here $Y_g(i,j)$, $M(i,j)$, and $X_g(i,j)$ are exactly the "last-move" scores described above: best scores ending with a gap in $X$, a match/mismatch, or a gap in $Y$, respectively. The final global-alignment score is

$$\text{score} = \max\left\{Y_g(n,m),\ M(n,m),\ X_g(n,m)\right\}.$$

```python
# Run Gotoh affine-gap alignment
gotoh_result = nw_gotoh_matrix(
    seq1=X,
    seq2=Y,
    alphabet_to_index=alphabet_to_index,
    score_matrix=score_matrix,
    gap_open=gap_open,
    gap_extend=gap_extend,
    return_matrices=True,
)

print("Gotoh alignment score:", gotoh_result.total_score)
print()
print("Optimal alignment (affine gaps):")
print("X:", gotoh_result.seq_align1)
print("Y:", gotoh_result.seq_align2)

# Simple match / mismatch / gap pattern for the alignment:
match_line = "".join(
    "|" if a == b else " " if a == "-" or b == "-" else "x"
```

```
    for a, b in zip(gotoh_result.seq_align1, gotoh_result.seq_align2)
)
print("   " + match_line)

# Visualize the three Gotoh layers (Yg, M, Xg) with traceback path
fig_gotoh = plot_gotoh_matrices(
    gotoh_result,
    X=X,
    Y=Y,
    figsize=(10, 6),
    marker_size=26,
)

title = (
    "Gotoh affine-gap alignment\n"
    f"X: {gotoh_result.seq_align1}\n"
    f"Y: {gotoh_result.seq_align2}"
)
fig_gotoh.suptitle(title, fontsize=16, font='monospace')
fig_gotoh.tight_layout()
plt.show()
```

```
Gotoh alignment score: -6.0

Optimal alignment (affine gaps):
X: GATTACA
Y: G--TTCA
   |  |x||
```

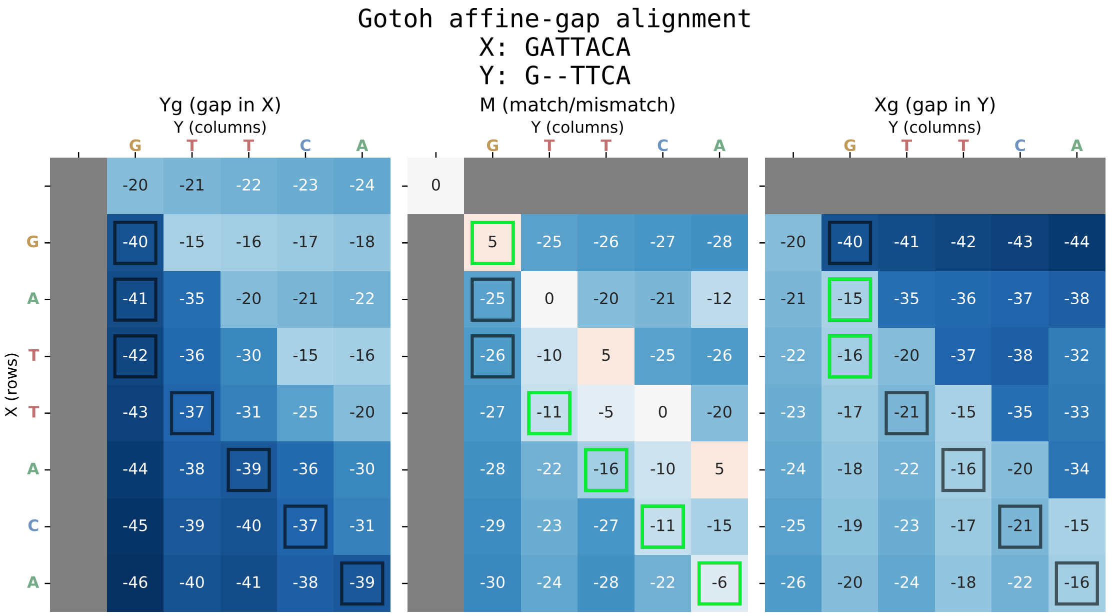

##### 1.7.5 Reading the Gotoh DP figure

The figure above shows the three dynamic-programming layers used by Gotoh's affine-gap algorithm:

- **Left panel (Yg)**: best scores for paths that end in a gap in $X$ (horizontal move).
- **Middle panel (M)**: best scores for paths that end in a match or mismatch (diagonal move).
- **Right panel (Xg)**: best scores for paths that end in a gap in $Y$ (vertical move).

Each cell shows the score for that state at position $(i, j)$, with color indicating how favorable it is (red good; blue bad). The squares trace one optimal alignment path: for each $(i, j)$ on the path we draw a faint marker on all three panels and a bright marker on the panel whose state is active there. As you follow the markers across panels, you can see when the alignment switches between "in a gap in $X$", "in a match/mismatch", and "in a gap in $Y$" to realize the affine gap penalties.

##### 1.7.6 Gotoh recurrences

Let $\sigma(a, b)$ be the substitution score for aligning $a$ with $b$, and let $g_s$ and $g_e$ be the gap-start and gap-extend penalties.

For $1 \le i \le n$ and $1 \le j \le m$ the three-layer DP updates are

$$\begin{aligned}
&\textbf{Y-layer:} && Y_g(i,j) = && && \max\Big\{Y_g(i,j-1)+g_e,\ M(i,j-1)+g_s,\ X_g(i,j-1)+g_s\Big\},\\
&\textbf{M-layer:} && M(i,j) = && \sigma(X_i,Y_j) \quad + && \max\Big\{Y_g(i-1,j-1),\ M(i-1,j-1),\ X_g(i-1,j-1)\Big\},\\
&\textbf{X-layer:} && X_g(i,j) = && && \max\Big\{X_g(i-1,j)+g_e,\ M(i-1,j)+g_s,\ Y_g(i-1,j)+g_s\Big\}.
\end{aligned}$$

Here $Y_g(i, j)$, $M(i, j)$, and $X_g(i, j)$ are exactly the "last-move" scores introduced above: best scores ending with a gap in $X$, a match/mismatch, or a gap in $Y$, respectively.

The final global-alignment score is

$$\text{score} = \max\big\{Y_g(n,m),\ M(n,m),\ X_g(n,m)\big\}.$$

For global alignment, we initialize

$$M(0,0) = 0, \qquad Y_g(0,0) = X_g(0,0) = -\infty,$$

$$Y_g(0,j) = g_s + (j-1)\,g_e \quad \text{for } j \ge 1,$$

$$X_g(i,0) = g_s + (i-1)\,g_e \quad \text{for } i \ge 1,$$

and set all other entries with $i = 0$ or $j = 0$ to $-\infty$.

##### 1.7.7 The $K_{3,3}$ gadget view

A helpful way to picture Gotoh's three-layer DP, adapted from a diagram in *Algebraic Statistics for Computational Biology* (Pachter and Sturmfels, eds.), is to treat each cell $(i, j)$ as a small $K_{3,3}$-style gadget with six ports: three incoming ($H_{in}$, $D_{in}$, $V_{in}$) and three outgoing ($H_{out}$, $D_{out}$, $V_{out}$). The internal wiring within each gadget encodes the gap-open vs gap-extend logic.

The figure below shows this structure for a small alignment grid:

- **Green** diagonal edges: match (positive substitution score)
- **Red** diagonal edges: mismatch (negative substitution score)
- **Blue** horizontal/vertical edges: gap-extend penalty $g_e$
- **Orange** internal edges: gap-initiate penalty $(g_s - g_e)$
- **Gray** internal edges: zero-cost transitions (e.g., any state → diagonal out)

```
# Draw the K, gadget DAG for the example sequences
fig_k33 = draw_affine_k33_dag(
    X,
    Y,
    match=MATCH,
    mismatch=MISMATCH,
    figsize=(8, 9),
)
```

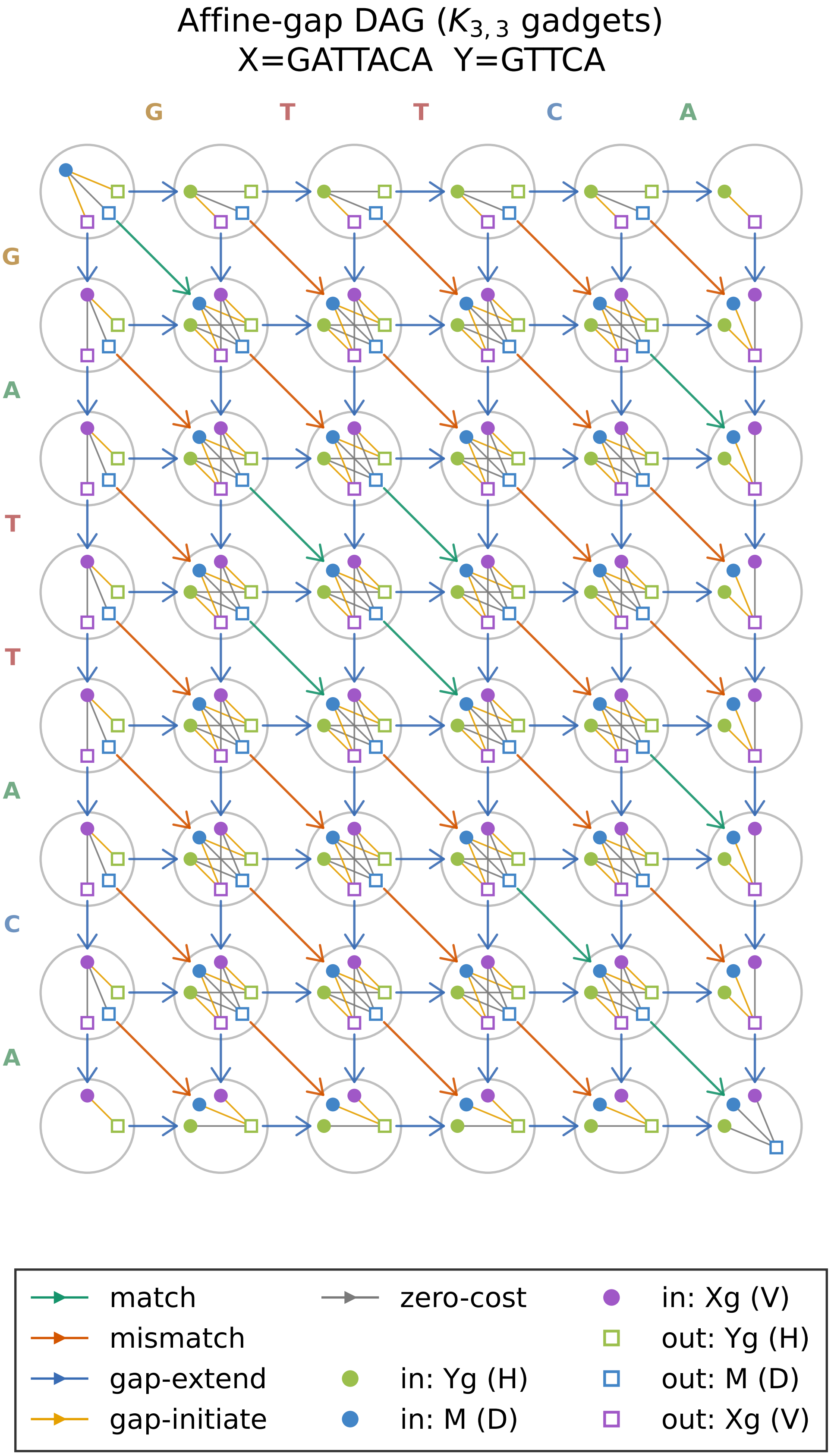
Affine-gap DAG ($K_{3,3}$ gadgets)
X=GATTACA Y=GTTCA
G
T
T
C
A
G
A
T
T
A
C
A
match
mismatch
gap-extend
gap-initiate
zero-cost
in: Yg (H)
in: M (D)
in: Xg (V)
out: Yg (H)
out: M (D)
out: Xg (V)

#### 1.8 From Gotoh to NW-flex

The NW-flex algorithm builds directly on this three-state Gotoh view:

- It keeps the same scoring scheme (match, mismatch, affine gaps).
- It augments the DP graph with **extra predecessor** (**EP**) sets $E(i)$ for selected rows $i$, which can be viewed as adding additional edges between rows in the DAG.
- This lets a designated block $Z$ "flex" (for example, an STR region), while the flanking regions $A$ and $B$ remain globally aligned.

The next notebook, *NW-flex single-block*, will use the full NW-flex core (`dp_core`, `ep_patterns`, `aligners`) together with our DP plotting helpers to show how the sets $E(i)$ modify the DP grid and how row jumps encode flexible substrings $Z^*$ in an $A \cdot Z^* \cdot B$ reference.

```
from nwflex.plot import draw_flex_dag

ans = draw_flex_dag(
        seq1= X,
        seq2= Y,
        match= 5,
        mismatch= -5,
        spacing= 1.0,
        gadget_radius= 0.35,
        inner_radius_scale= 0.65,
        max_radius_scale= 0.35,
        port_marker_size= 5.0,
        max_marker_size= 0.04,
        ep_marker_size= 4.0,
        ep_radius_scale= 1.2,
        circle_alpha= 0.5,
        circle_linewidth= 1.0,
        edge_linewidth= 1.0,
        edge_alpha= 0.9,
        lw_internal= 0.7,
        internal_alpha= 0.9,
        mutation_scale= 10.0,
        edge_shrink_factor= 0.15,
        show_legend= True,
        show_internal= True,
        figsize= (9, 10),
        title= None,
        nt_color_map= None,
    )
```

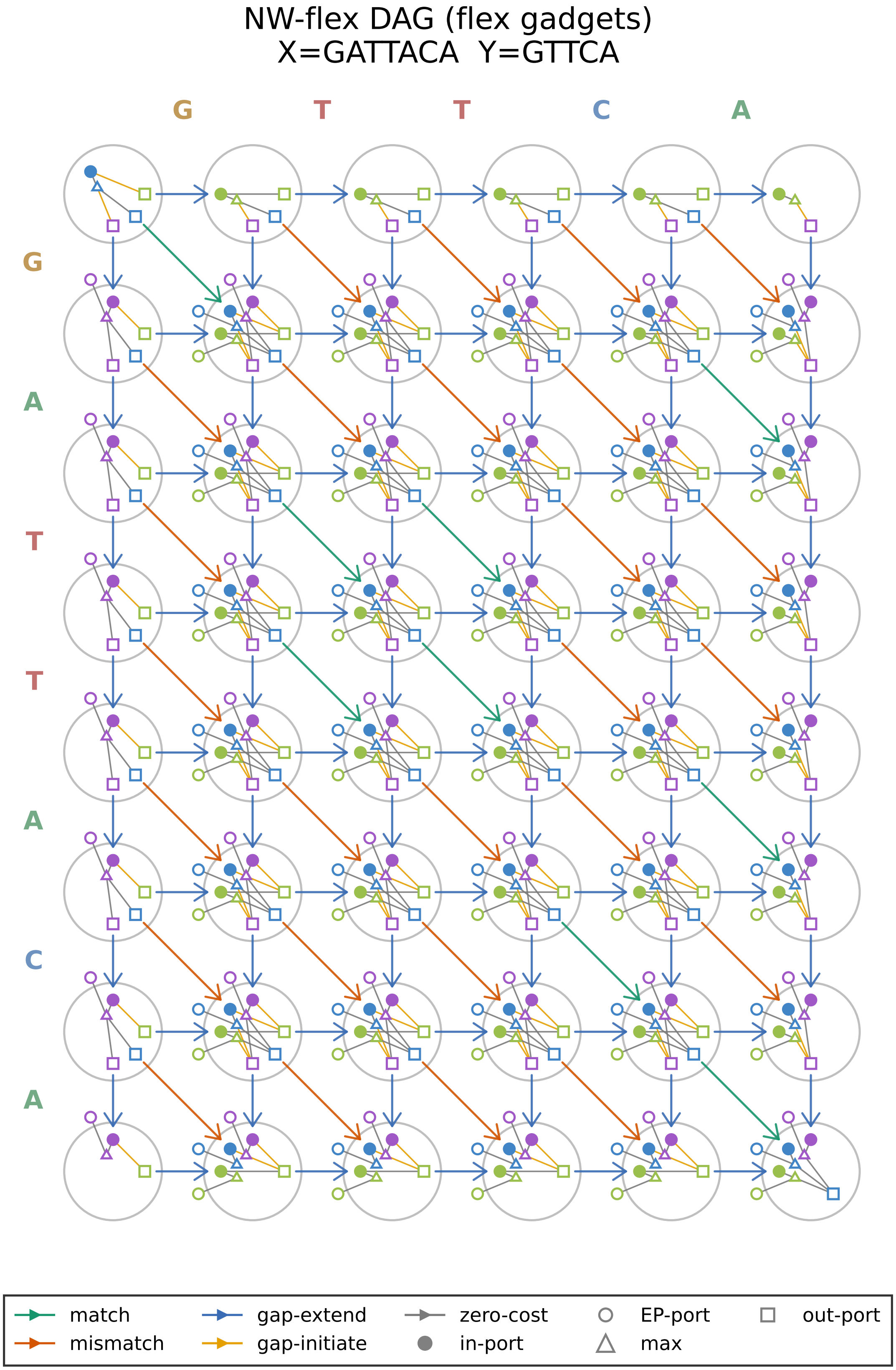
NW-flex DAG (flex gadgets)
X=GATTACA  Y=GTTCA
G
T
T
C
A
G
A
T
T
A
C
A
match
mismatch
gap-extend
gap-initiate
zero-cost
in-port
EP-port
max
out-port

### 2 Notebook 2: NW-flex for X = A·Z·B

#### 2.1 Overview

In this notebook we build directly on the basic NW-flex core:

- We define an A·Z·B locus

$$X = A \cdot Z \cdot B$$

  and a read $Y$.

- We construct three extra-predecessor patterns $E(i)$:
  - standard global NW (no extra predecessors),
  - a single flexible block on $Z$,
  - a semiglobal configuration (global in $Y$, local in $X$).
- We run the NW-flex DP core with each $E(i)$, compare scores and alignments, and visualize the DP matrices and paths.

We work directly with `run_flex_dp` so that the effect of the extra predecessors is explicit in the code.

#### 2.2 Setup and imports

```
#  Notebook magic: autoreload modules
%load_ext autoreload
%autoreload 2

import matplotlib.pyplot as plt

from nwflex.dp_core import FlexInput, run_flex_dp
from nwflex.ep_patterns import (
    build_EP_standard,
    build_EP_single_block,
    build_EP_semiglobal,
)
from nwflex.validation import get_default_scoring
from nwflex.plot import (
    plot_flex_matrices,
    plot_score_system,
    plot_ep_comparison,
    build_AZB_regions,
)
from nwflex.dp_core import AlignmentResult
```

```
The autoreload extension is already loaded. To reload it, use:
  %reload_ext autoreload
```

##### 2.3 X = A·Z·B locus and scoring

We specify the A·Z·B decomposition explicitly and then form the full reference and read:

- $A = \texttt{G}$
- $Z = \texttt{ATTA}$
- $B = \texttt{CA}$

so that

$$\begin{aligned} X = \texttt{GATTACA} &= A \cdot Z \cdot B \\ Y = \texttt{GTTCA} &= A \cdot Z^{*} \cdot B, \quad Z^{*} = \texttt{TT} \subseteq Z \end{aligned}$$

```python
# Reference X and read Y
A = "G"
Z = "ATTA"
B = "CA"

X = A + Z + B
Zstar = "TT"
Y = A + Zstar + B

print("X (reference):", X)
print("Y (read):     ", Y)

# Indices for the flexible block Z
s = len(A)
e = len(A) + len(Z)
print("Block indices: s =", s, "e =", e)
print("A =", X[:s], "Z =", X[s:e], "B =", X[e:])
```

```
X (reference): GATTACA
Y (read):      GTTCA
Block indices: s = 1 e = 5
A = G Z = ATTA B = CA
```

###### 2.3.1 Scoring

We use the default scoring scheme:

- match: $+5$
- mismatch: $-5$
- gap start: $-20$
- gap extend: $-1$

```python
# Scoring scheme
score_matrix, gap_open, gap_extend, alphabet_to_index = get_default_scoring()

# Display the default score scheme:
clean_map = {str(k): v for k, v in alphabet_to_index.items()}
print("Score matrix:\n", score_matrix.astype(int))
print("gap_open:", gap_open)
print("gap_extend:", gap_extend)
print("alphabet_to_index:", clean_map)
fig = plot_score_system(score_matrix, gap_open, gap_extend, alphabet_to_index)
plt.show()
```

```
Score matrix:
 [[ 5 -5 -5 -5]
 [-5  5 -5 -5]
 [-5 -5  5 -5]
 [-5 -5 -5  5]]
gap_open: -20.0
gap_extend: -1.0
alphabet_to_index: {'A': 0, 'C': 1, 'G': 2, 'T': 3}
```

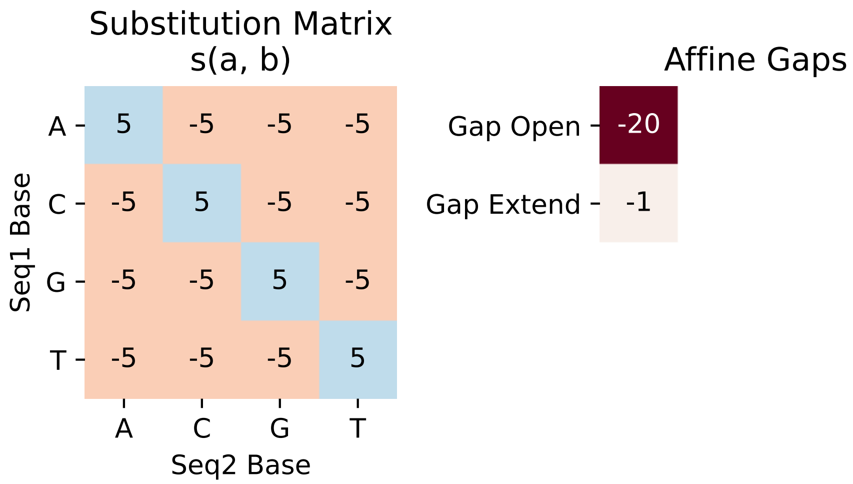

```python
n = len(X)

# Build the three EP patterns
EP_standard = build_EP_standard(n)
EP_block = build_EP_single_block(n, s, e)
EP_semiglobal = build_EP_semiglobal(n)

# For plotting, define leaders for each pattern:
# - Standard NW: no leaders (no EP edges)
```

```python
# - Single-block: leader row is s (edges from s skip into Z)
# - Semiglobal: leader row is 0 (edges from source skip into X)
leaders_list = [
    [],    # Standard NW: no leaders
    [s],   # Single-block: s is the leader
    [0],   # Semiglobal: 0 is the leader
]

# Define background regions (A·Z·B decomposition)
# Colors: A=blue, Z=orange, B=purple, X=grey
color_A, color_Z, color_B, color_X = "#d0e0ff", "#ffe0d0", "#e0d0f0", "#e0e0e0"
label_A, label_Z, label_B, label_X = "#2060a0", "#c06020", "#7030a0", "#404040"

bg_standard = [
    {'start': 1, 'end': n, 'color': color_X,
     'label': 'X', 'label_color': label_X},
]
bg_block = [
    {'start': 1, 'end': s, 'color': color_A,
     'label': 'A', 'label_color': label_A},
    {'start': s + 1, 'end': e, 'color': color_Z,
     'label': 'Z', 'label_color': label_Z},
    {'start': e + 1, 'end': n, 'color': color_B,
     'label': 'B', 'label_color': label_B},
]
bg_semiglobal = [
    {'start': 1, 'end': n, 'color': color_Z,
     'label': 'local\nin X', 'label_color': label_Z},
]

backgrounds = [bg_standard, bg_block, bg_semiglobal]

# Row annotations (for labeling special rows)
# For single-block: leader is s, closer is e+1
# For semiglobal: leader is 0 (s=0), closer is n+1 (e+1 where e=n)
annot_standard = {}
annot_block = {s: "(s)", e + 1: "(e+1)"}
annot_semiglobal = {0: "(s)", n + 1: "(e+1)"}

row_annotations = [annot_standard, annot_block, annot_semiglobal]

# Titles
titles = [
    "Standard NW\n$E(i) = \\varnothing$",
    "Single-block flex\non Z",
    "Semiglobal\n(local in X)",
]
```

```
# Plot with styling matched to the DAG plot
# Boxes are sized to match DAG: 2 columns wide, 1 row tall
# row_spacing > node_height creates gaps for standard edges
fig = plot_ep_comparison(
    EP_list=[EP_standard, EP_block, EP_semiglobal],
    leaders_list=leaders_list,
    titles=titles,
    X=X,
    backgrounds=backgrounds,
    row_annotations_list=row_annotations,
    figsize=(12, 14),
    wspace=0.05,
    # Match DAG plot styling
    closer_edge_color="#7030a0",
    leader_curvature=0.3,
    closer_curvature=0.25,
    ep_linewidth=3.0,
    sequence_fontsize=18.0,  # Match DAG
    sequence_fontweight="semibold",
    region_fontsize=28.0,
    row_label_fontsize=12.0,
    row_label_fontweight="normal",  # Not bold
    legend_fontsize=16.0,
    # Box sizing and spacing
    node_width=2.0,      # 2 DAG columns wide
    node_height=0.7,     # Slightly shorter than row_spacing
    row_spacing=1.2,     # More space between rows for standard edges
    box_label_fontsize=14.
)
plt.show()
```

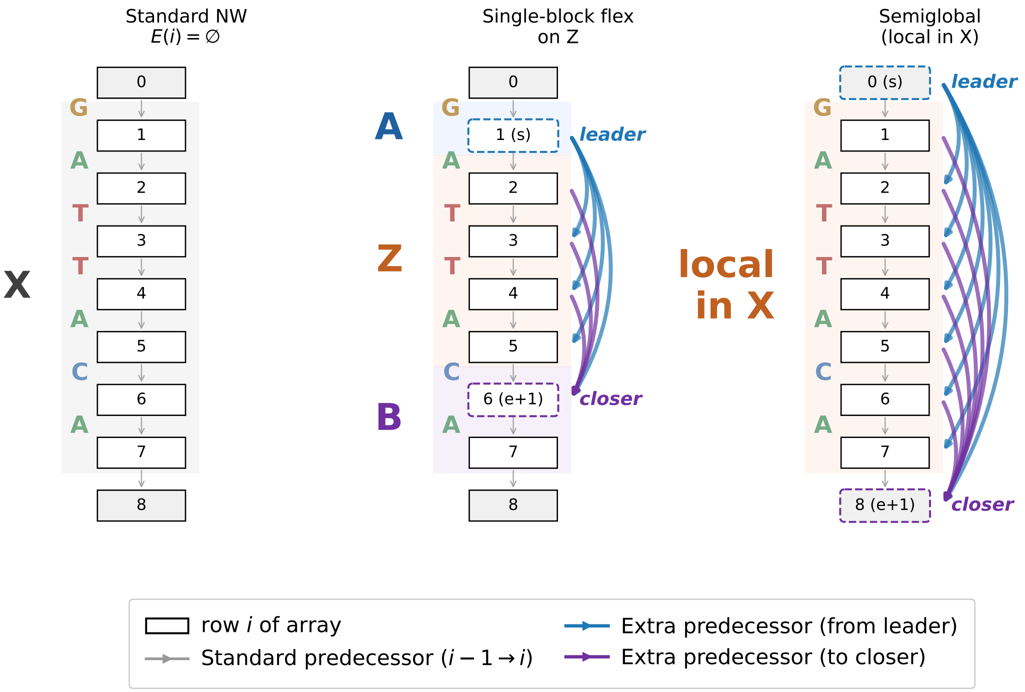

##### 2.3.2 Interpreting the EP patterns

The three panels above illustrate how extra predecessors modify the DP graph:

- **Standard NW** (left): No EP edges. Every row $i$ has only its standard predecessor $i - 1$. The DP proceeds row by row with no shortcuts.
- **Single-block flex** (center): Row $s = 2$ is the leader. Blue edges from $s$ reach into the flexible region $Z$, allowing the DP to skip the beginning of $Z$. Purple edges from rows within $Z$ converge at the closer row $e + 1 = 9$, allowing the DP to exit $Z$ early. Together, these edges let NW-flex align any contiguous substring $Z^* \subseteq Z$.
- **Semiglobal** (right): Row 0 is the leader for the entire sequence. Blue edges from the source reach every row, allowing alignment to start anywhere in $X$. Purple edges from all rows converge at the final row, allowing alignment to end anywhere. This realizes “local in $X$, global in $Y$” alignment.

```
# Single-panel DAG with EP arcs on the right
from nwflex.plot import plot_dag_with_ep_arcs

fig_dag_ep = plot_dag_with_ep_arcs(
    EP=EP_block,
    leaders=[s],
    X=X,
    Y=Y,
    s=s,
```

```
    e=e,
    node_radius = 0.15,
    figsize=(10, 10),
    color_A=color_A,
    color_Z=color_Z,
    color_B=color_B,
    label_A=label_A,
    label_Z=label_Z,
    label_B=label_B,
    closer_edge_color="#7030a0",
    leader_curvature=0.3,
    closer_curvature=0.25,
    title="Alignment DAG with EP edges",
    dag_linewidth=2.0,
    ep_linewidth=3.0,
    sequence_fontweight="semibold",
    region_fontsize=28.0,
    box_label_fontsize=16.0,
    legend_fontsize=16.0,
    legend_bbox_to_anchor=(0.45, -0.18)

)

plt.show()
```

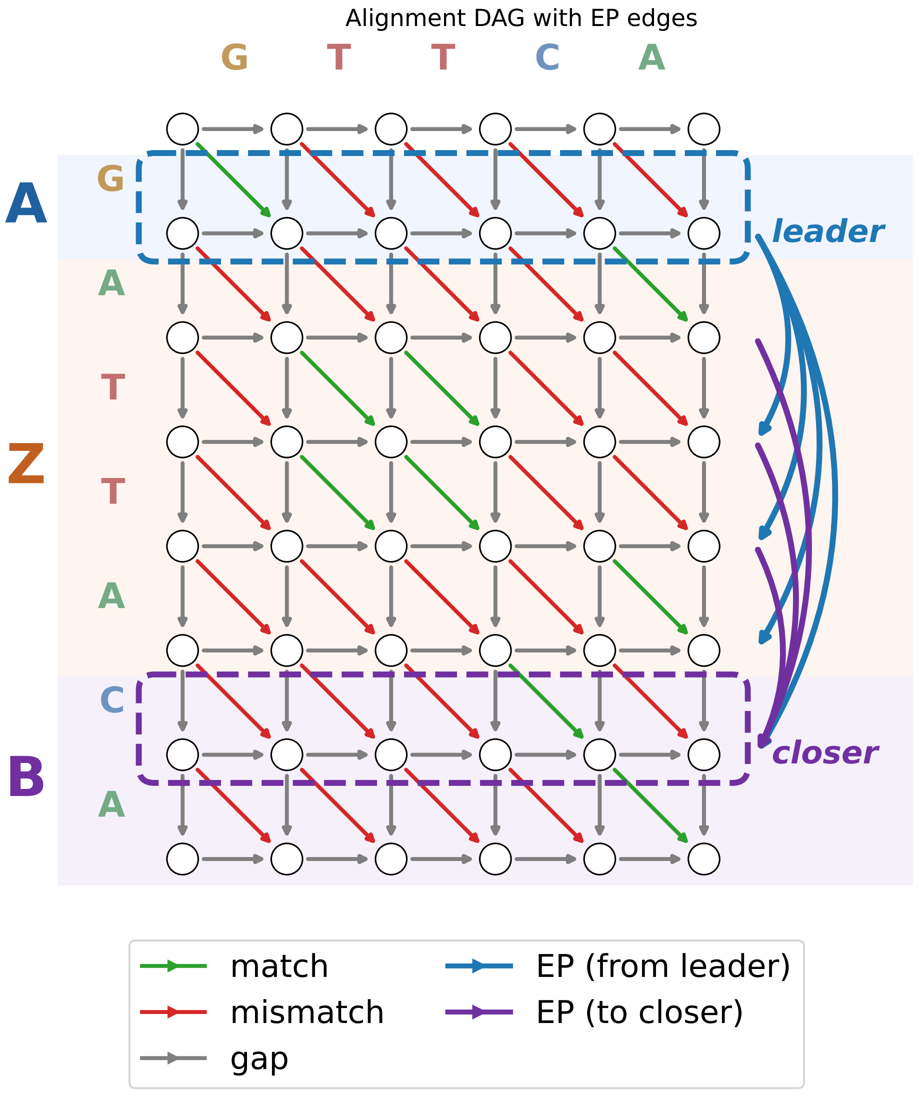

##### 2.3.3 The alignment DAG with EP edges

The figure above shows the alignment DAG for $X = A \cdot Z \cdot B$ and $Y$, augmented with extra predecessor (EP) edges on the right margin:

- **Standard DAG edges** (interior): Gray horizontal/vertical edges represent gaps; green/red diagonals represent matches/mismatches. These connect adjacent cells $(i, j) \to (i + 1, j + 1)$, $(i, j) \to (i + 1, j)$, or $(i, j) \to (i, j + 1)$.
- **EP edges** (right margin): Curved arcs connecting non-adjacent rows.
  - **Leader edges** (blue): From row $s$ into the flexible block $Z$. These allow the DP to "jump into" $Z$ from the leader row.
  - **Closer edges** (purple): From rows within $Z$ to the closer row $e + 1$. These allow the DP to "jump out" of $Z$ to resume global alignment in $B$.

- **Region shading**: The A·Z·B decomposition is shown as horizontal bands — blue for $A$, orange for $Z$, purple for $B$.

The EP edges act on **entire rows**: when the DP uses a leader edge from row $s$ to row $i > s+1$, it skips the intervening characters of $X$ without paying gap penalties. This is what enables NW-flex to find the best contiguous substring $Z^* \subseteq Z$ in a single pass.

##### 2.4 Running the NW-flex core

We wrap `run_flex_dp` in a small helper that takes $(X, Y, E)$ and returns the score, aligned sequences, the DP path, the row jumps, and the full DP state.

We then run:

- standard NW (EP_standard),
- single-block NW-flex (EP_block),
- semiglobal NW-flex (EP_semiglobal),

and compare their scores and alignments.

```
def run_with_EP(X, Y, EP):
    """Run NW-flex DP with a given EP pattern and return the result."""
    config = FlexInput(
        X=X,
        Y=Y,
        score_matrix=score_matrix,
        gap_open=gap_open,
        gap_extend=gap_extend,
        extra_predecessors=EP,
        alphabet_to_index=alphabet_to_index,
    )
    res = run_flex_dp(
        config,
        return_data=True,
    )
    return config, res

# Run alignment with each EP pattern
cfg_std, res_std = run_with_EP(X, Y, EP_standard)
cfg_blk, res_blk = run_with_EP(X, Y, EP_block)
cfg_sg,  res_sg  = run_with_EP(X, Y, EP_semiglobal)

print("Standard NW score:", res_std.score)
print(res_std.X_aln)
print(res_std.Y_aln)
print()

print("Single-block NW-flex score:", res_blk.score, sep="\n")
print(res_blk.X_aln)
```

```
print(res_blk.Y_aln)
print("Row jumps:", *res_blk.jumps, sep="\n")
print()

print("Semiglobal NW-flex score:", res_sg.score, sep="\n")
print(res_sg.X_aln)
print(res_sg.Y_aln)
print("Row jumps:", *res_sg.jumps, sep="\n")
```

```
Standard NW score: -6.0
GATTACA
G--TTCA

Single-block NW-flex score:
25.0
GTTCA
GTTCA
Row jumps:
RowJump(from_row=1, to_row=3, col=2, state=1)
RowJump(from_row=4, to_row=6, col=4, state=1)

Semiglobal NW-flex score:
5.0
GATTA
GTTCA
Row jumps:
RowJump(from_row=8, to_row=5, col=5, state=1)
```

In this example:

- Standard NW must pay for deleting part of $Z$ with gaps.
- The single-block EP pattern lets the DP jump over a substring of $Z$ and treat it as unaligned, effectively aligning $A \cdot Z^* \cdot B$.
- The semiglobal EP pattern is more permissive at the ends: the entire $X$ can flex, which is appropriate when we want to align the whole read $Y$ to any part of the reference sequence $X$.

##### 2.5 Visualizing the DP matrices and paths

We now visualize the $Y_g$, $M$, and $X_g$ score layers for each configuration using `plot_flex_matrices`, highlighting:

- the flexible block rows (for the $ A·Z·B $ case), and
- the best DP path and row jumps that arise from extra predecessors.
- the start and end positions of the optimal internal substring $Z^*$.

```
fig_size = (10, 5)
ms = 30
GOLD = '#DAA520'  # Gold color matching jump edges
```

```python
# Standard NW (no region boxes - standard NW doesn't use A·Z·B decomposition)
# (no start and end rows (start before -1 and end after n))
fig_std = plot_flex_matrices(
    result=res_std,
    X=X,
    Y=Y,
    s=-2,
    e=n+1,
    marker_size=ms,
    figsize=fig_size,
)
fig_std.suptitle("Standard NW (E(i) = empty)")
fig_std.tight_layout(w_pad=2.0)
plt.show()

# Single-block NW-flex with A·Z·B region boxes
# Get Z* boundaries from alignment jumps
# jumps[0] is entry jump (from A into Z*), jumps[1] is exit jump (from Z* to B)
entry_jump = res_blk.jumps[0]   # Jump from row s (last of A) into Z*
exit_jump = res_blk.jumps[1]    # Jump from Z* to row e+1 (first of B)
zstar_start = entry_jump.to_row    # First row of Z* (where we land after entry)
zstar_end = exit_jump.from_row     # Last row of Z* (where we jump from)

regions_block = build_AZB_regions(s=s, e=e, n=len(X))
# Add Z* as a dotted gold box inside Z, with label at top
regions_block.append({
    'start': zstar_start, 'end': zstar_end,
    'label': 'Z*', 'color': GOLD,
    'linestyle': ':', 'label_position': 'top',
})
fig_blk = plot_flex_matrices(
    result=res_blk,
    X=X,
    Y=Y,
    s=s,
    e=e,
    marker_size=ms,
    figsize=fig_size,
    regions=regions_block,
)
fig_blk.suptitle("Single-block NW-flex on Z")
fig_blk.tight_layout(w_pad=2.0)
plt.show()

# Semiglobal NW-flex visualization:
# here the flexible block spans all of X (s=0, e=n), so only X region shown
# Get X* boundaries from alignment jumps
```

```
#
# For semiglobal, we need to check for two types of jumps:
# 1. Terminal jump: from_row = n+1 (virtual) -> to_row = actual end row (xstar_end)
# 2. Entry jump: from_row = 0 (leader) -> to_row = actual start row (xstar_start)
#
# If there's no entry jump from row 0, alignment starts at row 1.
# If there's no terminal jump to a row < n, alignment ends at row n.

n_sg = len(X)

# Find terminal jump (from virtual row n+1) to determine xstar_end
terminal_jump = None
for jmp in res_sg.jumps:
    if jmp.from_row == n_sg + 1:
        terminal_jump = jmp
        break

# Find entry jump (from leader row 0) to determine xstar_start
entry_jump_sg = None
for jmp in res_sg.jumps:
    if jmp.from_row == 0 and jmp.to_row > 1:
        entry_jump_sg = jmp
        break

# Determine X* boundaries
xstar_start = entry_jump_sg.to_row if entry_jump_sg is not None else 1
xstar_end = terminal_jump.to_row if terminal_jump is not None else n_sg

regions_semiglobal = [
    {'start': 1, 'end': len(X), 'label': 'X', 'color': '#d04000'},
    {'start': xstar_start, 'end': xstar_end, 'label': 'X*',
     'color': GOLD, 'linestyle': ':', 'label_position': 'top'},
]

fig_sg = plot_flex_matrices(
    result=res_sg,
    X=X,
    Y=Y,
    s=0,
    e=len(X),
    marker_size=ms,
    figsize=fig_size,
    regions=regions_semiglobal,
)
fig_sg.suptitle("Semiglobal-like NW-flex (global in Y, local in X)")
fig_sg.tight_layout(w_pad=2.0)
```

```
plt.show()
```

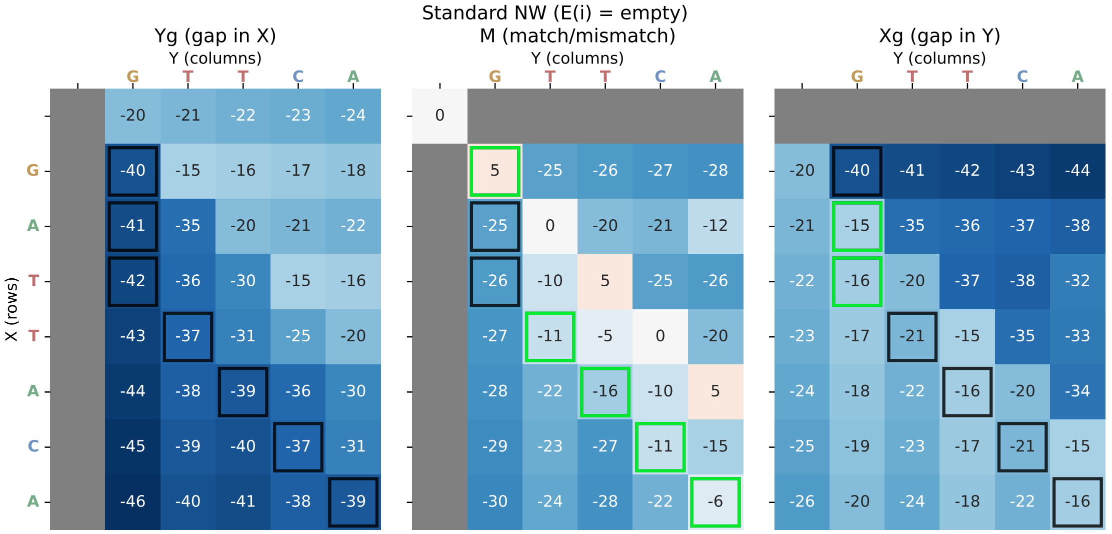
Standard NW (E(i) = empty)
Yg (gap in X)
M (match/mismatch)
Xg (gap in Y)
Y (columns)
X (rows)

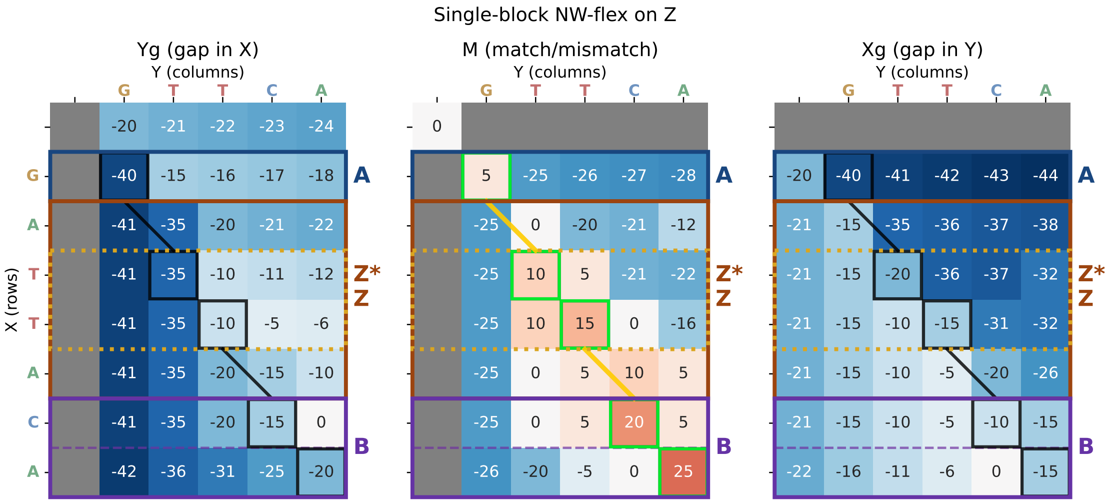
Single-block NW-flex on Z
Yg (gap in X)
M (match/mismatch)
Xg (gap in Y)
Y (columns)
X (rows)
A
Z*
Z
B

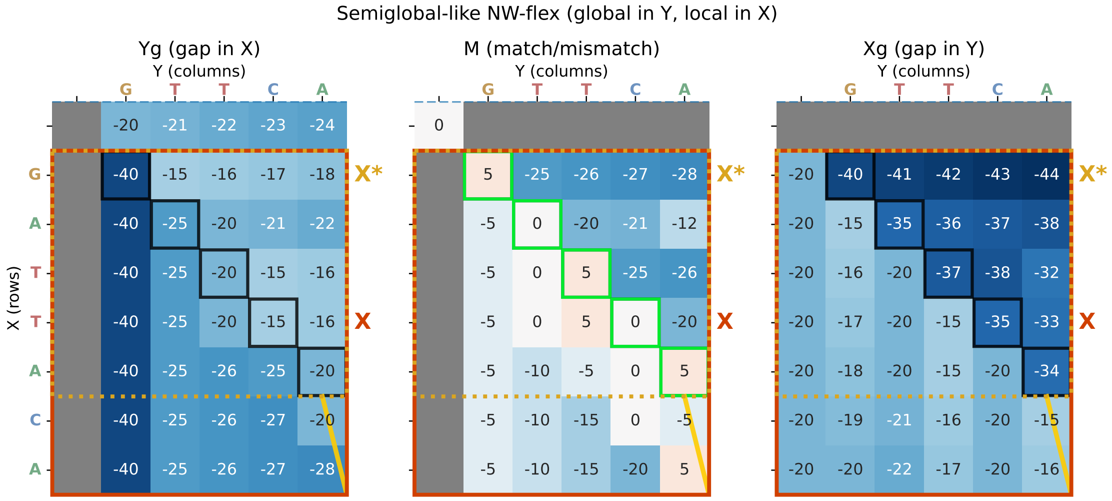

##### 2.5.1 Interpreting the DP matrix visualizations

The three figures above show the Gotoh DP matrices ($Y_g$, $M$, $X_g$) for each alignment mode, with the optimal path traced in black and green. The square is green if the corresponding cell is part of the optimal path. Row jumps are shown as gold diagonal lines. Dashed horizontal lines mark the **leader row** (blue) and **closer row** (purple) that bound the flexible region.

**Standard NW** (top): The alignment proceeds row by row without shortcuts. The path must traverse all rows of $X$, paying gap penalties to accommodate the length difference between $Z$ and $Z^*$. No region boxes are shown because standard NW treats $X$ uniformly.

**Single-block NW-flex** (middle): The $A \cdot Z \cdot B$ decomposition is highlighted with colored boxes: - **A** (blue): The prefix flank, aligned globally. - **Z** (orange, solid): The flexible block where the DP can skip rows. - **Z*** (gold, dotted): The actual substring of $Z$ that was aligned — the optimal $Z^* \subseteq Z$. The gold color matches the jump edges. - **B** (purple): The suffix flank, aligned globally.

The dashed blue line marks row $s$ (the leader), and the dashed purple line marks row $e + 1$ (the closer). The gold diagonal lines show where the path "jumps" — entering $Z^*$ from $A$ and exiting to $B$ — skipping the unaligned portions of $Z$ without gap penalties.

**Semiglobal NW-flex** (bottom): The entire sequence $X$ is treated as a flexible block: - **X** (orange, solid): The full reference, all of which can flex. - **X*** (gold, dotted): The actual aligned portion of $X$.

This realizes "local in $X$, global in $Y$" alignment: the read $Y$ is fully aligned, but only the best-matching substring of $X$ participates.

#### 2.6 Semi-global alignment with `free_X` flag

The semi-global alignment ("local in X, global in Y") can be executed using the `free_X=True` flag in `FlexInput`, which:

1. Initializes `Xg[i, 0] = 0` for all rows (free leading gaps in X)

2. Allows traceback to start at any row in the final column

This replaces the need for the EP-based semiglobal pattern for this use case. We verify correctness with a test case where $Y$ is a short substring of $X$.

```
# Show example of semi-global alignment with free_X flag

X_sg = "TCAGAGATTACACGTAC"
Y_sg = "GATTACA"

# Build a standard EP pattern (no extra predecessors needed with free_X)
EP_sg_test = build_EP_standard(len(X_sg))

# Run with free_X=True (semi-global: local in X, global in Y)
config_sg_free = FlexInput(
    X=X_sg,
    Y=Y_sg,
    score_matrix=score_matrix,
    gap_open=gap_open,
    gap_extend=gap_extend,
    extra_predecessors=EP_sg_test,
    alphabet_to_index=alphabet_to_index,
    free_X=True,  # <-- semi-global: local in X
    free_Y=False, # global in Y
)

score_sg_free, X_aln_sg, Y_aln_sg, path_sg, jumps_sg, data_sg = run_flex_dp(
    config_sg_free,
    return_data=True,
).to_tuple()

print("Semi-global (free_X=True) alignment:")
print(f"  X = {X_sg}")
print(f"  Y = {Y_sg}")
print(f"  Score: {score_sg_free} (expected: {len(Y_sg)*5})")
print(f"  X_aln: {X_aln_sg}")
print(f"  Y_aln: {Y_aln_sg}")
print()

# Compare with standard global alignment
config_sg_global = FlexInput(
    X=X_sg,
    Y=Y_sg,
    score_matrix=score_matrix,
    gap_open=gap_open,
    gap_extend=gap_extend,
    extra_predecessors=EP_sg_test,
    alphabet_to_index=alphabet_to_index,
```

```
    free_X=False,  # standard global
)

score_sg_global, X_aln_global, Y_aln_global, _, _, _ = run_flex_dp(
    config_sg_global,
    return_data=True,
).to_tuple()

print("Standard global alignment:")
print(f"  Score: {score_sg_global}")
print(f"  X_aln: {X_aln_global}")
print(f"  Y_aln: {Y_aln_global}")
print()

# Verify
assert score_sg_free == len(Y_sg)*5, f"Expected score {len(Y_sg)*5}, got␣
 ↪{score_sg_free}"
print(" Semi-global alignment test passed!")
```

```
Semi-global (free_X=True) alignment:
  X = TCAGAGATTACACGTAC
  Y = GATTACA
  Score: 35.0 (expected: 35)
  X_aln: TCAGAGATTACACGTAC
  Y_aln: -----GATTACA-----

Standard global alignment:
  Score: -13.0
  X_aln: TCAGAGATTACACGTAC
  Y_aln: -----GATTACA-----

 Semi-global alignment test passed!
```

```
# Visualize the DP matrices for semi-global alignment
# Wrap result for plotting
res_sg_free = AlignmentResult(
    score=score_sg_free,
    X_aln=X_aln_sg,
    Y_aln=Y_aln_sg,
    jumps=jumps_sg,
    data=data_sg,
    path=path_sg,
)

# Determine X* boundaries from the aligned sequences
# Find which rows of X were actually used (non-gap characters in X_aln)
# The alignment tells us which substring of X was matched
```

```python
x_pos = 0  # position in X
xstar_positions = []
for x_char, y_char in zip(X_aln_sg, Y_aln_sg):
    if x_char != '-':
        x_pos += 1
        if y_char != '-':
            xstar_positions.append(x_pos)

xstar_start_sg = min(xstar_positions)
xstar_end_sg = max(xstar_positions)

regions_sg_free = [
    {'start': 1, 'end': len(X_sg), 'label': 'X', 'color': '#d04000'},
    {'start': xstar_start_sg, 'end': xstar_end_sg, 'label': 'X*',
     'color': GOLD, 'linestyle': ':', 'label_position': 'top'},
]

fig_sg_free = plot_flex_matrices(
    result=res_sg_free,
    X=X_sg,
    Y=Y_sg,
    s=0,
    e=len(X_sg),
    marker_size=14,
    figsize=fig_size,
    regions=regions_sg_free,
)
fig_sg_free.suptitle("Semi-global with free_X=True (local in X, global in Y)")
fig_sg_free.tight_layout(w_pad=2.0)
plt.show()
```

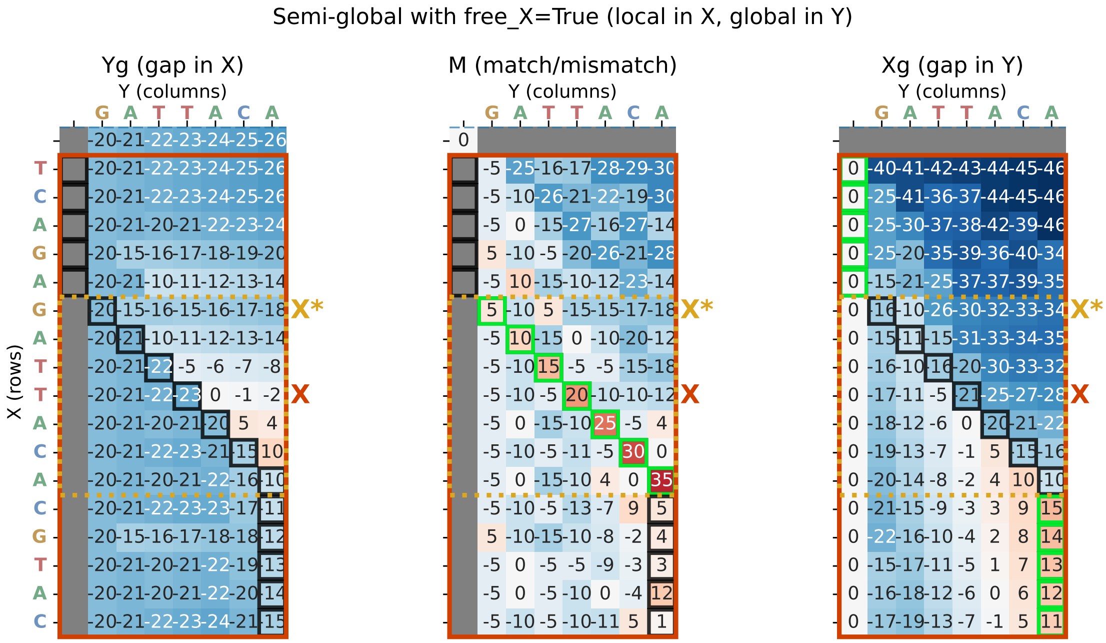

#### 2.7 Semi-global alignment with `free_Y` flag

The `free_Y=True` flag performs semi-global alignment that is "global in X, local in Y". This:

1. Initializes `Yg[0, j] = 0` for all columns (free leading gaps in Y)
2. Allows traceback to start at any column in the final row

This is useful when the read $Y$ extends beyond the reference $X$. For example, when a read partially overlaps the reference at either end. The unaligned portion of $Y$ (the overhang) incurs no penalty.

Note: Unlike `free_X`, the `free_Y` mode cannot be achieved with EP patterns alone, since EP operates on rows (reference positions), not columns (read positions).

```python
# Example of semi-global alignment with free_Y and free_X flag

X_sg = "GATTACA"
Y_sg = "ACACC"

# Build a standard EP pattern (no extra predecessors needed with free_Y)
EP_sg_test = build_EP_standard(len(X_sg))

# Run with free_Y=True (semi-global: global in X, local in Y)
config_sg_free = FlexInput(
    X=X_sg,
    Y=Y_sg,
    score_matrix=score_matrix,
    gap_open=gap_open,
```

```
    gap_extend=gap_extend,
    extra_predecessors=EP_sg_test,
    alphabet_to_index=alphabet_to_index,
    free_X=True,
    free_Y=True,  # <-- semi-global: local in Y
)

score_sg_free, X_aln_sg, Y_aln_sg, path_sg, jumps_sg, data_sg = run_flex_dp(
    config_sg_free,
    return_data=True,
).to_tuple()

print("Semi-global (free_Y=True) alignment:")
print(f"  X = {X_sg}")
print(f"  Y = {Y_sg}")
print(f"  Score: {score_sg_free}")
print(f"  X_aln: {X_aln_sg}")
print(f"  Y_aln: {Y_aln_sg}")
print()

# Compare with standard global alignment
config_sg_global = FlexInput(
    X=X_sg,
    Y=Y_sg,
    score_matrix=score_matrix,
    gap_open=gap_open,
    gap_extend=gap_extend,
    extra_predecessors=EP_sg_test,
    alphabet_to_index=alphabet_to_index,
    free_Y=False,  # standard global
)

score_sg_global, X_aln_global, Y_aln_global, _, _, _ = run_flex_dp(
    config_sg_global,
    return_data=True,
).to_tuple()

print("Standard global alignment:")
print(f"  Score: {score_sg_global}")
print(f"  X_aln: {X_aln_global}")
print(f"  Y_aln: {Y_aln_global}")
print()
```

```
Semi-global (free_Y=True) alignment:
  X = GATTACA
  Y = ACACC
  Score: 15.0
  X_aln: GATTACA--
```

```
  Y_aln: ----ACACC

Standard global alignment:
  Score: -26.0
  X_aln: GATTACA
  Y_aln: --ACACC
```

##### 2.8 Summary

- Extra predecessor sets $E(i)$ let us express different alignment modes (standard, \$ A $\cdot Z \cdot B\$flex, semiglobal)inaunifiedDPframework.Thesingle-blockEPpatternrealizesS_{\text{flex}}(X, Y) = \max_{Z^* \subseteq Z}$ NWG$(A \cdot Z^* \cdot B, Y)$ in a single pass.
- The semiglobal EP pattern uses a full-length flexible block so that the entire reference can flex while the read remains globally aligned.

  The `free_X` flag in `FlexInput` provide a simpler way to achieve semi-global alignment without EP edges:

  - `free_X=True`: local in X, global in Y (find best substring of X matching Y)
  - `free_Y=True`: global in X, local in Y (align all of Y to any part of X)

  In the next notebook we confirm that the NW-flex satisfies the guarantee of identifying an optimal alignment over all substrings.

#### 3 Notebook 3: Validating NW-flex optimality

##### 3.1 Overview

In this notebook we confirm that the single-block NW-flex configuration returns the expected **flex optimum** defined as

$$S_{\text{flex}}(X, Y) = \max_{Z^* \subseteq Z} \mathrm{NWG}(A \cdot Z^* \cdot B,\ Y)$$

for references of the form $X = A \cdot Z \cdot B$, where $Z^*$ ranges over contiguous substrings of $Z$.

This notebook builds on:

- **Notebook 1**: the Needleman–Wunsch/Gotoh recurrences and DP basics,
- **Notebook 2**: the NW-flex core with extra-predecessor (EP) patterns.

We will:

1. Verify that NW-flex with $E(i) = \varnothing$ (no extra predecessors) matches an independent NWG implementation.
2. Show the EP pattern for a single flexible block $Z$.
3. Enumerate all substrings $Z^* \subseteq Z$ on an explicit example and compare the naive maximum to the NW-flex score.
4. Run manual and randomized tests to confirm score equality.
5. Compare alignment strings (not just scores) to verify correctness.
6. Visualize the DP matrices and row jumps for the explicit example.

##### 3.2 Setup and imports

We use:

- validation helpers from `validation.py`: `nwg_global`, `check_standard_case`, `check_AZB_case`, `check_alignment_validity`, and `run_mutated_AZB_tests`,
- the single-block aligner `align_single_block` from `aligners.py`,
- the EP pattern builder `build_EP_single_block` from `ep_patterns.py`,
- and plotting helpers from `nwflex.plot`: `plot_score_system`, `plot_ep_pattern`, `draw_ep_background`, and `plot_flex_matrices_publication`.

All calls use the default scoring scheme: - match: $+5$, mismatch: $-5$ - gap open: $-20$ - gap extend: $-1$

Simulation parameters can be adjusted in the "Global test parameters" cell below.

```
#  Notebook magic: autoreload modules
%load_ext autoreload
%autoreload 2

import random
import numpy as np
import pandas as pd
import matplotlib.pyplot as plt
import matplotlib.gridspec as gridspec

from nwflex.validation import (
    get_default_scoring,
    nwg_global,
    check_standard_case,
    check_AZB_case,
    check_alignment_validity,
    run_mutated_AZB_tests,
)
from nwflex.aligners import align_single_block
from nwflex.ep_patterns import build_EP_single_block
from nwflex.plot import (
    plot_score_system,
    plot_ep_pattern,
    draw_ep_background,
    plot_flex_matrices_publication,
    REGION_FILL_COLORS,
    REGION_LABEL_COLORS,
)

from nwflex.plot.colors import HEATMAP_COLORMAPS
grid_color_map = HEATMAP_COLORMAPS['diverging']

# ==============================================================================
```

```
# Global test parameters - adjust these to control validation thoroughness
# ==============================================================================
NUM_MUTATION_VARIANTS = 100    # Number of mutated Y variants per base case
NUM_RANDOM_CASES = 100         # Number of random A·Z·B tests to run
NUM_SCORING_TESTS = 100        # Number of tests per scoring scheme variant

# Load default scoring
score_matrix, gap_open, gap_extend, alphabet_to_index = get_default_scoring()

# Show the scoring system
fig = plot_score_system(score_matrix, gap_open, gap_extend, alphabet_to_index)
plt.show()
```

```
The autoreload extension is already loaded. To reload it, use:
  %reload_ext autoreload
```

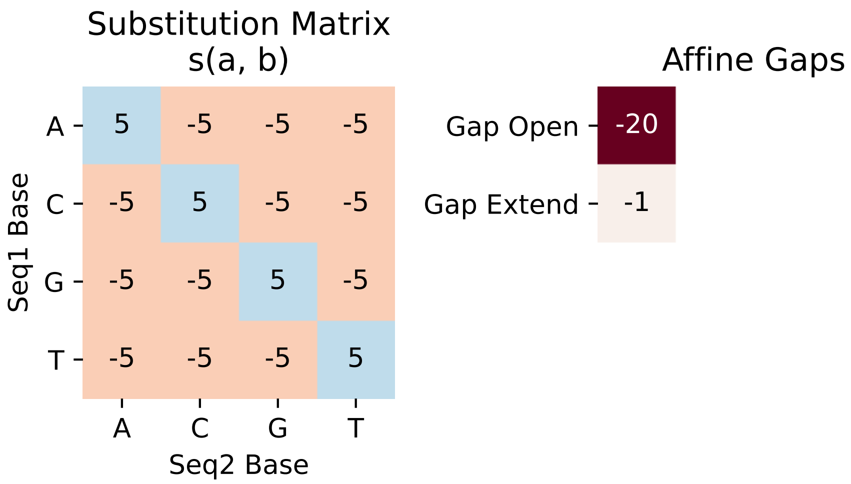

##### 3.3 Base case: NW-flex with no extra predecessors

As a sanity check, we first verify that NW-flex with $E(i) = \varnothing$ for all rows (via `align_standard`) gives the same score as the independent Needleman–Wunsch/Gotoh implementation `nwg_global`. The helper `check_standard_case` runs both and returns their scores for comparison.

This confirms that our DP core is correct before we add extra predecessors.

```
# Test cases covering different alignment scenarios:
# - Identical sequences (perfect match)
# - Single deletion (one base missing in Y)
# - Substitution (one mismatch)
# - Multiple gaps (contraction/expansion)
# - Mixed operations (substitution + indel)
```

```python
examples = [
    ("GATTACA", "GATTACA", "identical"),
    ("GATTACA", "GATACA", "single deletion"),
    ("ACGT", "ACCT", "single substitution"),
    ("GGGATATGGG", "GGGATGGG", "internal deletion"),
    ("AACGT", "ACGTT", "deletion + insertion"),
]

results = []
for X, Y, desc in examples:
    flex_score, nwg_score = check_standard_case(X, Y)
    match = "[OK]" if flex_score == nwg_score else "[FAIL]"
    results.append({
        "case": desc, "X": X, "Y": Y,
        "flex": int(flex_score), "nwg": int(nwg_score), "match": match
    })
    assert flex_score == nwg_score, "Standard NW-flex != nwg_global!"

df = pd.DataFrame(results)
print("Standard NW-flex matches nwg_global on all examples:")
print()
print(df.to_string(index=False))
print()
print("[OK] All base case tests passed.")
```

```
Standard NW-flex matches nwg_global on all examples:

                case          X        Y  flex  nwg match
           identical    GATTACA  GATTACA    35   35  [OK]
     single deletion    GATTACA   GATACA    10   10  [OK]
 single substitution       ACGT     ACCT    10   10  [OK]
   internal deletion GGGATATGGG GGGATGGG    19   19  [OK]
deletion + insertion      AACGT    ACGTT    -5   -5  [OK]

[OK] All base case tests passed.
```

##### 3.4 Single-block validation: explicit $A \cdot Z \cdot B$ example

We now validate the core claim: for references of the form

$$X = A \cdot Z \cdot B,$$

the single-block NW-flex configuration returns

$$S_{\text{flex}}(X, Y) = \max_{Z^* \subseteq Z} \text{NWG}(A \cdot Z^* \cdot B,\ Y).$$

##### 3.4.1 Defining the test sequences

```
# Define a small A·Z·B example
# Reference X and read Y
A = "ACT"
Z = "GATTACA"
B = "CAT"

X = A + Z + B
Zstar = "TTA"
Y = A + Zstar + B

print("A:", A)
print("Z:", Z)
print("B:", B)
print("X (reference):", X)
print("Y (read):     ", Y)

# Block boundary indices
s = len(A)          # leader row (last row of A)
e = len(A) + len(Z) # end of Z; closer row is e+1
n = len(X)

print(f"\nBlock indices: s = {s}, e = {e}")
print(f"  A = X[0:{s}] = '{X[:s]}'")
print(f"  Z = X[{s}:{e}] = '{X[s:e]}'")
print(f"  B = X[{e}:{n}] = '{X[e:]}'")
print(f"  Leader row: {s}, Closer row: {e+1}")
```

```
A: ACT
Z: GATTACA
B: CAT
X (reference): ACTGATTACACAT
Y (read):      ACTTTACAT

Block indices: s = 3, e = 10
  A = X[0:3] = 'ACT'
  Z = X[3:10] = 'GATTACA'
  B = X[10:13] = 'CAT'
  Leader row: 3, Closer row: 11
```

##### 3.4.2 The single-block EP pattern

Before running the alignment, let's visualize the extra-predecessor pattern $E(i)$ for this single-block configuration:

- Rows inside $Z$ (rows $s+1$ to $e$) have $E(i) = \{s\}$: they can jump from the **leader** row $s$.
- The **closer** row $e+1$ has $E(e+1) = \{s, s+1, \ldots, e\}$: it can jump from the leader or any row in $Z$.

- All other rows have $E(i) = \varnothing$.

```
# Build the EP pattern
EP = build_EP_single_block(n=n, s=s, e=e)

# Define A·Z·B region colors (matching notebook 02)
color_A, color_Z, color_B = "#d0e0ff", "#ffe0d0", "#e0d0f0"
label_A, label_Z, label_B = "#2060a0", "#c06020", "#7030a0"

regions = [
    {'start': 1, 'end': s, 'color': color_A, 'label': 'A', 'label_color':␣
 ↪label_A},
    {'start': s + 1, 'end': e, 'color': color_Z, 'label': 'Z', 'label_color':␣
 ↪label_Z},
    {'start': e + 1, 'end': n, 'color': color_B, 'label': 'B', 'label_color':␣
 ↪label_B},
]

# Visualize the pattern with A·Z·B region shading (matching notebook 02 style)
fig, ax = plot_ep_pattern(
    EP=EP,
    leaders=[s],
    X=X,
    title="Single-block EP pattern for A·Z·B",
    figsize=(5, 8),
    row_annotations={s: "(s)", e+1: "(e+1)"},
    node_height=0.7,
    row_spacing=1.2,
    node_width=2.0,
    sequence_fontsize=14.0,
    row_label_fontsize=10.0,
    sequence_fontweight="semibold",
    ep_linewidth=3.0,
    box_label_fontsize=12.0,
)

# Add the A·Z·B background regions
draw_ep_background(
    ax, n,
    regions=regions,
    row_spacing=1.2,
    node_height=0.7,
    node_width=2.0,
    region_fontsize=20.0,
    region_fontweight="bold",
)
```

```
plt.show()
```

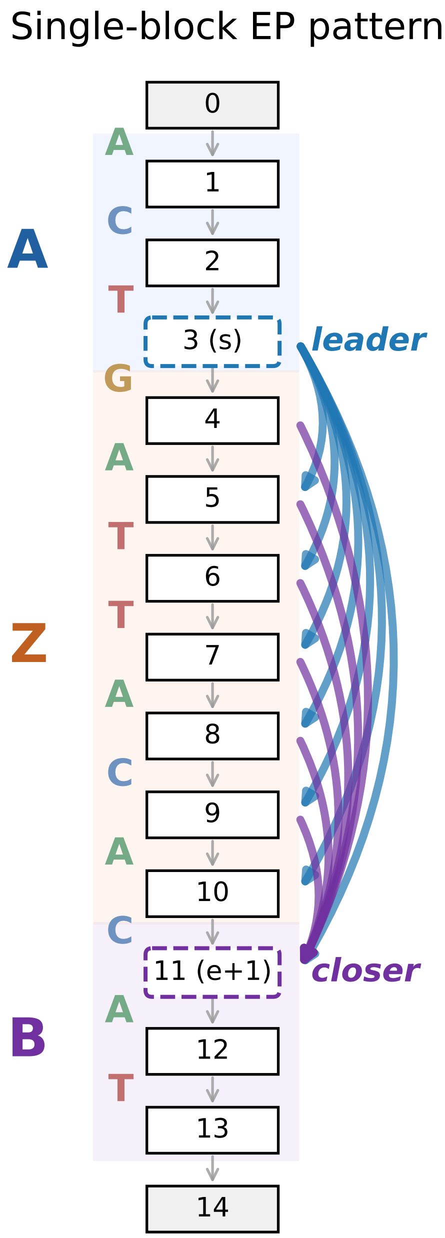

###### 3.4.3 Enumerate all substrings $Z^*$ and compute NWG scores

The naive baseline enumerates all contiguous substrings $Z^* = X[i : j]$ with $s \leq i \leq j \leq e$, forms the effective reference $X' = A \cdot Z^* \cdot B$, and computes $\mathrm{NWG}(X', Y)$.

The flex optimum is the maximum over all such substrings.

```python
naive_rows = []
best_score = -np.inf
best_pairs = []

for i in range(s, e + 1):
    for j in range(i, e + 1):
        Z_star = X[i:j]              # contiguous substring of Z
        X_prime = A + Z_star + B     # effective reference
        score = nwg_global(
            X_prime, Y,
            score_matrix, gap_open, gap_extend, alphabet_to_index,
        )
        naive_rows.append((i, j, Z_star, X_prime, score))
        if score > best_score:
            best_score = score
            best_pairs = [(i, j, Z_star, X_prime)]
        elif score == best_score:
            best_pairs.append((i, j, Z_star, X_prime))

# Build DataFrame for display
df_naive = pd.DataFrame(naive_rows, columns=["i", "j", "Z*", "X'=A·Z*·B",␣
↪"score"])

# Mark optimal rows
df_naive["opt"] = df_naive["score"].apply(lambda x: "*" if x == best_score else␣
↪"")

print(f"Naive enumeration of Z* substrings (best score = {int(best_score)}):")
print()
print(df_naive.to_string(index=False))

print(f"Total of {len(naive_rows)} Z* substrings evaluated.")
print(f"Total of {len(set(df_naive['Z*']))} distinct substrings evaluated.")

print(f"\nOptimal Z* choices achieving score {int(best_score)}:")
for i, j, zst, xp in best_pairs:
    print(f"  Z* = '{zst}' → X' = '{xp}'")
```

```
Naive enumeration of Z* substrings (best score = 45):

 i  j      Z*      X'=A·Z*·B  score opt
 3  3                ACTCAT    8.0
 3  4       G       ACTGCAT    4.0
 3  5      GA      ACTGACAT   10.0
 3  6     GAT     ACTGATCAT   15.0
 3  7    GATT    ACTGATTCAT    5.0
 3  8   GATTA   ACTGATTACAT   24.0
```

```
 3  9  GATTAC  ACTGATTACCAT    4.0
 3 10 GATTACA ACTGATTACACAT    3.0
 4  4                ACTCAT    8.0
 4  5       A       ACTACAT   14.0
 4  6      AT      ACTATCAT   10.0
 4  7     ATT     ACTATTCAT   25.0
 4  8    ATTA    ACTATTACAT   25.0
 4  9   ATTAC   ACTATTACCAT    5.0
 4 10  ATTACA  ACTATTACACAT   13.0
 5  5                ACTCAT    8.0
 5  6       T       ACTTCAT   14.0
 5  7      TT      ACTTTCAT   20.0
 5  8     TTA     ACTTTACAT   45.0   *
 5  9    TTAC    ACTTTACCAT   25.0
 5 10   TTACA   ACTTTACACAT   24.0
 6  6                ACTCAT    8.0
 6  7       T       ACTTCAT   14.0
 6  8      TA      ACTTACAT   20.0
 6  9     TAC     ACTTACCAT   25.0
 6 10    TACA    ACTTACACAT   15.0
 7  7                ACTCAT    8.0
 7  8       A       ACTACAT   14.0
 7  9      AC      ACTACCAT    0.0
 7 10     ACA     ACTACACAT   25.0
 8  8                ACTCAT    8.0
 8  9       C       ACTCCAT    4.0
 8 10      CA      ACTCACAT   10.0
 9  9                ACTCAT    8.0
 9 10       A       ACTACAT   14.0
10 10                ACTCAT    8.0
Total of 36 Z* substrings evaluated.
Total of 26 distinct substrings evaluated.

Optimal Z* choices achieving score 45:
  Z* = 'TTA' → X' = 'ACTTTACAT'
```

The table above shows all 36 possible substrings $Z^*$ of $Z =$ GATTACA (26 unique). The asterisks mark the optimal choice(s).

In this example, the best substring is $Z^* =$ TTA ($i = 5; j = 8$).

This yields $X' =$ ACTTTACAT — an exact match with the read $Y$ resulting in the highest possible score, $9 \times 5 = 45$ (all matches, no gaps).

##### 3.4.4 Single-block NW-flex on the same example

Now we run NW-flex with the single-block EP pattern on the same $(X, Y, s, e)$ and verify that it achieves the same score as the naive maximum.

```
res_blk = align_single_block(
    X=X, Y=Y, s=s, e=e,
    score_matrix=score_matrix,
    gap_open=gap_open,
    gap_extend=gap_extend,
    alphabet_to_index=alphabet_to_index,
    return_data=True,
)

print(f"NW-flex single-block score: {res_blk.score}")
print(f"Naive maximum score:        {best_score}")
print()
print("NW-flex alignment:")
print(f"  X_aln: {res_blk.X_aln}")
print(f"  Y_aln: {res_blk.Y_aln}")

assert res_blk.score == best_score, "NW-flex score does not match naive maximum!"
print("\n[OK] NW-flex single-block score equals the naive max over Z*.")
```

```
NW-flex single-block score: 45.0
Naive maximum score:        45.0

NW-flex alignment:
  X_aln: ACTTTACAT
  Y_aln: ACTTTACAT

[OK] NW-flex single-block score equals the naive max over Z*.
```

#### 3.5 Systematic validation with `check_AZB_case`

We have shown score equality on one explicit example. Now we use the helper `check_AZB_case(A, Z, B, Y)` to run both NW-flex and the naive baseline on multiple test cases.

##### 3.5.1 Manual test cases

```
# Interesting test cases that exercise different aspects of flex alignment:
manual_cases = [
    # (A, Z, B, Y, description)
    # Case 1: Read matches a proper substring of Z (contraction)
    ("ACG", "TATATAT", "ACG", "ACGTATACG",
     "Y has shorter repeat than Z"),
    # Case 2: Read matches flanks but skips Z entirely (Z* = ϵ)
    ("GATTACA", "CCCC", "GATTACA", "GATTACAGATTACA",
     "Y skips Z entirely"),
    # Case 3: Read has insertion relative to best Z*
    ("AT", "GCGCGC", "AT", "ATGCGGGCGCAT",
     "Y has insertion in Z region"),
    # Case 4: Z contains repeats, Y matches subset
```

```
    ("CAT", "AGAGAGAG", "CAT", "CATAGAGCAT",
     "Y matches partial repeat"),
    # Case 5: Flanks have mismatches, Z must compensate
    ("AAAA", "TGTGTG", "CCCC", "AAAATGTCCCC",
     "Flanks match, Z contracts"),
    # Case 6: Complex case with all features
    ("GCAT", "ATATATATAT", "GCAT", "GCATATATGCAT",
     "STR-like contraction"),
]

results = []
for A0, Z0, B0, Y0, desc in manual_cases:
    flex_score, naive_score = check_AZB_case(A0, Z0, B0, Y0)
    match = "[OK]" if flex_score == naive_score else "[FAIL]"
    results.append({
        "description": desc,
        "A": A0, "Z": Z0, "B": B0, "Y": Y0,
        "flex": int(flex_score), "naive": int(naive_score), "match": match
    })
    assert flex_score == naive_score, f"Mismatch: A={A0}, Z={Z0}, B={B0}, Y={Y0}"

df = pd.DataFrame(results)
print("Manual A·Z·B test cases: NW-flex vs naive baseline")
print()
print(df.to_string(index=False))
print()
print("[OK] All manual test cases passed.")
```

```
Manual A·Z·B test cases: NW-flex vs naive baseline

                description       A          Z       B             Y  flex
naive match
Y has shorter repeat than Z     ACG    TATATAT     ACG     ACGTATACG    45
45   [OK]
         Y skips Z entirely GATTACA       CCCC GATTACA GATTACAGATTACA    70
70   [OK]
Y has insertion in Z region      AT     GCGCGC      AT   ATGCGGGCGCAT    29
29   [OK]
   Y matches partial repeat     CAT   AGAGAGAG     CAT     CATAGAGCAT    50
50   [OK]
  Flanks match, Z contracts    AAAA     TGTGTG    CCCC    AAAATGTCCCC    55
55   [OK]
       STR-like contraction    GCAT ATATATATAT    GCAT   GCATATATGCAT    60
60   [OK]

[OK] All manual test cases passed.
```

##### 3.5.2 Manual test cases with mutations

To stress-test the validation further, we introduce random mutations to the read sequence $Y$ from each base case above.
The `run_mutated_AZB_tests` function generates mutated variants by applying:

- **Substitutions** at rate 15% per base (each base has a 15% chance of being replaced by a different nucleotide),
- **Insertions** at rate 8% per base (random base inserted after),
- **Deletions** at rate 8% per base (base removed).

For each of the 6 manual cases, we generate 100 mutated variants of $Y$ and verify that NW-flex still matches the naive baseline on every mutated input.

```
# Run mutated tests on the interesting cases
results = run_mutated_AZB_tests(
    manual_cases,
    num_mutations=NUM_MUTATION_VARIANTS,
    sub_rate=0.15,
    indel_rate=0.08,
    seed=888,
)

df_mutated = pd.DataFrame(results)

# Summary statistics
n_total = len(df_mutated)
n_match = df_mutated["match"].sum()

print(f"Mutated sequence validation: {n_match}/{n_total} tests passed")
print(f"Substitution rate: 15%, Indel rate: 8%")
print()

# Aggregate by base case (description, A, Z, B, Y), preserving original order
summary = (
    df_mutated
    .groupby(["description", "A", "Z", "B", "Y"], sort=False)
    .agg(passed=("match", "sum"), total=("match", "count"))
    .reset_index()
)

def format_result(r):
    if r['passed'] == r['total']:
        return f"[OK] {r['passed']}/{r['total']}"
    else:
        return f"[FAIL] {r['passed']}/{r['total']}"

summary["result"] = summary.apply(format_result, axis=1)
```

```
print(f"Mutation test summary by base case ({n_match}/{n_total} total passed):")
print()
cols = ["description", "A", "Z", "B", "Y", "result"]
print(summary[cols].to_string(index=False))
```

```
Mutated sequence validation: 600/600 tests passed
Substitution rate: 15%, Indel rate: 8%

Mutation test summary by base case (600/600 total passed):

               description       A          Z       B              Y
result
Y has shorter repeat than Z     ACG    TATATAT     ACG      ACGTATACG  [OK]
100/100
         Y skips Z entirely GATTACA       CCCC GATTACA GATTACAGATTACA  [OK]
100/100
Y has insertion in Z region      AT     GCGCGC      AT    ATGCGGGCGCAT  [OK]
100/100
   Y matches partial repeat     CAT   AGAGAGAG     CAT     CATAGAGCAT  [OK]
100/100
  Flanks match, Z contracts    AAAA     TGTGTG    CCCC    AAAATGTCCCC  [OK]
100/100
       STR-like contraction    GCAT ATATATATAT    GCAT   GCATATATGCAT  [OK]
100/100
```

##### 3.5.3 Randomized test cases

We generate small random $A \cdot Z \cdot B$ examples and verify that NW-flex always matches the naive baseline. Sequences are kept short so that the $O(|Z|^2)$ naive enumeration remains fast.

```
DNA = "ACGT"

def random_dna(rng, length: int) -> str:
    return "".join(rng.choice(list(DNA)) for _ in range(length))

def run_random_AZB_tests(num_cases: int = 20, seed: int = 0):
    """Run randomized A·Z·B validation tests."""
    rng = np.random.default_rng(seed)
    for idx in range(num_cases):
        # Random lengths (small to keep naive enumeration cheap)
        len_A = rng.integers(4, 6)
        len_Z = rng.integers(4, 10)
        len_B = rng.integers(4, 6)
        len_Y = rng.integers(6, 10)

        A0 = random_dna(rng, len_A)
        Z0 = random_dna(rng, len_Z)
        B0 = random_dna(rng, len_B)
```

```
        Y0 = random_dna(rng, len_Y)

        flex_score, naive_score = check_AZB_case(A0, Z0, B0, Y0)
        if flex_score != naive_score:
            print(f"Mismatch: A={A0}, Z={Z0}, B={B0}, Y={Y0}")
            print(f"  flex={flex_score}, naive={naive_score}")
            raise AssertionError("Random A·Z·B test failed.")
    print(f"[OK] All {num_cases} random A·Z·B tests passed.")
```

```
random_seed = random.randint(0, 10000)
print(f"Random seed: {random_seed}")
run_random_AZB_tests(num_cases=NUM_RANDOM_CASES, seed=random_seed)
```

```
Random seed: 5489

[OK] All 100 random A·Z·B tests passed.
```

##### 3.6 Alignment string comparison

So far we have validated that **scores** match. The paper also claims that **alignments** are valid. We use `check_alignment_validity` to verify that the aligned sequences from NW-flex are consistent with the naive baseline.

Note: there may be multiple optimal alignments with the same score. We verify that both methods produce valid alignments achieving the optimal score, not necessarily that they produce *identical* alignment strings.

```
# Test alignment validity on our explicit example
res = align_single_block(
    X=A + Z + B,
    Y=Y,
    s=len(A),
    e=len(A) + len(Z),
    score_matrix=score_matrix,
    gap_open=gap_open,
    gap_extend=gap_extend,
    alphabet_to_index=alphabet_to_index,
)

valid, msg = check_alignment_validity(res)

print("Alignment validation for explicit example:")
print(f"  A = '{A}', Z = '{Z}', B = '{B}', Y = '{Y}'")
print()
print(f"NW-flex alignment (score = {res.score}):")
print(f"  X_aln: {res.X_aln}")
print(f"  Y_aln: {res.Y_aln}")
print()
```

```
print(f"Validity check: {msg}")
if valid:
    print("[OK] Alignment validation passed.")
```

```
Alignment validation for explicit example:
  A = 'ACT', Z = 'GATTACA', B = 'CAT', Y = 'ACTTTACAT'

NW-flex alignment (score = 45.0):
  X_aln: ACTTTACAT
  Y_aln: ACTTTACAT

Validity check: Valid alignment of length 9
[OK] Alignment validation passed.
```

##### 3.6.1 Testing with different scoring schemes

The paper states that validation was performed "across a range of scoring schemes." We verify score equality with alternative gap penalties.

```
# Alternative scoring schemes to test
scoring_variants = [
    ("Default", score_matrix, -20.0, -1.0),
    ("Smaller gap open", score_matrix, -10.0, -1.0),
    ("Larger gap extend", score_matrix, -20.0, -3.0),
    ("Balanced gaps", score_matrix, -4.0, -4.0),
]

results = []
for name, sm, go, ge in scoring_variants:
    # Run a few random tests with this scoring
    rng = np.random.default_rng(888)

    passed = 0
    for _ in range(NUM_SCORING_TESTS):
        len_A = rng.integers(4, 6)
        len_Z = rng.integers(4, 10)
        len_B = rng.integers(4, 6)
        len_Y = rng.integers(6, 10)

        A0 = random_dna(rng, len_A)
        Z0 = random_dna(rng, len_Z)
        B0 = random_dna(rng, len_B)
        Y0 = random_dna(rng, len_Y)

        flex_score, naive_score = check_AZB_case(
            A0, Z0, B0, Y0,
            score_matrix=sm,
            gap_open=go,
```

```
            gap_extend=ge,
        )
        if flex_score == naive_score:
            passed += 1

    status = "[OK]" if passed == NUM_SCORING_TESTS else "[FAIL]"
    results.append({
        "scheme": name,
        "gap_open": int(go),
        "gap_extend": int(ge),
        "result": f"{status} {passed}/{NUM_SCORING_TESTS}"
    })

df_scoring = pd.DataFrame(results)
n_tests = NUM_SCORING_TESTS
print(f"Validation with different scoring schemes ({n_tests} random tests each):
 ↪")
print()
print(df_scoring.to_string(index=False))
```

```
Validation with different scoring schemes (100 random tests each):

           scheme  gap_open  gap_extend       result
          Default       -20          -1 [OK] 100/100
 Smaller gap open       -10          -1 [OK] 100/100
Larger gap extend       -20          -3 [OK] 100/100
    Balanced gaps        -4          -4 [OK] 100/100
```

##### 3.7 Visualizing the DP matrices and row jumps

For the explicit $A \cdot Z \cdot B$ example, we visualize the DP matrices and the optimal path to see how NW-flex realizes the substring selection.

The path shows: - Standard DP in the flank $A$, - A jump from the leader row $s$ into the flex block $Z$, - Standard DP within the chosen substring $Z^*$, - A jump from $Z$ to the closer row $e{+}1$, - Standard DP in the flank $B$.

The **row jumps** reveal which substring $Z^*$ was selected. From the jump coordinates we can recover the optimal indices $(a, b)$ such that $Z^* = Z[a : b]$ (in Python slice notation):

- The **entry jump** (into $Z$) lands at row $i_{\text{in}}$; the start index is $a = i_{\text{in}} - s - 1$.
- The **exit jump** (to the closer row $e{+}1$) departs from row $i_{\text{out}}$; the end index is $b = i_{\text{out}} - s$.

```
# Show row jumps from the alignment result
print("Row jumps in the optimal path:")
if res_blk.jumps:
    for jump in res_blk.jumps:
        print(f"  Row {jump.from_row} → Row {jump.to_row} at column {jump.col}")
else:
```

```
    print("  (no jumps - alignment stayed within standard predecessor␣
↪structure)")

# Convert jump coordinates to Z* indices
# Entry jump: from leader s into Z; exit jump: from Z to closer e+1
zstar_rows = None
if len(res_blk.jumps) >= 2:
    entry_jump = res_blk.jumps[0]   # jump into Z
    exit_jump = res_blk.jumps[1]    # jump out of Z to closer

    # a = first row of Z* relative to Z (0-indexed into Z)
    # b = one past last row of Z* relative to Z
    a = entry_jump.to_row - s - 1
    b = exit_jump.from_row - s

    print(f"\nRecovered Z* indices from jumps:")
    entry_row = entry_jump.to_row
    print(f"  Entry jump lands at row {entry_row} → a = {entry_row} - {s} - 1 =␣
↪{a}")
    exit_row = exit_jump.from_row
    print(f"  Exit jump departs from row {exit_row} → b = {exit_row} - {s} =␣
↪{b}")
    print(f"  Z[{a}:{b}] = '{Z[a:b]}'")
    zstar = Z[a:b]
    print(f"\nVerification: Z* = '{zstar}' matches the optimal substring.")

    # Store Z* row bounds for region annotation
    zstar_start = entry_jump.to_row
    zstar_end = exit_jump.from_row

# Build A·Z·B regions for background fills

regions = [
    {'start': 1, 'end': s, 'label': 'A',
     'fill_color': REGION_FILL_COLORS['A'], 'label_color':␣
↪REGION_LABEL_COLORS['A']},
    {'start': s + 1, 'end': e, 'label': 'Z',
     'fill_color': REGION_FILL_COLORS['Z'], 'label_color':␣
↪REGION_LABEL_COLORS['Z']},
    {'start': e + 1, 'end': n, 'label': 'B',
     'fill_color': REGION_FILL_COLORS['B'], 'label_color':␣
↪REGION_LABEL_COLORS['B']},
] if s >= 1 else []

# Build Z* highlight (gold dotted box like leader/closer)
GOLD = '#DAA520'
```

```
row_highlights = []
if len(res_blk.jumps) >= 2:
    row_highlights.append({
        'start': zstar_start,
        'end': zstar_end,
        'label': 'Z*',
        'color': GOLD,
        'linestyle': ':',
        'label_position': 'top',
    })

# Create figure with GridSpec for the 3 matrices
fig = plt.figure(figsize=(10, 5))
gs = gridspec.GridSpec(1, 3, wspace=0.08)
axes = [fig.add_subplot(gs[i]) for i in range(3)]

# Plot using publication-quality plotter
plot_flex_matrices_publication(
    fig=fig,
    axes=axes,
    result=res_blk,
    X=X,
    Y=Y,
    s=s,
    e=e,
    regions=regions,
    row_highlights=row_highlights,
    colormap=grid_color_map,
    title_pad=18,
)

fig.suptitle("Single-block NW-flex DP matrices with optimal path", fontsize=14,␣
 ↪y=1.02)
plt.show()
```

```
Row jumps in the optimal path:
  Row 3 → Row 6 at column 4
  Row 8 → Row 11 at column 7

Recovered Z* indices from jumps:
  Entry jump lands at row 6 → a = 6 - 3 - 1 = 2
  Exit jump departs from row 8 → b = 8 - 3 = 5
  Z[2:5] = 'TTA'

Verification: Z* = 'TTA' matches the optimal substring.
```

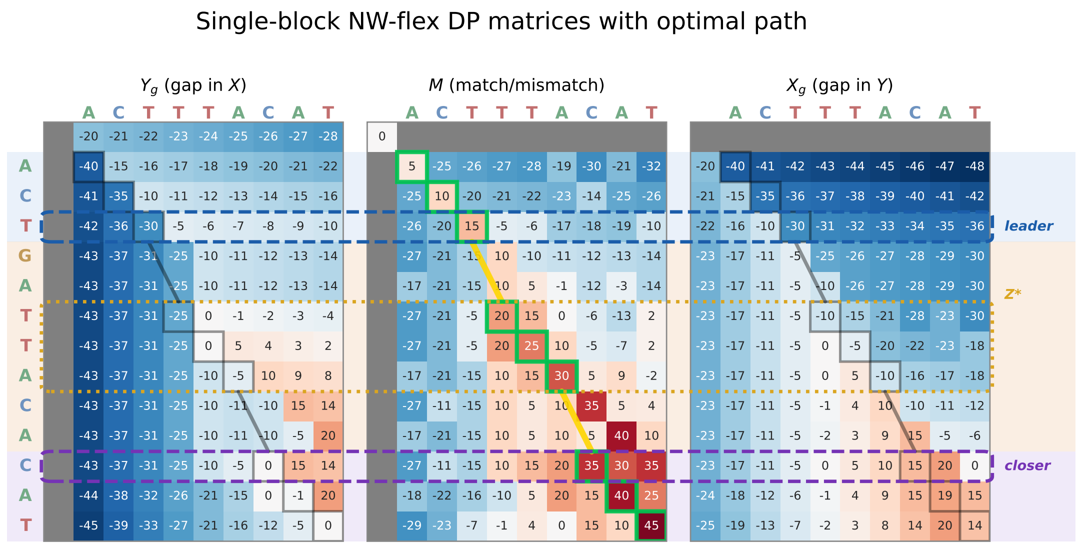

##### 3.8 Summary

This notebook validated the core NW-flex optimality claim for the single-block $A{\cdot}Z{\cdot}B$ configuration:

1. **Base case**: NW-flex with $E(i) = \varnothing$ matches an independent NWG implementation on all tested examples.
2. **Single-block flex**: NW-flex with the prescribed EP pattern achieves

$$S_{\text{flex}}(X, Y) = \max_{Z^* \subseteq Z} \text{NWG}(A \cdot Z^* \cdot B,\ Y)$$

   on both hand-crafted and randomized test cases.
3. **Alignment correctness**: The alignment strings from NW-flex are valid and consistent with the naive baseline.
4. **Scoring robustness**: Score equality holds across different gap penalty configurations.
5. **Path interpretation**: The DP visualizations show how NW-flex realizes the optimal $Z^*$ via row jumps at the leader and closer boundaries.

This validates the flex optimality guarantee for single-block configurations. In **Notebook 4**, we build on this foundation to handle STR-specific EP patterns with motif phase constraints.

### 4 Notebook 4: STR Specialization and Phase-Preserving Alignment

This notebook extends the NW-flex framework to handle **short tandem repeats** (STRs), where the flexible block $Z$ consists of repeated copies of a short motif $R$. We develop a phase-preserving EP pattern that correctly aligns reads whose repeat content may start or end mid-motif.

#### 4.1 Overview

In this notebook we:

1. **Define phased repeats** $Z^* = \mathrm{suf}(R, a) \cdot R^M \cdot \mathrm{pre}(R, b)$, representing repeat regions that may enter or exit the motif at arbitrary phases.
2. **Enumerate valid phase combinations** $(a, b, M)$ under the constraint $M + \mathbf{1}_{\{a>0\}} + \mathbf{1}_{\{b>0\}} \leq N$.
3. **Build the STR-specific EP pattern** using `build_EP_STR_phase`, which allows entry from the leader row and exit only through phase-compatible rows.
4. **Validate correctness** by verifying that NW-flex produces the expected perfect-match score for all valid phase combinations.
5. **Demonstrate phase-wise monotonicity** in the DP grid: for fixed column $j$ and motif phase $r$, scores along rows $i = s + 1 + r + nk$ are non-decreasing.
6. **Extend to multi-block STRs** (compound repeats) using `build_EP_multi_STR_phase`.

This notebook builds on:

- **Notebook 1**: Needleman–Wunsch basics and the Gotoh three-state DP.
- **Notebook 2**: NW-flex core with extra-predecessor (EP) patterns.
- **Notebook 3**: Validation of NW-flex optimality for single-block $A \cdot Z \cdot B$.

#### 4.2 Setup and imports

```
#  Notebook magic: autoreload modules
%load_ext autoreload
%autoreload 2

import pandas as pd
import matplotlib.pyplot as plt
import matplotlib.gridspec as gridspec
from IPython.display import display

# STR repeat utilities
from nwflex.repeats import (
    STRLocus,
    CompoundSTRLocus,
    phase_repeat,
    infer_abM_from_jumps,
)

# EP pattern builders
from nwflex.ep_patterns import (
    build_EP_standard,
    build_EP_single_block,
    build_EP_STR_phase,
    build_EP_multi_STR_phase,
```

```
)

# Alignment functions
from nwflex.aligners import align_with_EP

# Scoring and validation
from nwflex.validation import get_default_scoring

# Plotting
from nwflex.plot import (
    plot_flex_matrices_publication,
    plot_ep_comparison,
    REGION_FILL_COLORS,
    REGION_LABEL_COLORS,
)

from nwflex.plot.colors import HEATMAP_COLORMAPS
grid_color_map = HEATMAP_COLORMAPS['diverging']

# Set pandas display options
pd.set_option('display.max_colwidth', 30)
```

```
The autoreload extension is already loaded. To reload it, use:
  %reload_ext autoreload
```

##### 4.2.1 Scoring scheme

We use the same default scoring as in the other notebooks: match = +5, mismatch = -5, gap-open = -20, gap-extend = -1.

```
# Load default scoring scheme
score_matrix, gap_open, gap_extend, alphabet_to_index = get_default_scoring()

# Extract match score for later validation
match_score = score_matrix[alphabet_to_index['A'], alphabet_to_index['A']]

print(f"Scoring: match = {match_score}, mismatch = {score_matrix[0,1]}, "
      f"gap_open = {gap_open}, gap_extend = {gap_extend}")
```

```
Scoring: match = 5.0, mismatch = -5.0, gap_open = -20.0, gap_extend = -1.0
```

#### 4.3 Defining an STR locus

We consider an STR locus of the form

$$X = A \cdot R^N \cdot B,$$

where:

- $A$ is the left flank sequence,

- $R$ is the repeat motif of length $k = |R|$,
- $N$ is the number of motif copies in the reference,
- $B$ is the right flank sequence.

In our running example we use:

- $A =$ GAG
- $R =$ ACT (motif length $k = 3$)
- $N = 6$
- $B =$ GTCA

The STRLocus class from nwflex.repeats encapsulates this structure and provides convenient properties for the derived quantities.

```
# Define the STR locus
locus = STRLocus(A="GAG", R="ACT", N=6, B="GTCA")

print(locus)
print()
print(f"Reference X: {locus.X}")
```

```
STRLocus(A='GAG', R='ACT', N=6, B='GTCA')
  X = 'GAGACTACTACTACTACTACTGTCA' (n=25)
  s=3, e=21, k=3

Reference X: GAGACTACTACTACTACTACTGTCA
```

##### 4.3.1 Phased repeats inside a read

A read $Y$ from this locus may have a repeat region $Z^*$ that differs from the reference $Z = R^N$. In particular, $Z^*$ may:

- Start mid-motif (e.g., begin at the second base of $R$),
- End mid-motif (e.g., include only the first base of the final $R$),
- Have a different number of complete motif copies.

To describe such phased repeats, we define:

- $\mathrm{pre}(R, b)$: the length-$b$ **prefix** of $R$,
- $\mathrm{suf}(R, a)$: the length-$a$ **suffix** of $R$.

Given integers $a, b, M \geq 0$, we define a **phased repeat block**:

$$Z^* = \mathrm{phase}(R, a, b, M) = \mathrm{suf}(R, a) \cdot R^M \cdot \mathrm{pre}(R, b).$$

Intuitively: - $Z^*$ may start in the middle of a motif (suffix of length $a$), - contain $M$ full copies of $R$, - and end in the middle of the next motif (prefix of length $b$).

The corresponding read is $Y = A \cdot Z^* \cdot B$.

```
# Example: a = 2, b = 1, M = 2
a, b, M = 2, 1, 2
```

```python
# Build Z* using phase_repeat from the library
Zstar = phase_repeat(locus.R, a, b, M)

# Build the read Y
Y = locus.build_locus_variant(a, b, M)

print(f"Motif R = '{locus.R}' (k = {locus.k})")
print(f"Parameters: a = {a}, b = {b}, M = {M}")
print()
print(f"suf(R, {a}) = '{locus.R[-a:] if a > 0 else ''}'")
print(f"R^{M} = '{locus.R * M}'")
print(f"pre(R, {b}) = '{locus.R[:b] if b > 0 else ''}'")
print()
print(f"Z* = phase(R, {a}, {b}, {M}) = '{Zstar}'")
print(f"Y  = A + Z* + B = '{Y}'")
```

```
Motif R = 'ACT' (k = 3)
Parameters: a = 2, b = 1, M = 2

suf(R, 2) = 'CT'
R^2 = 'ACTACT'
pre(R, 1) = 'A'

Z* = phase(R, 2, 1, 2) = 'CTACTACTA'
Y  = A + Z* + B = 'GAGCTACTACTAGTCA'
```

###### 4.3.2 The phase constraint

We restrict attention to reads $Y = A \cdot Z^* \cdot B$ whose repeat content does not exceed that of the reference $X = A \cdot R^N \cdot B$. A convenient way to express this is:

$$M + \mathbf{1}_{\{a>0\}} + \mathbf{1}_{\{b>0\}} \ \leq \ N,$$

where $\mathbf{1}_{\{a>0\}}$ is 1 if $a > 0$ and 0 otherwise.

In words: - Each full motif contributes 1 to $M$, - A nonempty suffix $\mathrm{suf}(R, a)$ behaves like an additional partial repeat on the left, - A nonempty prefix $\mathrm{pre}(R, b)$ behaves like an additional partial repeat on the right.

Thus the maximal allowed $Z^*$ (in terms of repeat content) is any phased block with parameters $(a, b, M)$ satisfying this constraint.

###### 4.3.3 Enumerating all valid phase combinations

Given the constraint $M + \mathbf{1}_{\{a>0\}} + \mathbf{1}_{\{b>0\}} \leq N$, we can enumerate all valid $(a, b, M)$ triples that produce a phased repeat $Z^*$ compatible with the reference.

For each combination: - $a \in \{0, 1, \dots, k-1\}$ determines the entry phase (suffix length of $R$), - $b \in \{0, 1, \dots, k-1\}$ determines the exit phase (prefix length of $R$), - $M \geq 0$ is the number of complete motif copies.

The `valid_combinations()` function generates all such triples.

```
# Generate all valid (a, b, M) combinations for this locus
valid_combos = list(locus.valid_combinations())
n_combos = len(valid_combos)

print(f"Valid (a, b, M) combinations for N={locus.N}, k={locus.k}: {n_combos}")
print()

# Build a table showing all combinations
results = []
for a_i, b_i, M_i in valid_combos:
    Zstar_i = phase_repeat(locus.R, a_i, b_i, M_i)
    Y_i = locus.build_locus_variant(a_i, b_i, M_i)
    overhead = (1 if a_i > 0 else 0) + (1 if b_i > 0 else 0)
    results.append({
        'a': a_i,
        'b': b_i,
        'M': M_i,
        'overhead': overhead,
        'total': M_i + overhead,
        '|Z*|': len(Zstar_i),
        'Z*': Zstar_i,
    })

df_combos = pd.DataFrame(results)
caption = f"All valid (a, b, M) combinations for R = '{locus.R}', N = {locus.N}"
styled = df_combos.style\
    .set_caption(caption)\
    .set_properties(**{'white-space': 'nowrap', 'padding': '2px 6px'})

display(styled)
```

```
Valid (a, b, M) combinations for N=6, k=3: 51

<pandas.io.formats.style.Styler at 0x7c13cb4006d0>
```

#### 4.4 STR-specific EP pattern

We now build the STR-specific EP pattern that allows the DP to discover the optimal phase parameters $(a, b, M)$ automatically.

Recall from Notebook 2 that for a single flexible block $X = A \cdot Z \cdot B$, we define: - **Leader row** $s = |A|$: the last row of flank $A$, - **Closer row** $e + 1$: the first row of flank $B$, where $e = |A| + |Z|$.

For the STR case where $Z = R^N$, we modify the exit condition to preserve motif phase:

**Entry into the block**: For rows $i \in \{s+1, \ldots, e\}$ inside the repeat block, we set $E(i) = \{s\}$. This allows the alignment to jump from the leader row $s$ into any position within the block, effectively choosing the entry phase $a$.

**Exit from the block**: At the closer row $e+1$, we set

$$E(e+1) = \{s,\ e-k+1,\ e-k+2,\ \ldots,\ e-1\}.$$

The key insight is that only the last $k$ rows of the block (plus the leader $s$) can exit to row $e+1$. This restriction enforces **phase-compatible exits**: the departure row $i$ determines the exit phase $b = (i - e) \mod k$.

The leader $s$ appears in $E(e+1)$ to handle the case where the read has no repeat content at all ($a = b = M = 0$), allowing the alignment to skip the entire block.

```
# Build the three EP patterns for comparison
n = locus.n
s, e, k = locus.s, locus.e, locus.k

EP_standard = build_EP_standard(n)
EP_block = build_EP_single_block(n, s, e)
EP_str = build_EP_STR_phase(n, s, e, k)

print(f"Reference length n = {n}")
print(f"Leader row s = {s}, Closer row e+1 = {e+1}, Motif length k = {k}")
print()

# Show the EP sets for the closer row
print("Extra predecessors at closer row (e+1):")
print(f"  Single-block: E({e+1}) = {EP_block[e+1]}")
print(f"  STR-phase:    E({e+1}) = {EP_str[e+1]}")
```

```
Reference length n = 25
Leader row s = 3, Closer row e+1 = 22, Motif length k = 3

Extra predecessors at closer row (e+1):
  Single-block: E(22) = [3, 4, 5, 6, 7, 8, 9, 10, 11, 12, 13, 14, 15, 16, 17,
18, 19, 20]
  STR-phase:    E(22) = [3, 19, 20]
```

##### 4.4.1 Visualizing the EP patterns

The figure below compares the three EP configurations:

1. **Standard NW**: No extra predecessors ($E(i) = \varnothing$ for all $i$).
2. **Single-block flex**: All rows in $Z$ can exit to the closer.
3. **STR-phase flex**: Only the last $k$ rows of $Z$ (plus leader) can exit.

```
# Colors for background regions
color_A, color_Z, color_B, color_X = "#d0e0ff", "#ffe0d0", "#e0d0f0", "#e0e0e0"
label_A, label_Z, label_B, label_X = "#2060a0", "#c06020", "#7030a0", "#404040"
```

```
# Background regions for each pattern
bg_standard = [
    {'start': 1, 'end': n, 'color': color_X,
     'label': 'X', 'label_color': label_X}
]
bg_azb = [
    {'start': 1, 'end': s, 'color': color_A,
     'label': 'A', 'label_color': label_A},
    {'start': s + 1, 'end': e, 'color': color_Z,
     'label': 'Z', 'label_color': label_Z},
    {'start': e + 1, 'end': n, 'color': color_B,
     'label': 'B', 'label_color': label_B},
]
bg_str = [
    {'start': 1, 'end': s, 'color': color_A,
     'label': 'A', 'label_color': label_A},
    {'start': s + 1, 'end': e, 'color': color_Z,
     'label': '$R^N$', 'label_color': label_Z},
    {'start': e + 1, 'end': n, 'color': color_B,
     'label': 'B', 'label_color': label_B},
]

# Titles
titles = [
    "Standard NW\n$E(i) = \\varnothing$",
    "Single-block flex\n$E(e+1) = \\{s, s{+}1, \\ldots, e{-}1\\}$",
    f"STR-phase flex (k={k})\n$E(e+1) = \\{{s, e{{-}}k{{+}}1, \\ldots,␣
 ↪e{{-}}1\\}}$",
]

# Row annotations
annot_std = {}
annot_azb = {s: "(s)", e + 1: "(e+1)"}

fig = plot_ep_comparison(
    EP_list=[EP_standard, EP_block, EP_str],
    leaders_list=[[], [s], [s]],
    titles=titles,
    X=locus.X,
    backgrounds=[bg_standard, bg_azb, bg_str],
    row_annotations_list=[annot_std, annot_azb, annot_azb],
    figsize=(14, 18),
    wspace=0.05,
    closer_edge_color="#7030a0",
    leader_curvature=0.3,
    closer_curvature=0.25,
    ep_linewidth=3.0,
```

```
    sequence_fontsize=14.0,
    sequence_fontweight="semibold",
    region_fontsize=24.0,
    row_label_fontsize=10.0,
    row_label_fontweight="normal",
    legend_fontsize=14.0,
    row_spacing=1.2,
    node_height=0.7,
    legend_bbox_y=0.04,
    bottom_margin=0.08,
)
plt.show()
```

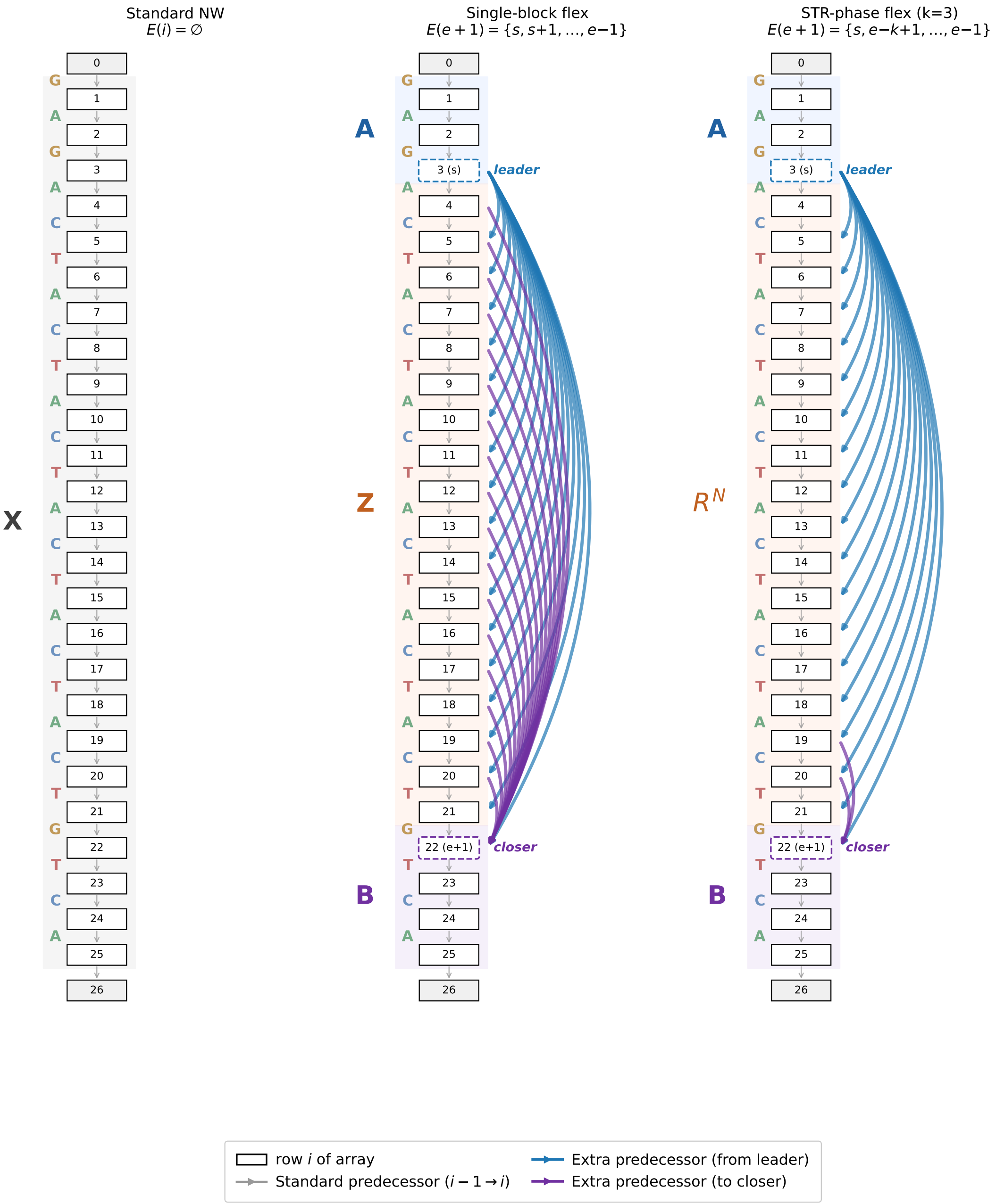

#### 4.5 Validating NW-flex for all phase combinations

Having defined the STR EP pattern, we now verify that it correctly handles **all** valid phase combinations $(a, b, M)$ satisfying the constraint $M + \mathbf{1}_{\{a>0\}} + \mathbf{1}_{\{b>0\}} \leq N$.

For any such triple, the read $Y = A \cdot Z^* \cdot B$ should align perfectly to the reference $X = A \cdot R^N \cdot B$, achieving a score of $|Y| \times \text{match\_score}$ (with no gap penalties in the repeat region).

We verify this by: 1. Iterating over all valid $(a, b, M)$ combinations, 2. Constructing the corresponding $Z^*$ and read $Y$, 3. Running NW-flex with the STR EP pattern, 4. Checking that the alignment score matches the expected perfect-match score.

```
# Validate all phase combinations
validation_results = []

for a_i, b_i, M_i in valid_combos:
    # Build Z* and Y
    Zstar_i = phase_repeat(locus.R, a_i, b_i, M_i)
    Y_i = locus.build_locus_variant(a_i, b_i, M_i)

    # Expected score: perfect match of all |Y| bases
    expected_score = match_score * len(Y_i)

    # Run NW-flex with STR EP pattern
    result = align_with_EP(
        X=locus.X,
        Y=Y_i,
        score_matrix=score_matrix,
        gap_open=gap_open,
        gap_extend=gap_extend,
        alphabet_to_index=alphabet_to_index,
        extra_predecessors=EP_str,
        return_data=False,
    )

    # Check if score matches expected
    passed = result.score == expected_score

    validation_results.append({
        'a': a_i,
        'b': b_i,
        'M': M_i,
        '|Z*|': len(Zstar_i),
        'Z*': Zstar_i,
        '|Y|': len(Y_i),
        'Expected': int(expected_score),
        'Score': int(result.score),
        'Match': "[OK]" if passed else "[FAIL]",
    })

# Display results
df_validation = pd.DataFrame(validation_results)
caption = f"NW-flex validation: R = '{locus.R}' (k={locus.k}), N = {locus.N}"
styled = df_validation.style\
    .set_caption(caption)\
    .set_properties(**{'white-space': 'nowrap', 'padding': '2px 6px'})
```

```
display(styled)

# Summary
n_pass = sum(1 for r in validation_results if r['Match'] == "[OK]")
n_total = len(validation_results)
print(f"\nPassed: {n_pass}/{n_total} combinations")

# Show any failures
failures = [r for r in validation_results if r['Match'] == "[FAIL]"]
if failures:
    print("\nFailing cases:")
    for r in failures:
        exp, got = r['Expected'], r['Score']
        print(f"  a={r['a']}, b={r['b']}, M={r['M']}: expected={exp}, got={got}")
```

```
<pandas.io.formats.style.Styler at 0x7c13ca584750>

Passed: 51/51 combinations
```

#### 4.6 Running NW-flex on an example STR alignment

We now run NW-flex on a specific example and examine the results in detail. The traceback records row jumps when the alignment uses an extra predecessor edge. For an STR locus:

- **Entry phase** $a$: When the traceback jumps from the leader row $s$ into the block, the landing row determines $a$.
- **Exit phase** $b$: When the traceback jumps from some row $i$ in the block to the closer row $e + 1$, the departure row determines $b$.
- **Complete units** $M$: The number of complete motif copies is computed from the total length of $Z^*$ and the phases $a$, $b$.

The `infer_abM_from_jumps()` function recovers these parameters from the traceback.

```
# Run alignment with our example (a=2, b=1, M=2)
result = align_with_EP(
    X=locus.X,
    Y=Y,
    score_matrix=score_matrix,
    gap_open=gap_open,
    gap_extend=gap_extend,
    alphabet_to_index=alphabet_to_index,
    extra_predecessors=EP_str,
    return_data=True,
)

print(f"Alignment score: {result.score}")
print(f"Aligned X: {result.X_aln}")
```

```
print(f"Aligned Y: {result.Y_aln}")
print()
print(f"Row jumps: {result.jumps}")
```

```
Alignment score: 80.0
Aligned X: GAGCTACTACTAGTCA
Aligned Y: GAGCTACTACTAGTCA

Row jumps: [RowJump(from_row=3, to_row=11, col=4, state=1), RowJump(from_row=19,
to_row=22, col=13, state=1)]
```

```
# Infer (a, b, M) from the row jumps
a_inf, b_inf, M_inf = infer_abM_from_jumps(result.jumps, locus.s, locus.e, locus.
↪k)

if a_inf is not None:
    Zstar_inf = phase_repeat(locus.R, a_inf, b_inf, M_inf)
else:
    Zstar_inf = "?"

print("Phase inference from traceback:")
print(f"  Original (a, b, M) = ({a}, {b}, {M})")
print(f"  Inferred (a, b, M) = ({a_inf}, {b_inf}, {M_inf})", end="")
print(f" {'[OK]' if (a, b, M) == (a_inf, b_inf, M_inf) else '[FAIL]'}")
print()
print(f"  Original Z* = '{Zstar}'")
print(f"  Inferred Z* = '{Zstar_inf}'", end="")
print(f" {'[OK]' if Zstar_inf == Zstar else '[FAIL]'}")
```

```
Phase inference from traceback:
  Original (a, b, M) = (2, 1, 2)
  Inferred (a, b, M) = (2, 1, 2) [OK]

  Original Z* = 'CTACTACTA'
  Inferred Z* = 'CTACTACTA' [OK]
```

##### 4.6.1 Visualizing the DP matrices

We plot the three DP layers ($Y_g$, $M$, $X_g$) with the optimal path overlaid. The row jumps show how the alignment enters and exits the repeat block.

```
# Plot the DP matrices with alignment path using publication-quality plotter
n = len(locus.X)

# Build A·Z·B regions for background fills
regions = [
    {'start': 1, 'end': locus.s, 'label': 'A',
```

```
     'fill_color': REGION_FILL_COLORS['A'], 'label_color':␣
↪REGION_LABEL_COLORS['A']},
    {'start': locus.s + 1, 'end': locus.e, 'label': 'Z',
     'fill_color': REGION_FILL_COLORS['Z'], 'label_color':␣
↪REGION_LABEL_COLORS['Z']},
    {'start': locus.e + 1, 'end': n, 'label': 'B',
     'fill_color': REGION_FILL_COLORS['B'], 'label_color':␣
↪REGION_LABEL_COLORS['B']},
] if locus.s >= 1 else []

# Create figure with GridSpec for the 3 matrices
fig = plt.figure(figsize=(14, 8))
gs = gridspec.GridSpec(1, 3, wspace=0.06)
axes = [fig.add_subplot(gs[i]) for i in range(3)]

# Plot using publication-quality plotter
plot_flex_matrices_publication(
    fig=fig,
    axes=axes,
    result=result,
    X=locus.X,
    Y=Y,
    s=locus.s,
    e=locus.e,
    regions=regions,
    colormap=grid_color_map,
    title_pad=18,
    nuc_label_fontsize=8,
)

title = (f"NW-flex STR mode\n"
         f"$X = A·R^{{{locus.N}}}·B, Y = A·Z*·B$\n"
         f"$Z^* = R(a,b,M) = ({a},{b},{M})$")
fig.suptitle(title, y=1.05)
plt.show()
```

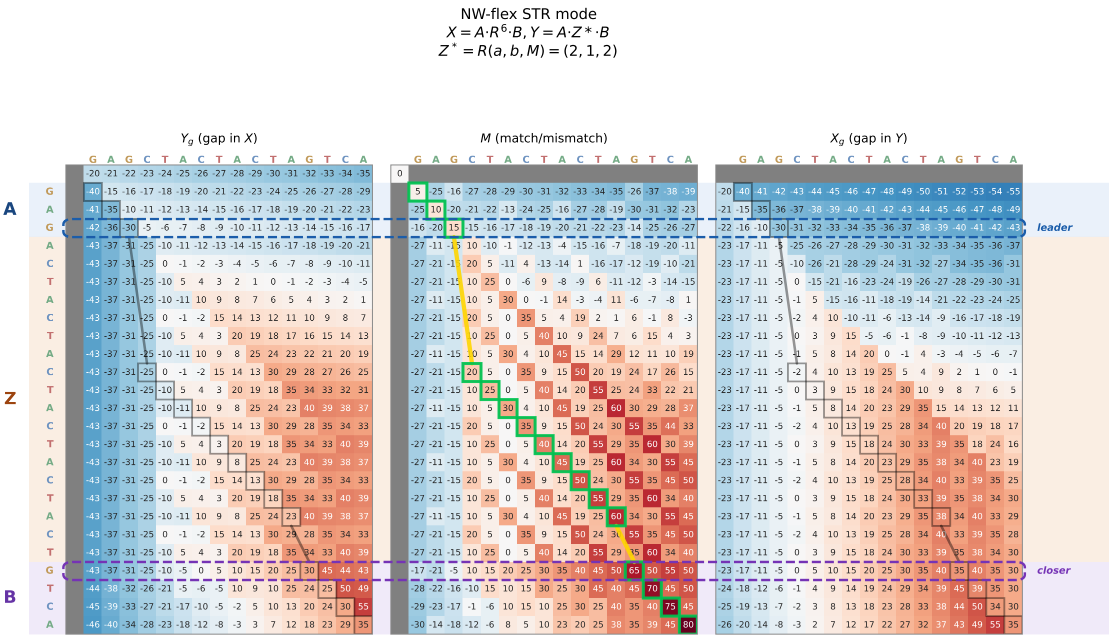

#### 4.7 Scalar plateau of $S_{\text{flex}}(X_N, Y)$ vs $N$

A key property of the STR EP pattern is that the flex score $S_{\text{flex}}(X_N, Y)$ is **non-decreasing** in the reference copy number $N$ and **plateaus** once $X_N = A \cdot R^N \cdot B$ contains enough copies to optimally align the read.

For a read $Y = A \cdot Z^* \cdot B$ where $Z^* = \text{phase}(R, a, b, M)$, the "repeat content" is

$$\text{repeat content} = M + \mathbf{1}_{\{a>0\}} + \mathbf{1}_{\{b>0\}}.$$

The flex score plateaus precisely when $N$ reaches this value: the reference then contains exactly enough repeat material to accommodate $Z^*$ with a perfect match.

We demonstrate this by plotting the flex score for **four different reads**, each with a different phase combination $(a, b, M)$:

| Case | $(a, b, M)$ | Repeat content | Description |
|---|---|---|---|
| 1 | $(0, 0, 3)$ | 3 | No partial repeats |
| 2 | $(1, 0, 2)$ | 3 | Left partial only |
| 3 | $(0, 2, 2)$ | 3 | Right partial only |
| 4 | $(2, 1, 2)$ | 4 | Both partials |

```
def str_flex_score_for_N(N_test: int, A: str, R: str, B: str, Y_fixed: str) -> float:
    """Compute S_flex(X_N, Y) for X_N = A · R^N · B."""
    X_N = A + R * N_test + B
```

```python
    s_N = len(A)
    e_N = len(A) + len(R) * N_test
    k_N = len(R)

    EP_N = build_EP_STR_phase(len(X_N), s=s_N, e=e_N, k=k_N)
    result_N = align_with_EP(
        X=X_N,
        Y=Y_fixed,
        score_matrix=score_matrix,
        gap_open=gap_open,
        gap_extend=gap_extend,
        alphabet_to_index=alphabet_to_index,
        extra_predecessors=EP_N,
        return_data=False,
    )
    return result_N.score

# Define four test cases with different (a, b, M) combinations
test_cases = [
    (0, 0, 3, "$(0,0,3)$: no partials"),       # repeat content = 3
    (1, 0, 2, "$(1,0,2)$: left partial"),      # repeat content = 3
    (0, 2, 2, "$(0,2,2)$: right partial"),     # repeat content = 3
    (2, 1, 2, "$(2,1,2)$: both partials"),     # repeat content = 4
]

# Vary N from 1 to 10
N_values = list(range(1, 11))

# Plot all cases on the same figure
plt.figure(figsize=(9, 5))
colors = ['#1f77b4', '#ff7f0e', '#2ca02c', '#d62728']
markers = ['o', 's', '^', 'D']

for idx, (a_test, b_test, M_test, label) in enumerate(test_cases):
    # Build Y for this (a, b, M)
    Y_test = locus.build_locus_variant(a_test, b_test, M_test)

    # Compute repeat content (plateau point)
    overhead = (1 if a_test > 0 else 0) + (1 if b_test > 0 else 0)
    repeat_content = M_test + overhead

    # Compute scores for each N
    scores = [
        str_flex_score_for_N(N_i, locus.A, locus.R, locus.B, Y_test)
        for N_i in N_values
    ]
```

```python
    # Plot curve
    plt.plot(
        N_values, scores, marker=markers[idx], linewidth=2, markersize=7,
        color=colors[idx], label=f"{label}, plateau at N={repeat_content}"
    )

    # Add vertical line at plateau point
    plt.axvline(
        x=repeat_content, color=colors[idx],
        linestyle=':', alpha=0.5, linewidth=1.5
    )

plt.xlabel("$N$ (copy number in $X_N = A \\cdot R^N \\cdot B$)", fontsize=11)
plt.ylabel("$S_{\\mathrm{flex}}(X_N, Y)$", fontsize=11)
title = "Monotone plateau: score stabilizes at " \
        "$N = M + \\mathbf{1}_{a>0} + \\mathbf{1}_{b>0}$"
plt.title(title, fontsize=12)
plt.legend(loc='lower right', fontsize=9)
plt.grid(True, alpha=0.3)
plt.xticks(N_values)
plt.tight_layout()
plt.show()
```

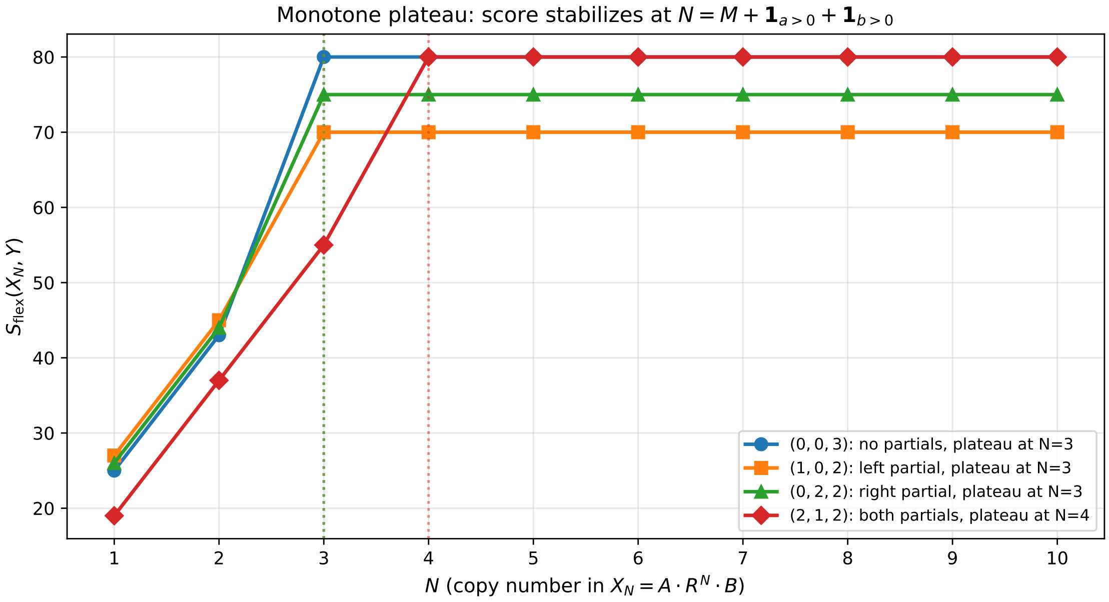

Each curve reaches its maximum (plateau) exactly when $N = M + \mathbf{1}_{\{a>0\}} + \mathbf{1}_{\{b>0\}}$, as indicated by the dotted vertical lines. The three cases with repeat content 3 all plateau at $N = 3$, while case 4 (with both partial repeats) plateaus at $N = 4$.

This confirms that the STR EP pattern correctly identifies the minimum reference length needed to optimally align any phased read.

#### 4.8 Phase-wise monotonicity in the DP grid

The scalar plateau we observed above is a consequence of a finer-grained property: **phase-wise monotonicity** in each layer of the Gotoh DP.

Inside the STR block $Z = R^N$ (rows $s+1, \ldots, e$), we group rows by their **motif phase**:

- For a given phase offset $r \in \{0, 1, \ldots, k-1\}$, consider the row indices

$$i_n = s + 1 + r + n \cdot k$$

  such that $s + 1 \leq i_n \leq e$.

For a fixed column $j$ and **each Gotoh layer** ($Y_g$, $M$, $X_g$), the sequence of scores along these phase-equivalent rows is **non-decreasing** in $n$:

$$L(i_0, j) \leq L(i_1, j) \leq L(i_2, j) \leq \cdots$$

where $L \in \{Y_g, M, X_g\}$.

Intuitively: once the repeat block is long enough to represent a good partial match, extending it by whole-copy increments of $R$ cannot reduce the score. The scores rise and then plateau, mirroring the scalar behavior we saw earlier.

```
# Get DP data from the alignment result
data = result.data
n_rows, n_cols = data.M.shape
print(f"DP grid shape: {n_rows} rows × {n_cols} columns")

# Helper to get rows for a given phase r inside Z
def phase_rows(s: int, e: int, k: int, r: int):
    """Get row indices inside Z with phase r."""
    rows = []
    i = s + 1 + r
    while i <= e:
        rows.append(i)
        i += k
    return rows

# Check monotonicity for all phases, columns, and layers
layers = [('Yg', data.Yg), ('M', data.M), ('Xg', data.Xg)]
violations = []

for layer_name, layer_mat in layers:
    for r in range(k):
        rows_r = phase_rows(s, e, k, r)
        if len(rows_r) <= 1:
            continue
        for j in range(1, n_cols):
```

```
            vals = [layer_mat[i, j] for i in rows_r]
            for idx in range(len(vals) - 1):
                if vals[idx + 1] < vals[idx]:
                    violations.append((layer_name, r, j, rows_r, vals))
                    break

if not violations:
    print("[OK] All phase-wise sequences are non-decreasing in Yg, M, and Xg.")
else:
    print(f"[FAIL] Found {len(violations)} violations:")
    for (layer, r, j, rows_r, vals) in violations[:5]:
        print(f"  {layer}: phase r={r}, col j={j}, values={vals}")
```

```
DP grid shape: 26 rows × 17 columns
[OK] All phase-wise sequences are non-decreasing in Yg, M, and Xg.
```

##### 4.8.1 Visualizing phase-wise monotonicity

We plot the score sequences along rows $i = s + 1 + r + n \cdot k$ for each phase $r$ and each Gotoh layer ($Y_g$, $M$, $X_g$). Each curve shows how the score changes as we add more repeat copies ($n$ increases).

Non-decreasing curves rising to a plateau confirm the phase-wise monotonicity property—the same pattern we saw in the scalar flex score.

```
def plot_phase_monotonicity_lines(data, s, e, k, Y, figsize=None):
    """
    Plot score sequences along rows i = (s+1) + r + n*k for each phase r
    and Gotoh layer (Yg, M, Xg).

    Each curve shows how the score changes as we add more repeat copies (n).
    Non-decreasing curves rising to a plateau confirm phase-wise monotonicity.
    """
    Yg, M_mat, Xg = data.Yg, data.M, data.Xg
    n_rows, n_cols = Yg.shape

    # Show columns based on Y length (the repeat region of Y)
    cols_to_show = min(n_cols, len(Y) + 1)

    # Layers to plot
    layers = [
        ('$Y_g$ (Y-gap)', Yg, '#1f77b4'),
        ('$M$ (match)', M_mat, '#2ca02c'),
        ('$X_g$ (X-gap)', Xg, '#ff7f0e'),
    ]

    # Get phases with multiple rows
    def get_phase_rows(r):
        rows = []
```

```
        i = s + 1 + r
        while i <= e:
            rows.append(i)
            i += k
        return rows

    phases_with_rows = [(r, get_phase_rows(r)) for r in range(k)]
    phases_with_rows = [(r, rows) for r, rows in phases_with_rows if len(rows) >
↪1]

    n_phases = len(phases_with_rows)
    n_layers = len(layers)

    if figsize is None:
        figsize = (4.5 * n_phases, 3 * n_layers)

    fig, axes = plt.subplots(n_layers, n_phases, figsize=figsize, sharex='col')

    # Handle edge cases
    if n_phases == 1:
        axes = axes.reshape(-1, 1)
    if n_layers == 1:
        axes = axes.reshape(1, -1)

    # Use plasma colormap for column colors
    cmap = plt.cm.RdYlBu
    col_colors = [
        cmap(j / max(1, cols_to_show - 2))
        for j in range(1, cols_to_show)
    ]

    for layer_idx, (layer_name, layer_mat, layer_color) in enumerate(layers):
        for phase_idx, (r, rows_r) in enumerate(phases_with_rows):
            ax = axes[layer_idx, phase_idx]

            # n values (repeat copy index)
            n_vals = list(range(len(rows_r)))

            # Plot a line for each column j
            for col_idx, j in enumerate(range(1, cols_to_show)):
                scores = [layer_mat[i, j] for i in rows_r]

                label = f"j={j}" if col_idx < 8 else None
                ax.plot(
                    n_vals, scores,
                    marker='o', markersize=5,
                    linestyle='-', linewidth=1.8,
```

```
                    color=col_colors[col_idx],
                    label=label,
                    alpha=0.85,
                )

            # Styling
            if layer_idx == 0:
                ax.set_title(f"Phase $r = {r}$", fontsize=11,␣
↪fontweight='semibold')
            if layer_idx == n_layers - 1:
                ax.set_xlabel("Repeat index $n$", fontsize=10)
            if phase_idx == 0:
                ax.set_ylabel(f"{layer_name}", fontsize=10, color=layer_color)

            ax.grid(True, alpha=0.3)
            ax.set_xticks(n_vals)

            # Add row index annotations on bottom row
            if layer_idx == n_layers - 1:
                labels = [f"{n}\n(i={rows_r[n]})" for n in n_vals]
                ax.set_xticklabels(labels, fontsize=8)
            else:
                ax.set_xticklabels([])

    # Add shared legend at bottom
    handles, labels = axes[0, 0].get_legend_handles_labels()
    fig.legend(
        handles, labels,
        loc='upper center',
        bbox_to_anchor=(0.5, 0.02),
        ncol=min(10, len(handles)),
        fontsize=9,
        title="Column $j$ (position in Y)",
        title_fontsize=10,
    )

    fig.suptitle(
        f"Phase-wise monotonicity: scores rise to plateau (k={k})",
        fontsize=12, fontweight='semibold',
    )

    plt.tight_layout(rect=[0, 0.06, 1, 0.95])

    return fig

# Plot the phase monotonicity lines
fig = plot_phase_monotonicity_lines(data, s, e, k, Y)
```

```
plt.show()
```

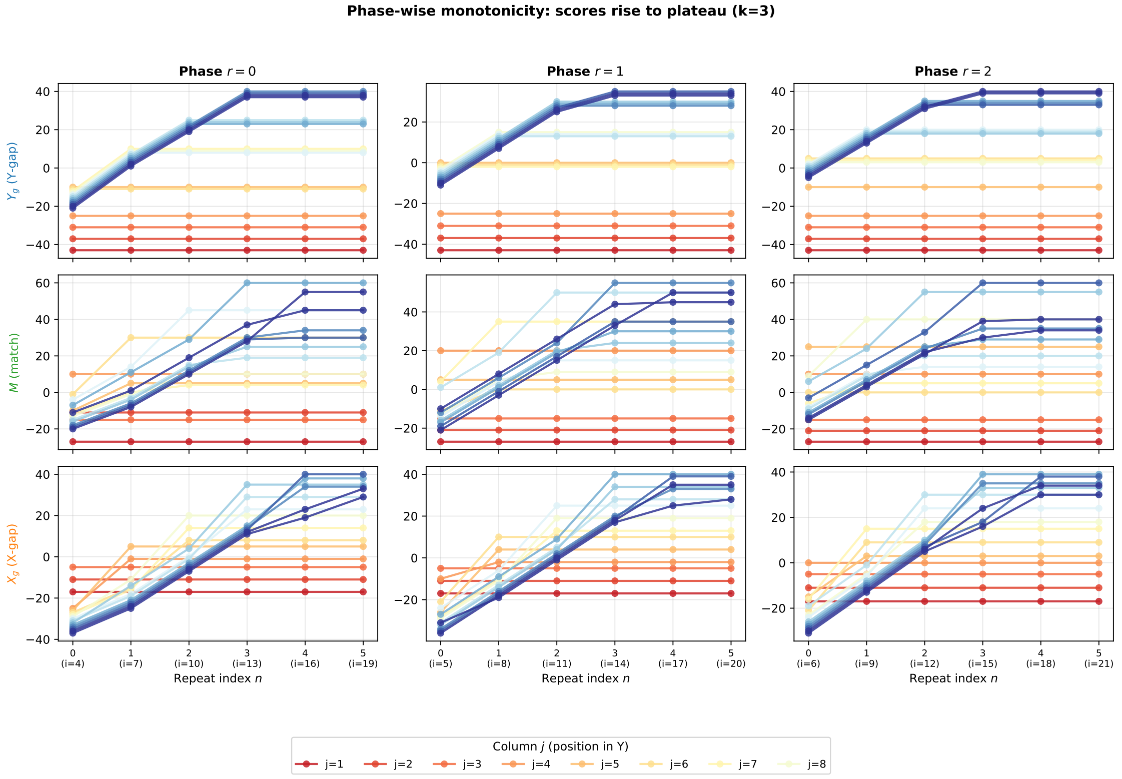

The rising-plateau curves in each panel mirror the scalar flex score behavior: as we move deeper into the repeat block along the same motif phase, the DP can always "reuse" or extend a good partial match without losing score.

This phase-wise monotonicity in the DP grid is the mechanism underlying the scalar plateau: once the reference contains enough copies, further extension cannot improve the alignment.

#### 4.9 Multi-block STR example (compound repeats)

Real STR loci sometimes contain multiple adjacent repeat blocks with different motifs. A **compound STR** has the form:

$$X = A \cdot R_1^{N_1} \cdot R_2^{N_2} \cdot B$$

where $R_1$ and $R_2$ are different motifs (possibly with different lengths).

The `build_EP_multi_STR_phase()` function constructs an EP pattern that handles multiple STR blocks by taking the union of the individual block patterns. Each block has its own leader row, closer row, and phase-preserving exit condition.

We demonstrate this with a compound locus containing two blocks: - Block 1: $R_1 =$ `AC` with $N_1 = 4$ - Block 2: $R_2 =$ `TGG` with $N_2 = 3$

```python
# Define a compound STR locus with two blocks
compound = CompoundSTRLocus(
    A="GAT",
    blocks=[("AC", 4), ("TGG", 3)],  # (R1, N1), (R2, N2)
    B="CTA",
)

print(f"Compound locus: X = A · R1^N1 · R2^N2 · B")
print(f"  A = '{compound.A}'")
print(f"  R1 = 'AC', N1 = 4")
print(f"  R2 = 'TGG', N2 = 3")
print(f"  B = '{compound.B}'")
print()
print(f"Reference X: {compound.X} (length {compound.n})")
print()

# Get block boundaries
boundaries = compound.block_boundaries()
print("Block boundaries (s, e, k):")
for i, (s_i, e_i, k_i) in enumerate(boundaries):
    print(f"  Block {i+1}: s={s_i}, e={e_i}, k={k_i}")
```

```
Compound locus: X = A · R1^N1 · R2^N2 · B
  A = 'GAT'
  R1 = 'AC', N1 = 4
  R2 = 'TGG', N2 = 3
  B = 'CTA'

Reference X: GATACACACACTGGTGGTGGCTA (length 23)

Block boundaries (s, e, k):
  Block 1: s=3, e=11, k=2
  Block 2: s=11, e=20, k=3
```

```python
# Build the multi-block EP pattern
n_compound = compound.n
EP_multi = build_EP_multi_STR_phase(n_compound, boundaries)

# Create a read with partial content from both blocks
# Block 1: use (a1=1, b1=0, M1=2) -> suf(AC,1)·AC^2 = "C" + "ACAC" = "CACAC"
# Block 2: use (a2=0, b2=2, M2=1) -> TGG^1·pre(TGG,2) = "TGG" + "TG" = "TGGTG"
R1, N1 = compound.blocks[0]
R2, N2 = compound.blocks[1]

Z1_star = phase_repeat(R1, a=1, b=0, M=2)  # "CACAC"
Z2_star = phase_repeat(R2, a=0, b=2, M=1)  # "TGGTG"
```

```python
Y_compound = compound.A + Z1_star + Z2_star + compound.B

print(f"Read Y with partial content from both blocks:")
print(f"  Z1* = phase('{R1}', 1, 0, 2) = '{Z1_star}'")
print(f"  Z2* = phase('{R2}', 0, 2, 1) = '{Z2_star}'")
print(f"  Y = A + Z1* + Z2* + B = '{Y_compound}'")
print(f"  |Y| = {len(Y_compound)}")
```

```
Read Y with partial content from both blocks:
  Z1* = phase('AC', 1, 0, 2) = 'CACAC'
  Z2* = phase('TGG', 0, 2, 1) = 'TGGTG'
  Y = A + Z1* + Z2* + B = 'GATCACACTGGTGCTA'
  |Y| = 16
```

```python
# Run alignment
result_compound = align_with_EP(
    X=compound.X,
    Y=Y_compound,
    score_matrix=score_matrix,
    gap_open=gap_open,
    gap_extend=gap_extend,
    alphabet_to_index=alphabet_to_index,
    extra_predecessors=EP_multi,
    return_data=True,
)

# Expected score: perfect match
expected_compound = match_score * len(Y_compound)

print(f"Alignment score: {result_compound.score}")
print(f"Expected score:  {expected_compound}")
print(f"Match: {'[OK]' if result_compound.score == expected_compound else␣
 ↪'[FAIL]'}")
print()
print(f"Aligned X: {result_compound.X_aln}")
print(f"Aligned Y: {result_compound.Y_aln}")
print()
print("Row jumps:", *result_compound.jumps, sep="\n")
```

```
Alignment score: 80.0
Expected score:  80.0
Match: [OK]

Aligned X: GATCACACTGGTGCTA
Aligned Y: GATCACACTGGTGCTA

Row jumps:
RowJump(from_row=3, to_row=7, col=4, state=1)
```

```
RowJump(from_row=11, to_row=15, col=9, state=1)
RowJump(from_row=19, to_row=21, col=14, state=1)
```

##### 4.9.1 Visualizing the multi-block alignment

We plot the DP matrices for the compound locus. The two repeat blocks are highlighted separately, showing how the alignment jumps through each block independently.

```
# For multi-block, we use the first block's boundaries for the main visualization
s1, e1, k1 = boundaries[0]
s2, e2, k2 = boundaries[1]
n = len(compound.X)

# Build regions for compound STR (A, R1 block, R2 block, B)
regions = [
    {'start': 1, 'end': s1, 'label': 'A',
     'fill_color': REGION_FILL_COLORS['A'], 'label_color':␣
↪REGION_LABEL_COLORS['A']},
    {'start': s1 + 1, 'end': e1, 'label': 'R1',
     'fill_color': REGION_FILL_COLORS['Z'], 'label_color':␣
↪REGION_LABEL_COLORS['Z']},
    {'start': e1 + 1, 'end': e2, 'label': 'R2',
     'fill_color': '#E8D5B7', 'label_color': '#8B7355'},  # Tan for R2
    {'start': e2 + 1, 'end': n, 'label': 'B',
     'fill_color': REGION_FILL_COLORS['B'], 'label_color':␣
↪REGION_LABEL_COLORS['B']},
] if s1 >= 1 else []

# Extract R1* and R2* boundaries from jumps
# Multi-block compound STR has 3 jumps:
#   Jump 0: entry into R1 (from leader s1)
#   Jump 1: exit R1 / entry R2 (from R1 to R2)
#   Jump 2: exit R2 (to closer)
# Use cyan and yellow - highly visible against red background
BROWN = '#8B4513'      # Light Brown - visible against red
YELLOW = '#FFD700'     # Gold/yellow - visible against red
row_highlights = []

if len(result_compound.jumps) >= 3:
    # R1* boundaries: entry is jump[0].to_row, exit is jump[1].from_row
    r1_star_start = result_compound.jumps[0].to_row
    r1_star_end = result_compound.jumps[1].from_row

    # R2* boundaries: entry is jump[1].to_row, exit is jump[2].from_row
    r2_star_start = result_compound.jumps[1].to_row
    r2_star_end = result_compound.jumps[2].from_row
```

```python
    row_highlights = [
        {'start': r1_star_start, 'end': r1_star_end, 'label': 'R1*',
         'color': YELLOW, 'linestyle': '--', 'label_position': 'top'},
        {'start': r2_star_start, 'end': r2_star_end, 'label': 'R2*',
         'color': BROWN, 'linestyle': '--', 'label_position': 'top'},
    ]

# Create figure with GridSpec for the 3 matrices
fig = plt.figure(figsize=(16, 8))
gs = gridspec.GridSpec(1, 3, wspace=0.06)
axes = [fig.add_subplot(gs[i]) for i in range(3)]

# Plot using publication-quality plotter
plot_flex_matrices_publication(
    fig=fig,
    axes=axes,
    result=result_compound,
    X=compound.X,
    Y=Y_compound,
    s=s1,  # Use first block's leader
    e=e2,  # Use second block's end
    regions=regions,
    row_highlights=row_highlights,
    colormap=grid_color_map,
    title_pad=18,
    nuc_label_fontsize=7,
)

fig.suptitle(f"Multi-block STR alignment: X = A·R1^{{{N1}}}·R2^{{{N2}}}·B", y=1.02)
plt.show()
```

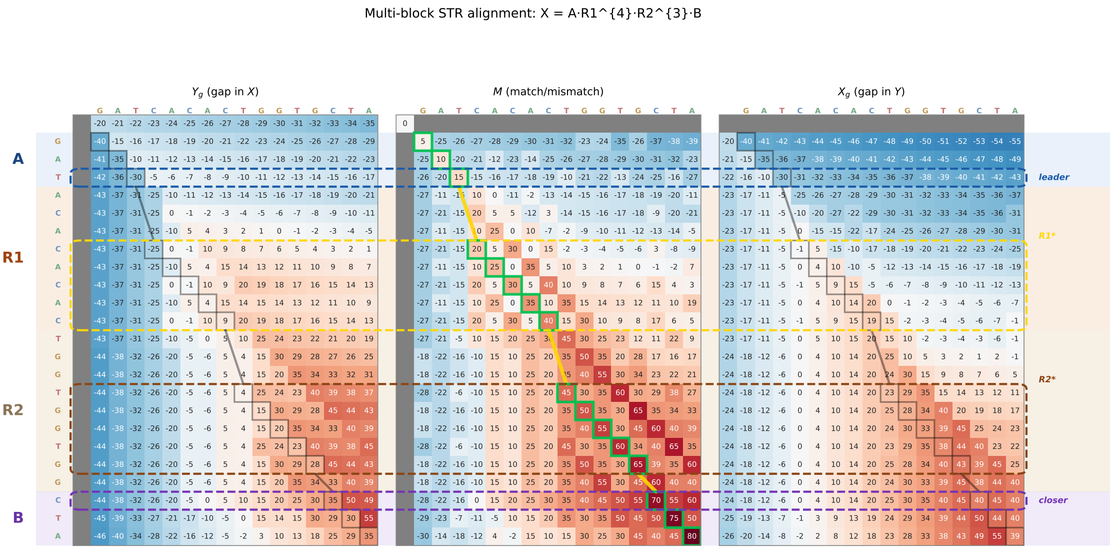

---

#### 4.10 Summary

In this notebook, we developed the **STR specialization** of the NW-Flex algorithm, demonstrating how the general framework from Notebooks 02-03 applies to Short Tandem Repeat analysis.

##### 4.10.1 Key Concepts

1. **STR Locus Structure**: An STR locus has the form $X = A \cdot R^N \cdot B$, where:
   - $A$ = left flanking sequence (leader)
   - $R$ = repeat unit of length $k$
   - $N$ = number of repeat copies
   - $B$ = right flanking sequence (trailer)
2. **Phased Query Sequences**: Valid query sequences for STR loci have the form:

   $$Z^* = \text{suf}(R, a) \cdot R^M \cdot \text{pre}(R, b)$$

   where $a, b \in \{0, 1, \ldots, k-1\}$ define the phase and $M$ is the number of complete repeats.
3. **The Fundamental Constraint**: The $(a, b, M)$ triplet must satisfy:

   $$M + \mathbf{1}_{a>0} + \mathbf{1}_{b>0} \leq N$$

   This ensures the query can be accommodated within the reference.
4. **EP Pattern for STRs**: The `build_EP_STR_phase` function constructs the equal-penalty pattern that:
   - Allows entry from the leader row $s$ into any row of the repeat block
   - Allows exit only from phase-compatible rows $\{s, e-k+1, \ldots, e\}$
   - Encodes the phase information $(a, b)$ in the entry/exit points

##### 4.10.2 Validation Results

We verified that **all valid** $(a, b, M)$ **combinations** produce:

- Perfect alignment scores (equal to query length × match bonus)
- Correct phase inference from jump analysis
- Proper constraint satisfaction

##### 4.10.3 Monotonicity Properties

1. **Scalar plateau**: The flex score $S_{\text{flex}}(X_N, Y)$ is non-decreasing in $N$ and plateaus exactly at

   $$N = M + \mathbf{1}_{a>0} + \mathbf{1}_{b>0}$$

   the minimum reference copy number needed to optimally align the read.
2. **Phase-wise monotonicity**: For fixed column $j$ and phase $r$, the DP scores $F(i, j)$ along rows $i = s + 1 + r + nk$ are non-decreasing in $n$.

###### 4.10.4 Multi-block Extension

The framework extends naturally to **compound STR loci** with multiple repeat blocks:

$$X = A \cdot R_1^{N_1} \cdot R_2^{N_2} \cdot B$$

Using `CompoundSTRLocus` and `build_EP_multi_STR_phase`, each block maintains its own phase parameters while sharing a common alignment coordinate system.

---

###### 4.10.5 Next Steps

- **Notebook 05**: Explores the C++ accelerated implementation for production-scale analysis
- **Application**: Apply STR-EP to real genomic data with biological repeat units

### 5 Notebook 5: Cython-backed NW-flex — correctness and speed

This notebook demonstrates the Cython implementation of the NW-flex DP core and verifies that it produces identical results to the pure Python version while achieving substantial speedups on larger inputs.

#### 5.1 Overview

The pure Python implementation in `dp_core.py` is designed for clarity and easy modification. For performance-sensitive applications, we provide a Cython-backed implementation in `fast.py` that:

1. **Encodes sequences** as integer arrays for fast indexing.
2. **Converts EP patterns** to a compact interval representation for efficient iteration in tight loops.
3. **Runs the DP core in C** via Cython, with typed memory views and disabled bounds checking.
4. **Reuses the Python traceback** to recover aligned sequences and row jumps.

Both implementations share the same API and produce identical results:

| Function | Module | Use case |
|---|---|---|
| `run_flex_dp` | `dp_core.py` | Teaching, debugging, modification |
| `run_flex_dp_fast` | `fast.py` | Production, benchmarking |

This notebook is organized as follows:

1. **Setup and imports** — load modules and scoring.
2. **EP interval encoding** — explain the compact representation used by Cython.
3. **Sanity check: Python vs Cython** — verify identical scores, alignments, and DP matrices on a small example.
4. **Benchmarking** — compare run times across sequence lengths for both standard NW and single-block NW-flex.
5. **Summary** — key takeaways and usage guidance.

#### 5.2 Setup and imports

```
#  Notebook magic: autoreload modules
%load_ext autoreload
%autoreload 2

import numpy as np
import pandas as pd
import matplotlib.pyplot as plt

# Core NW-flex functions
from nwflex.dp_core import FlexInput, run_flex_dp
from nwflex.fast import run_flex_dp_fast

# EP pattern builders
from nwflex.ep_patterns import build_EP_standard, build_EP_single_block
from nwflex.ep_intervals import ep_to_intervals

# Scoring and alignment helpers
from nwflex.validation import get_default_scoring
from nwflex.dp_core import AlignmentResult

# Plotting
from nwflex.plot import plot_flex_matrices, plot_score_system

# Import benchmarking utilities from validation module
from nwflex.validation import benchmark_python_vs_cython

## random number generator
rng = np.random.default_rng(888)

## validation parameters
lengths = [50, 100, 200, 400, 800]
n_samples = 10
cython_multiplier = 100
```

```
The autoreload extension is already loaded. To reload it, use:
  %reload_ext autoreload
```

##### 5.2.1 Scoring scheme

We use the same default scoring as in the other notebooks: match = +5, mismatch = -5, gap-open = -20, gap-extend = -1.

```
score_matrix, gap_open, gap_extend, alphabet_to_index = get_default_scoring()

fig = plot_score_system(score_matrix, gap_open, gap_extend, alphabet_to_index)
plt.show()
```

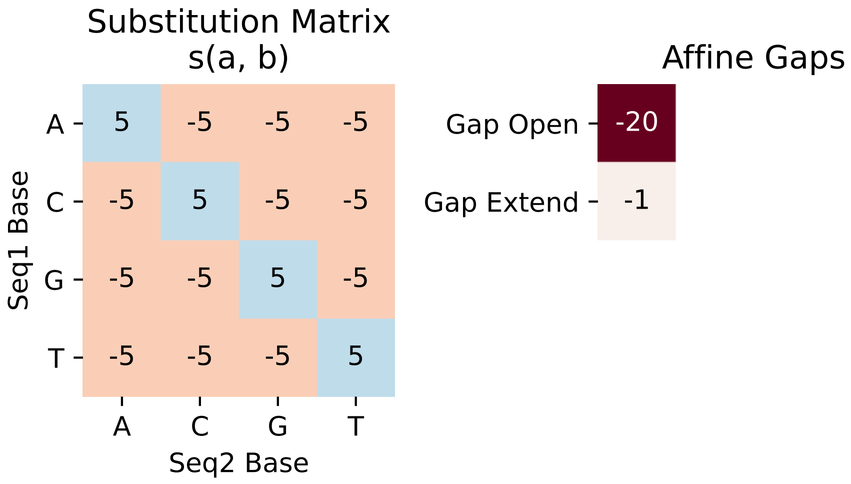

##### 5.3 EP interval encoding

The NW-flex DP uses **extra predecessor sets** $E(i)$ to augment the baseline Gotoh recurrence. In Python, these are naturally represented as a list of lists:

```
EP[i] = [r0, r1, ...]  # predecessor rows for DP row i
```

For the Cython core, iterating over Python lists would be slow. Instead, we convert each $E(i)$ into **disjoint intervals** $[a, b]$ covering the sorted predecessor rows. This is handled by `ep_to_intervals`:

```
ep_counts, ep_starts, ep_ends = ep_to_intervals(EP)
```

where: - `ep_counts[i]` = number of intervals for row $i$ - `ep_starts[i, k]`, `ep_ends[i, k]` = bounds of the $k$-th interval

This representation is compact and allows the Cython inner loops to iterate over contiguous ranges without Python overhead.

###### 5.3.1 Example: single-block EP encoding

For a reference $X = A \cdot Z \cdot B$ with $|A| = 3$ and $|Z| = 4$ (so $s = 3$, $e = 7$), the EP pattern is:

- Rows $s + 1, \ldots, e$ (rows 4–7): $E(i) = \{s\} = \{3\}$
- Closer row $e + 1$ (row 8): $E(8) = \{3, 4, 5, 6, 7\}$

```
# Build single-block EP for n=10, s=3, e=7
n, s, e = 10, 3, 7
EP = build_EP_single_block(n, s, e)

print("EP list-of-lists representation:")
for i, ep_row in enumerate(EP):
    if ep_row:
        print(f"  E({i}) = {list(ep_row)}")
```

```python
# Convert to intervals
ep_counts, ep_starts, ep_ends = ep_to_intervals(EP)

print("\nInterval representation:")
for i in range(len(EP)):
    if ep_counts[i] > 0:
        intervals = [(ep_starts[i, k], ep_ends[i, k]) for k in␣
↪range(ep_counts[i])]
        print(f"  Row {i}: {ep_counts[i]} interval(s) -> {intervals}")
```

```
EP list-of-lists representation:
  E(4) = [3]
  E(5) = [3]
  E(6) = [3]
  E(7) = [3]
  E(8) = [3, 4, 5, 6]

Interval representation:
  Row 4: 1 interval(s) -> [(np.int32(3), np.int32(3))]
  Row 5: 1 interval(s) -> [(np.int32(3), np.int32(3))]
  Row 6: 1 interval(s) -> [(np.int32(3), np.int32(3))]
  Row 7: 1 interval(s) -> [(np.int32(3), np.int32(3))]
  Row 8: 1 interval(s) -> [(np.int32(3), np.int32(6))]
```

Notice that the closer row's EP set $\{3, 4, 5, 6, 7\}$ collapses into a single interval $[3, 7]$. This provides a simplified encoding of the EP patterns most common in practice. A more advanced encoding scheme could more efficiently handle leaders and closers, at the cost of added complexity.

##### 5.4 Sanity check: Python vs Cython on a small example

We verify that `run_flex_dp` (Python) and `run_flex_dp_fast` (Cython) produce identical results on a small example. We check:

1. **Scores** — the flex score $S_{\text{flex}}(X, Y)$
2. **Aligned sequences** — the gap-inserted strings
3. **Row jumps** — the EP-induced jumps recorded during traceback
4. **DP matrices** — the full $Y_g$, $M$, $X_g$ tables

We test both standard NW (EP = empty) and single-block configurations.

```python
X_small = "GGGATATGGG"
Y_small = "GGGATGGG"

print(f"X = {X_small}  (n = {len(X_small)})")
print(f"Y = {Y_small}  (m = {len(Y_small)})")
```

```
X = GGGATATGGG  (n = 10)
Y = GGGATGGG  (m = 8)
```

##### 5.4.1 Standard NW-flex (EP = empty)

With no extra predecessors, NW-flex reduces to standard Needleman–Wunsch/Gotoh.

```
# Build FlexInput for standard NW
n = len(X_small)
EP_std = build_EP_standard(n)

cfg_std = FlexInput(
    X=X_small,
    Y=Y_small,
    score_matrix=score_matrix,
    gap_open=gap_open,
    gap_extend=gap_extend,
    extra_predecessors=EP_std,
    alphabet_to_index=alphabet_to_index,
)

# Run both implementations with full data
py_result = run_flex_dp(cfg_std, return_data=True)
(score_py, X_aln_py, Y_aln_py,
 path_py, jumps_py, data_py) = py_result.to_tuple()
res_fast = run_flex_dp_fast(cfg_std, return_data=True)

print("Standard NW (EP = empty):")
print(f"  Python score:  {score_py}")
print(f"  Cython score:  {res_fast.score}")
print(f"  Scores match:  {score_py == res_fast.score}")
print()
print(f"  Python X_aln:  {X_aln_py}")
print(f"  Cython X_aln:  {res_fast.X_aln}")
print(f"  X_aln match:   {X_aln_py == res_fast.X_aln}")
print()
print(f"  Python Y_aln:  {Y_aln_py}")
print(f"  Cython Y_aln:  {res_fast.Y_aln}")
print(f"  Y_aln match:   {Y_aln_py == res_fast.Y_aln}")
print()
print(f"  Row jumps (Python): {jumps_py}")
print(f"  Row jumps (Cython): {res_fast.jumps}")
```

```
Standard NW (EP = empty):
  Python score:  19.0
  Cython score:  19.0
  Scores match:  True

  Python X_aln:  GGGATATGGG
  Cython X_aln:  GGGATATGGG
  X_aln match:   True
```

```
Python Y_aln:  GGG--ATGGG
Cython Y_aln:  GGG--ATGGG
Y_aln match:   True

Row jumps (Python): []
Row jumps (Cython): []
```

##### 5.4.2 Single-block NW-flex (A·Z·B)

Now we add a flexible block in the center of $X$. This exercises the EP refinement logic in both implementations.

```
# Single-block config: Z is roughly the middle 40% of X
n = len(X_small)
block_len = max(1, int(0.4 * n))
s = (n - block_len) // 2
e = s + block_len

print(f"Block configuration: s = {s}, e = {e}")
print(f"  A = X[:{s}] = '{X_small[:s]}'")
print(f"  Z = X[{s}:{e}] = '{X_small[s:e]}'")
print(f"  B = X[{e}:] = '{X_small[e:]}'")

EP_blk = build_EP_single_block(n, s, e)

cfg_blk = FlexInput(
    X=X_small,
    Y=Y_small,
    score_matrix=score_matrix,
    gap_open=gap_open,
    gap_extend=gap_extend,
    extra_predecessors=EP_blk,
    alphabet_to_index=alphabet_to_index,
)

# Run both implementations
py_result_blk = run_flex_dp(cfg_blk, return_data=True)
(score_py_blk, X_aln_py_blk, Y_aln_py_blk,
 path_py_blk, jumps_py_blk, data_py_blk) = py_result_blk.to_tuple()
res_fast_blk = run_flex_dp_fast(cfg_blk, return_data=True)

print("\nSingle-block NW-flex:")
print(f"  Python score:  {score_py_blk}")
print(f"  Cython score:  {res_fast_blk.score}")
print(f"  Scores match:  {score_py_blk == res_fast_blk.score}")
print()
print(f"  Python X_aln:  {X_aln_py_blk}")
```

```
print(f"  Cython X_aln:  {res_fast_blk.X_aln}")
print(f"  X_aln match:   {X_aln_py_blk == res_fast_blk.X_aln}")
print()
print(f"  Python Y_aln:  {Y_aln_py_blk}")
print(f"  Cython Y_aln:  {res_fast_blk.Y_aln}")
print(f"  Y_aln match:   {Y_aln_py_blk == res_fast_blk.Y_aln}")
print()
print(f"  Row jumps (Python): {jumps_py_blk}")
print(f"  Row jumps (Cython): {res_fast_blk.jumps}")
```

```
Block configuration: s = 3, e = 7
  A = X[:3] = 'GGG'
  Z = X[3:7] = 'ATAT'
  B = X[7:] = 'GGG'

Single-block NW-flex:
  Python score:  40.0
  Cython score:  40.0
  Scores match:  True

  Python X_aln:  GGGATGGG
  Cython X_aln:  GGGATGGG
  X_aln match:   True

  Python Y_aln:  GGGATGGG
  Cython Y_aln:  GGGATGGG
  Y_aln match:   True

  Row jumps (Python): [RowJump(from_row=3, to_row=6, col=4, state=1)]
  Row jumps (Cython): [RowJump(from_row=3, to_row=6, col=4, state=1)]
```

##### 5.4.3 DP matrix comparison

We verify that the full DP matrices ($Y_g$, $M$, $X_g$) are identical between Python and Cython. Even small floating-point discrepancies would indicate a bug in one implementation.

```
def compare_dp_matrices(data_py, data_fast, label: str):
    """
    Compare DP matrices from Python and Cython implementations.

    Note: The Python implementation uses -inf for unreachable cells,
    while Cython uses a large negative sentinel value (-1e300).
    We compare only the "reachable" cells (values > -1e30).
    """
    print(f"{label}:")

    THRESHOLD = -1e30  # Cells below this are considered "unreachable"
```

```python
    def compare_layer(py_arr, fast_arr):
        # Both implementations mark unreachable cells with very negative values
        py_reachable = py_arr > THRESHOLD
        fast_reachable = fast_arr > THRESHOLD

        # Check same cells are reachable
        if not np.array_equal(py_reachable, fast_reachable):
            return False

        # Compare reachable values
        mask = py_reachable
        if not mask.any():
            return True
        return np.allclose(py_arr[mask], fast_arr[mask])

    Yg_match = compare_layer(data_py.Yg, data_fast.Yg)
    M_match = compare_layer(data_py.M, data_fast.M)
    Xg_match = compare_layer(data_py.Xg, data_fast.Xg)

    print(f"  Yg matrices match: {Yg_match}")
    print(f"  M  matrices match: {M_match}")
    print(f"  Xg matrices match: {Xg_match}")

    if not (Yg_match and M_match and Xg_match):
        print("  WARNING: Matrix mismatch detected!")
    else:
        print("  All reachable DP cells are identical.")

compare_dp_matrices(data_py, res_fast.data, "Standard NW")
print()
compare_dp_matrices(data_py_blk, res_fast_blk.data, "Single-block NW-flex")
```

```
Standard NW:
  Yg matrices match: True
  M  matrices match: True
  Xg matrices match: True
  All reachable DP cells are identical.

Single-block NW-flex:
  Yg matrices match: True
  M  matrices match: True
  Xg matrices match: True
  All reachable DP cells are identical.
```

##### 5.4.4 Visualizing the DP matrices and path

As a visual sanity check, we plot the three DP layers ($Y_g$, $M$, $X_g$) with the optimal path overlaid for both implementations. The Cython implementation uses a sentinel value (`-1e300`) instead of

-inf for unreachable cells, so we convert these before plotting.

```
# Wrap Python results into an AlignmentResult for plotting
result_py_blk = AlignmentResult(
    score=score_py_blk,
    X_aln=X_aln_py_blk,
    Y_aln=Y_aln_py_blk,
    path=path_py_blk,
    jumps=jumps_py_blk,
    data=data_py_blk,
)

# Convert Cython sentinel values (-1e300) to -inf for consistent plotting
CYTHON_SENTINEL = -1e30  # Cython uses -1e300, anything below this is␣
↪"unreachable"

def normalize_cython_data(data):
    """Convert Cython sentinel values to -inf for consistent representation."""
    for arr in [data.Yg, data.M, data.Xg]:
        arr[arr < CYTHON_SENTINEL] = -np.inf

# Normalize Cython data before plotting
normalize_cython_data(res_fast_blk.data)

# Plot Python result using standard plotting
fig = plot_flex_matrices(
    result=result_py_blk,
    X=X_small,
    Y=Y_small,
    s=s,
    e=e,
    marker_size=16,
    figsize=(14, 5),
)
fig.suptitle("Single-block NW-flex: Python implementation", fontsize=14)
fig.tight_layout()
plt.show()

# Plot Cython result using standard plotting
fig = plot_flex_matrices(
    result=res_fast_blk,
    X=X_small,
    Y=Y_small,
    s=s,
    e=e,
    marker_size=16,
    figsize=(14, 5),
```

```
)
fig.suptitle("Single-block NW-flex: Cython implementation", fontsize=14)
fig.tight_layout()
plt.show()
```

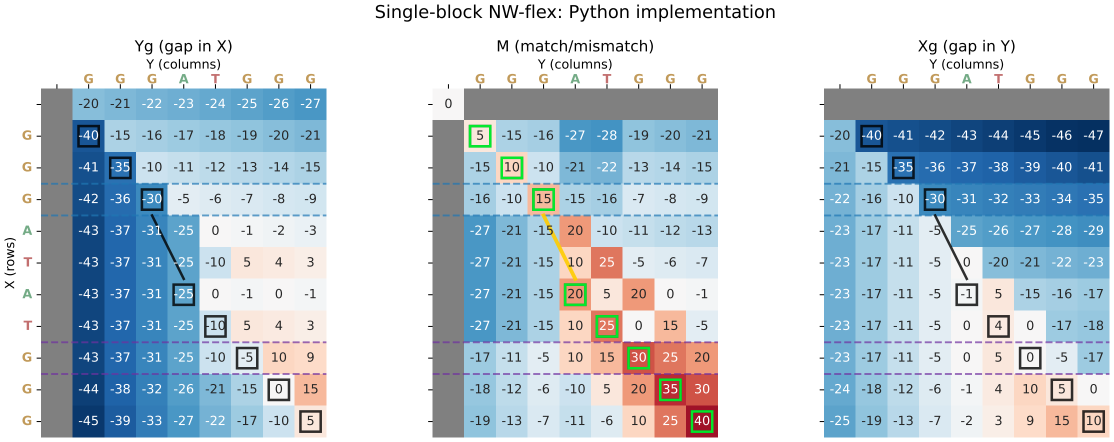

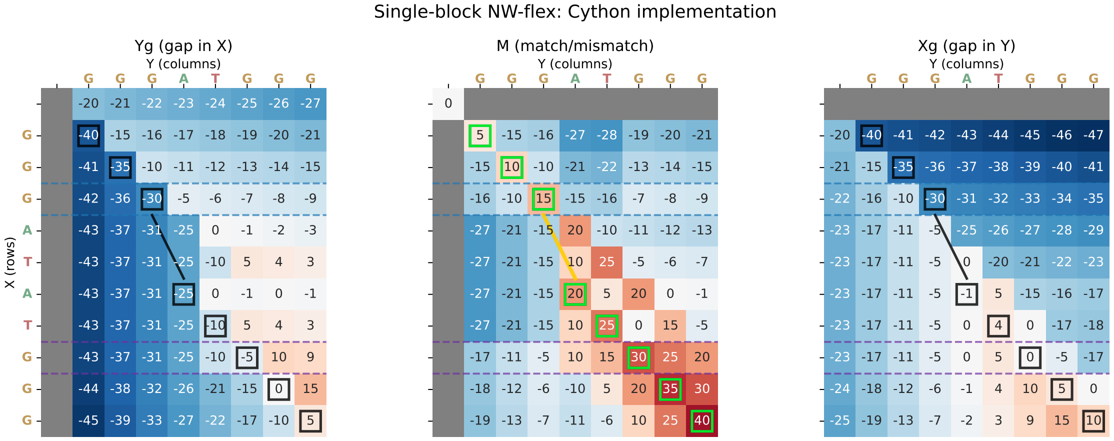

#### 5.5 Benchmarking: Python vs Cython

Now we compare run times on larger random sequences. We benchmark both:

1. **Standard NW-flex** (EP = empty) — baseline Needleman–Wunsch/Gotoh
2. **Single-block NW-flex** — with a flexible block in the center

Both have $O(|X| \cdot |Y|)$ time complexity; the Cython version should have smaller constants due to:

- Typed arrays and memory views (no Python object overhead)
- Disabled bounds checking (`boundscheck=False`)
- C-level loop iteration (no Python bytecode dispatch)
- Interval-based EP iteration (fewer loop iterations)

##### 5.5.1 Benchmark results

We test sequence lengths from 50 to 800, keeping $|X| = |Y|$ for simplicity. We use `timeit.Timer` for accurate averaging, running Cython 100× more iterations than Python to get reliable sub-millisecond measurements.

```
# Collect benchmark data
results_std = {}  # length -> (py_avg, cy_avg)
results_blk = {}

# Run standard NW-flex benchmarks
for length in lengths:
    py_avg, cy_avg = benchmark_python_vs_cython(
        length, length, rng, mode="standard",
        n_samples=n_samples, cython_multiplier=cython_multiplier
    )
    results_std[length] = (py_avg, cy_avg)

# Print standard results table
print("Standard NW-flex (EP = empty):")
print("-" * 70)
for length in lengths:
    py_avg, cy_avg = results_std[length]
    speedup = py_avg / cy_avg
    print(f"len = {length:4d}  "
          f"Python: {py_avg:.4f}s ({n_samples} runs)  "
          f"Cython: {cy_avg:.6f}s ({n_samples * cython_multiplier} runs)  "
          f"speedup: {speedup:.1f}x")

# Run single-block NW-flex benchmarks
for length in lengths:
    py_avg, cy_avg = benchmark_python_vs_cython(
        length, length, rng, mode="single_block",
        n_samples=n_samples, cython_multiplier=cython_multiplier
    )
    results_blk[length] = (py_avg, cy_avg)

# Print single-block results table
print("\n\nSingle-block NW-flex (A·Z·B):")
print("-" * 70)
for length in lengths:
    py_avg, cy_avg = results_blk[length]
    speedup = py_avg / cy_avg
    print(f"len = {length:4d}  "
          f"Python: {py_avg:.4f}s ({n_samples} runs)  "
          f"Cython: {cy_avg:.6f}s ({n_samples * cython_multiplier} runs)  "
          f"speedup: {speedup:.1f}x")
```

```
Standard NW-flex (EP = empty):
----------------------------------------------------------------------
len =   50  Python: 0.0123s (10 runs)  Cython: 0.000049s (1000 runs)  speedup:
249.6x
len =  100  Python: 0.0491s (10 runs)  Cython: 0.000126s (1000 runs)  speedup:
390.6x
len =  200  Python: 0.1956s (10 runs)  Cython: 0.000376s (1000 runs)  speedup:
520.0x
len =  400  Python: 0.8056s (10 runs)  Cython: 0.001894s (1000 runs)  speedup:
425.3x
len =  800  Python: 3.1867s (10 runs)  Cython: 0.007308s (1000 runs)  speedup:
436.0x

Single-block NW-flex (A·Z·B):
----------------------------------------------------------------------
len =   50  Python: 0.0191s (10 runs)  Cython: 0.000066s (1000 runs)  speedup:
287.9x
len =  100  Python: 0.0781s (10 runs)  Cython: 0.000168s (1000 runs)  speedup:
464.8x
len =  200  Python: 0.2848s (10 runs)  Cython: 0.000715s (1000 runs)  speedup:
398.5x
len =  400  Python: 1.1568s (10 runs)  Cython: 0.002762s (1000 runs)  speedup:
418.8x
len =  800  Python: 4.6050s (10 runs)  Cython: 0.010623s (1000 runs)  speedup:
433.5x
```

```python
# Extract timing arrays for plotting
py_avgs_std = [results_std[L][0] for L in lengths]
cy_avgs_std = [results_std[L][1] for L in lengths]
py_avgs_blk = [results_blk[L][0] for L in lengths]
cy_avgs_blk = [results_blk[L][1] for L in lengths]

# Compute y-scale ratio (Python range / Cython range)
py_max = max(max(py_avgs_std), max(py_avgs_blk))
py_min = min(min(py_avgs_std), min(py_avgs_blk))
py_range = py_max - py_min

cy_max = max(max(cy_avgs_std), max(cy_avgs_blk))
cy_min = min(min(cy_avgs_std), min(cy_avgs_blk))
cy_range = cy_max - cy_min

scale_ratio = py_range / cy_range if cy_range > 0 else 1

# Plot: Python (left) vs Cython (right) with separate y-axes
fig, axes = plt.subplots(1, 2, figsize=(12, 4))
```

```
# Left panel: Python times
ax = axes[0]
ax.plot(lengths, py_avgs_std, marker="o",
        label="Standard (EP = empty)", color="C0")
ax.plot(lengths, py_avgs_blk, marker="s",
        label="Single-block (A·Z·B)", color="C1")
ax.set_xlabel("Sequence length n")
ax.set_ylabel("Time (seconds)")
ax.set_title("Python implementation")
ax.legend()
ax.grid(True, alpha=0.3)

# Right panel: Cython times
ax = axes[1]
ax.plot(lengths, cy_avgs_std, marker="o",
        label="Standard (EP = empty)", color="C0")
ax.plot(lengths, cy_avgs_blk, marker="s",
        label="Single-block (A·Z·B)", color="C1")
ax.set_xlabel("Sequence length n")
ax.set_ylabel("Time (seconds)")
ax.set_title("Cython implementation")
ax.legend()
ax.grid(True, alpha=0.3)

title = (f"NW-flex: Python vs Cython "
         f"(y-scale ratio ~ {scale_ratio:.0f}×)")
fig.suptitle(title, fontsize=13)
plt.tight_layout()
plt.show()

# Print speedup summary
print("\nSpeedups (Python avg / Cython avg):")
print("-" * 50)
print("Standard NW-flex:")
for L in lengths:
    speedup = results_std[L][0] / results_std[L][1]
    print(f"  len = {L:4d}, speedup = {speedup:.1f}x")

print("\nSingle-block NW-flex:")
for L in lengths:
    speedup = results_blk[L][0] / results_blk[L][1]
    print(f"  len = {L:4d}, speedup = {speedup:.1f}x")
```

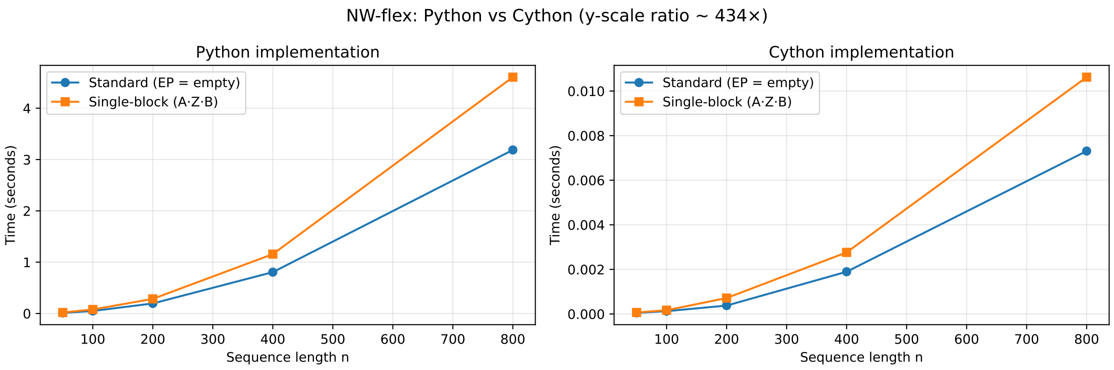

```
Speedups (Python avg / Cython avg):
--------------------------------------------------
Standard NW-flex:
  len =   50, speedup = 249.6x
  len =  100, speedup = 390.6x
  len =  200, speedup = 520.0x
  len =  400, speedup = 425.3x
  len =  800, speedup = 436.0x

Single-block NW-flex:
  len =   50, speedup = 287.9x
  len =  100, speedup = 464.8x
  len =  200, speedup = 398.5x
  len =  400, speedup = 418.8x
  len =  800, speedup = 433.5x
```

##### 5.5.2 Speedup summary

```python
# Calculate speedups
speedups_std = np.array([results_std[L][0] / results_std[L][1] for L in lengths])
speedups_blk = np.array([results_blk[L][0] / results_blk[L][1] for L in lengths])

# Create summary DataFrame
summary_data = []
for n, s_std, s_blk in zip(lengths, speedups_std, speedups_blk):
    summary_data.append({
        "n": n,
        "Standard EP": f"{s_std:.0f}×",
        "Single-block EP": f"{s_blk:.0f}×"
    })

# Add mean row
summary_data.append({
```

```
    "n": "Mean",
    "Standard EP": f"{speedups_std.mean():.0f}×",
    "Single-block EP": f"{speedups_blk.mean():.0f}×"
})

df_summary = pd.DataFrame(summary_data)

print("Speedup factors (Python time / Cython time)")
print()
print(df_summary.to_string(index=False))
```

```
Speedup factors (Python time / Cython time)

   n Standard EP Single-block EP
  50        250×            288×
 100        391×            465×
 200        520×            399×
 400        425×            419×
 800        436×            433×
Mean        404×            401×
```

##### 5.6 Summary

This notebook demonstrated the Cython-backed `run_flex_dp_fast` implementation:

1. **Correctness.** The Cython core produces identical scores, aligned sequences, row jumps, and DP matrices as the pure Python implementation.
2. **EP encoding.** The `ep_to_intervals` function converts list-of-lists EP patterns into a compact interval representation for efficient C iteration.
3. **Speedup.** The Cython implementation achieves significant speedups over pure Python (typically 200–500x depending on sequence length and EP complexity).
4. **Same API.** Both functions take a `FlexInput` and return the same result structure, making them interchangeable.

The high-level alignment helpers in `aligners.py` (e.g., `align_standard`, `align_single_block`, `align_str_phase`) use the Python core by default.

For performance-critical applications, you can construct a `FlexInput` and call `run_flex_dp_fast` directly.

#### 6 Notebook 6: STR Locus Simulation and Pileup

This notebook moves from the theoretical foundations of STR alignment (Notebook 4) to practical simulation and application. We simulate a sequencing experiment by generating reads from a genomic locus and aligning them to a reference template.

#### 6.1 Overview

In real-world scenarios, we align reads sampled from a larger genomic context (the "Full Locus") to a specific target reference (the "Mapping Locus"). This requires **semi-global alignment** where reads may extend beyond the reference boundaries or only partially overlap.

**Goals:** 1. **Define Loci**: Distinguish between the extended genomic region (for sampling) and the restricted reference (for alignment). 2. **Simulate Reads**: Generate synthetic reads with varying STR phases and start positions. 3. **Align with NW-flex**: Use the Cython-accelerated core to align reads to the reference, handling phase shifts and boundary conditions. 4. **Visualize Pileup**: Examine how multiple reads align to the locus, observing phase consistency and gap placement.

**Table of Contents:** 1. Define the STR Locus 2. Simulate Single Read 3. Align Read to Reference 4. Simulate and Align Multiple Reads (Pileup) 5. Summary

#### 6.2 Setup and imports

```
#  Notebook magic: autoreload modules
%load_ext autoreload
%autoreload 2

import numpy as np
import matplotlib.pyplot as plt

# Core NW-flex functions
from nwflex.dp_core import FlexInput
from nwflex.fast import run_flex_dp_fast
from nwflex.aligners import  expand_alignment_with_jumps, get_aligned_bases

# EP pattern builders
from nwflex.ep_patterns import build_EP_STR_phase

# Scoring and plotting
from nwflex.validation import get_default_scoring, random_dna
from nwflex.plot.matrix import plot_flex_matrices
from nwflex.repeats import STRLocus
```

```
The autoreload extension is already loaded. To reload it, use:
  %reload_ext autoreload
```

#### 6.3 Define the STR Locus

We define two related loci:

**1. Full locus (`locus_full`)**: The extended genomic region from which reads are sampled. - `A_full`: Extended left flank (e.g., 200bp) - `R^N`: Repeat region - `B_full`: Extended right flank (e.g., 200bp)

**2. Mapping locus (`locus`)**: The reference used for alignment, with shorter flanks. - `A`: Suffix of `A_full` (e.g., 50bp) - `R^N`: Same repeat region - `B`: Prefix of `B_full` (e.g., 50bp)

Reads sampled from the full locus may extend beyond the mapping locus boundaries, requiring **semi-global alignment** (free gaps at the start/end of the read relative to the reference).

```
Full locus (Sampling):    [--------A_full--------][---R^N---][--------B_full--------]
Mapping locus (Ref):                        [--A--][---R^N---][--B--]
Read Y (Sampled):                              [.......Y.......]
```

```
# Define locus parameters
rng = np.random.default_rng(888)

# Flank lengths
flank_full_len = 500  # Length of extended flanks (for read sampling)
flank_len      = 400   # Length of mapping flanks (for alignment reference)

# Generate random flanks
A_full = random_dna(flank_full_len, rng)
B_full = random_dna(flank_full_len, rng)

# Repeat parameters
R    = "AGG"   # Repeat motif
Nmax = 30      # Reference repeat count

# Extract inner flanks for mapping locus
A = A_full[-flank_len:]   # Suffix of A_full
B = B_full[:flank_len]    # Prefix of B_full

# Create loci
locus_full = STRLocus(A=A_full, R=R, N=Nmax, B=B_full)
locus      = STRLocus(A=A, R=R, N=Nmax, B=B)

print("Full locus (for sampling reads):")
print(f"  |A_full|={len(A_full)}, |R^N|={len(R)*Nmax}, |B_full|={len(B_full)}")
print(f"  |X_full|={locus_full.n}")
print()
print("Mapping locus (alignment reference):")
print(f"  |A|={len(A)}, |R^N|={len(R)*Nmax}, |B|={len(B)}")
print(f"  |X|={locus.n}, s={locus.s}, e={locus.e}")
```

```
Full locus (for sampling reads):
  |A_full|=500, |R^N|=90, |B_full|=500
  |X_full|=1090

Mapping locus (alignment reference):
  |A|=400, |R^N|=90, |B|=400
  |X|=890, s=400, e=490
```

#### 6.4 Simulate Single Read

To simulate a read: 1. **Build a variant** from `locus_full` using phase parameters `(a, b, M)`: - Variant = `A_full + Z* + B_full` where `Z* = suf(R,a) · R^M · pre(R,b)` 2. **Sample a read** by selecting a random start position and extracting `read_len` bases.

We store the variant string for later use (e.g., checking alignment correctness).

```
# Read simulation parameters
read_len = 150

# Generate a variant with specific phase (a, b, M)
# a=1 (1 base from R suffix), b=2 (2 bases from R prefix), M=4 (4 full repeats)
a, b, M = 1, 2, 4
variant = locus_full.build_locus_variant(a, b, M)

print(f"Variant parameters: a={a}, b={b}, M={M}")
print(f"Variant length: {len(variant)}")

# Sample a read that overlaps the repeat region
# Repeat region starts at index 200 in the full locus
start_pos = 450
Y = variant[start_pos : start_pos + read_len]

print(f"Read Y start position: {start_pos}")
print(f"Read Y length: {len(Y)}")
```

```
Variant parameters: a=1, b=2, M=4
Variant length: 1015
Read Y start position: 450
Read Y length: 150
```

#### 6.5 Align Read to Reference

We align read `Y` to the mapping locus reference `X = A + R^N + B` using the Cython-accelerated `run_flex_dp_fast`.

Key parameters: - `build_EP_STR_phase(n, s, e, k)`: Creates the EP pattern allowing row jumps within the repeat region `[s, e)` at motif boundaries (every `k` positions). - `free_X=True`: Read may not cover the full reference (semi-global in X). - `free_Y=True`: Read may extend beyond reference boundaries (semi-global in Y).

The alignment returns the score, aligned strings, path, and any row jumps that occurred. We also request `return_data=True` to visualize the DP matrices.

```
# Build reference and EP pattern (same for all reads to this locus)
score_matrix, gap_open, gap_extend, a2i = get_default_scoring()

X = locus.X
EP = build_EP_STR_phase(locus.n, locus.s, locus.e, locus.k)
```

```python
# Align Y to X
config = FlexInput(
    X=X,
    Y=Y,
    score_matrix=score_matrix,
    gap_open=gap_open,
    gap_extend=gap_extend,
    extra_predecessors=EP,
    alphabet_to_index=a2i,
    free_X=True,
    free_Y=True,
)

# Run Cython alignment
result = run_flex_dp_fast(config, return_data=True)

# Expand alignment to show jumps
X_exp, Y_exp = expand_alignment_with_jumps(X, Y, result.path, result.jumps)

print(f"Alignment score: {result.score}")
print(f"Row jumps: {result.jumps}")
print(f"\nAligned X: {result.X_aln}")
print(f"Aligned Y: {result.Y_aln}")
print("*"*50)
print(f"\nAligned X full: {X_exp}")
print(f"Aligned Y full: {Y_exp}")
```

```
Alignment score: 750.0
Row jumps: [RowJump(from_row=400, to_row=475, col=51, state=1),
RowJump(from_row=489, to_row=491, col=66, state=1)]

Aligned X: GAGCGGGTTTGCTGGCAATCATGACCCTCCCTACAAGTAACTCTAGACGGCGGTCCGTTAATTCTTCTT
GCCTTTAACGTTTCCCGTGCAGCCGCTTTCCCTCAAACCGAGGGTACTTTGTCTTTCGTTTGGCCGCGCGCCCGTATCGA
GGAAGCCCTGGCACACAACAGTTTTCGCACGGAATCGATGTCTCGACCGGGGAATAACGAGGTTAGGTTTTGCGAGACAA
CGTATACTGCATTTGAAGGGTACCATAAGTGTGGGGTTTTGGCAGCAAACAATTAATTCTACTCGCTACTCGCAGACGCA
TTCATATCCGGCTGGTGCTGAGTTAACTGCTCTATGTCTCCAGAGATTCCTTATAGTGATTACAGGGGTAGCAACTTGAA
CGCTTAACCGTGAGGAGGAGGAGGAGGTAAAAAGCAGCCGCTCAATTACTTTTCATGATCTCTCCAAATCGCTTATTCGC
AACGACGCGACAGTACGGTTGGCTAATAAGAACGACACCAACCGTAGCAAAACCATGGCCCAAATAATCTGAAGATATGG
CACTGTAACAGGTCAGGTACCAACTATACTCTAAGGTGCTCGTTCCGGCCGGTGACGCATTGTGAAAAAGAGCCGCAAAA
AAGCTCTTCCTGGTTCCAGAATAGACGCTGATCAGGCCTTAGAGTTAGTAGACACCCCGTGCACATAATGGAAACCCACT
TTTTCAAGGACGCATCTCCGCATAATACACTGGGTCTCGCGGGCACAGTCTCAGCCGCTTTGAGAGCCTTGGACTTGAGG
GACCTACCATTTTGCGCTTTACCTAA
Aligned Y: --------------------------------------------------------------------
-------------------------------------------------------------------------------
-------------------------------------------------------------------------------
-------------------------------------------------------------------------------
----------------------------------------AGAGATTCCTTATAGTGATTACAGGGGTAGCAACTTGAA
```

```
CGCTTAACCGTGAGGAGGAGGAGGAGGTAAAAAGCAGCCGCTCAATTACTTTTCATGATCTCTCCAAATCGCTTATTCGC
AACGACGCGACAGTACGGTTGGCTAATAAGA-------------------------------------------------
--------------------------------------------------------------------------------
--------------------------------------------------------------------------------
--------------------------------------------------------------------------------
--------------------------
**************************************************

Aligned X full: GAGCGGGTTTGCTGGCAATCATGACCCTCCCTACAAGTAACTCTAGACGGCGGTCCGTTAATTC
TTCTTGCCTTTAACGTTTCCCGTGCAGCCGCTTTCCCTCAAACCGAGGGTACTTTGTCTTTCGTTTGGCCGCGCGCCCGT
ATCGAGGAAGCCCTGGCACACAACAGTTTTCGCACGGAATCGATGTCTCGACCGGGGAATAACGAGGTTAGGTTTTGCGA
GACAACGTATACTGCATTTGAAGGGTACCATAAGTGTGGGGTTTTGGCAGCAAACAATTAATTCTACTCGCTACTCGCAG
ACGCATTCATATCCGGCTGGTGCTGAGTTAACTGCTCTATGTCTCCAGAGATTCCTTATAGTGATTACAGGGGTAGCAAC
TTGAACGCTTAACCGTAGGAGGAGGAGGAGGAGGAGGAGGAGGAGGAGGAGGAGGAGGAGGAGGAGGAGGAGGAGGAGGA
GGAGGAGGAGGAGGAGGAGGAGGAGGGTAAAAAGCAGCCGCTCAATTACTTTTCATGATCTCTCCAAATCGCTTATTCGC
AACGACGCGACAGTACGGTTGGCTAATAAGAACGACACCAACCGTAGCAAAACCATGGCCCAAATAATCTGAAGATATGG
CACTGTAACAGGTCAGGTACCAACTATACTCTAAGGTGCTCGTTCCGGCCGGTGACGCATTGTGAAAAAGAGCCGCAAAA
AAGCTCTTCCTGGTTCCAGAATAGACGCTGATCAGGCCTTAGAGTTAGTAGACACCCCGTGCACATAATGGAAACCCACT
TTTTCAAGGACGCATCTCCGCATAATACACTGGGTCTCGCGGGCACAGTCTCAGCCGCTTTGAGAGCCTTGGACTTGAGG
GACCTACCATTTTGCGCTTTACCTAA
Aligned Y full: ----------------------------------------------------------------
--------------------------------------------------------------------------------
--------------------------------------------------------------------------------
--------------------------------------------------------------------------------
----------------------------------------------AGAGATTCCTTATAGTGATTACAGGGGTAGCAAC
TTGAACGCTTAACCGT
--------------------------------------------------------------------------
GAGGAGGAGGAGGAG-GTAAAAAGCAGCCGCTCAATTACTTTTCATGATCTCTCCAAATCGCTTATTCGCAACGACGCGA
CAGTACGGTTGGCTAATAAGA-----------------------------------------------------------
--------------------------------------------------------------------------------
--------------------------------------------------------------------------------
--------------------------------------------------------------------------------
----------------
```

##### 6.5.1 Projecting alignment onto reference coordinates

After expanding the alignment to include jumped positions, we use `get_aligned_bases` to project the alignment onto the reference coordinates. This function returns:

- `ybases`: For each position in the reference X, the corresponding base from Y (or '-' if Y has a gap there). This has the same length as the unaligned reference.
- `xbases`: For each position in the read Y, the corresponding base from X (or '-' if X has a gap there). This has the same length as the unaligned read.

The `ybases` output is particularly useful for **pileup visualization**: it gives us the read's base at each reference position, which we can stack across multiple reads to see coverage and base composition at each locus position.

```python
# Project alignment onto reference coordinates for pileup visualization
# ybases: read bases at each reference position (length = |X|)
# xbases: reference bases at each read position (length = |Y|)
ybases, xbases = get_aligned_bases(X_exp, Y_exp)
print(f"\nAligned bases X: {xbases}")
print(f"Aligned bases Y: {ybases}")
```

```
Aligned bases X: AGAGATTCCTTATAGTGATTACAGGGGTAGCAACTTGAACGCTTAACCGTGAGGAGGAGGAGG
AGGTAAAAAGCAGCCGCTCAATTACTTTTCATGATCTCTCCAAATCGCTTATTCGCAACGACGCGACAGTACGGTTGGCT
AATAAGA
Aligned bases Y: ---------------------------------------------------------------
--------------------------------------------------------------------------------
--------------------------------------------------------------------------------
--------------------------------------------------------------------------------
-----------------------------------------------AGAGATTCCTTATAGTGATTACAGGGGTAGCAA
CTTGAACGCTTAACCGT
--------------------------------------------------------------------------
GAGGAGGAGGAGGAG-GTAAAAAGCAGCCGCTCAATTACTTTTCATGATCTCTCCAAATCGCTTATTCGCAACGACGCGA
CAGTACGGTTGGCTAATAAGA-----------------------------------------------------------
--------------------------------------------------------------------------------
--------------------------------------------------------------------------------
--------------------------------------------------------------------------------
----------------
```

```python
# Normalize Cython data for plotting (convert sentinels to -inf)
# Cython uses a large negative number for -inf, which matplotlib doesn't like␣
↪for color scaling
CYTHON_SENTINEL = -1e30
for arr in [result.data.Yg, result.data.M, result.data.Xg]:
    arr[arr < CYTHON_SENTINEL] = -np.inf

fig_size = (18, 15)
fig = plot_flex_matrices(
        result=result,
        X=X,
        Y=Y,
        s=locus.s,
        e=locus.e,
        marker_size=1,
        figsize=fig_size,
        annotate=False,
        tick_fontsize=4
    )
plt.show()
```

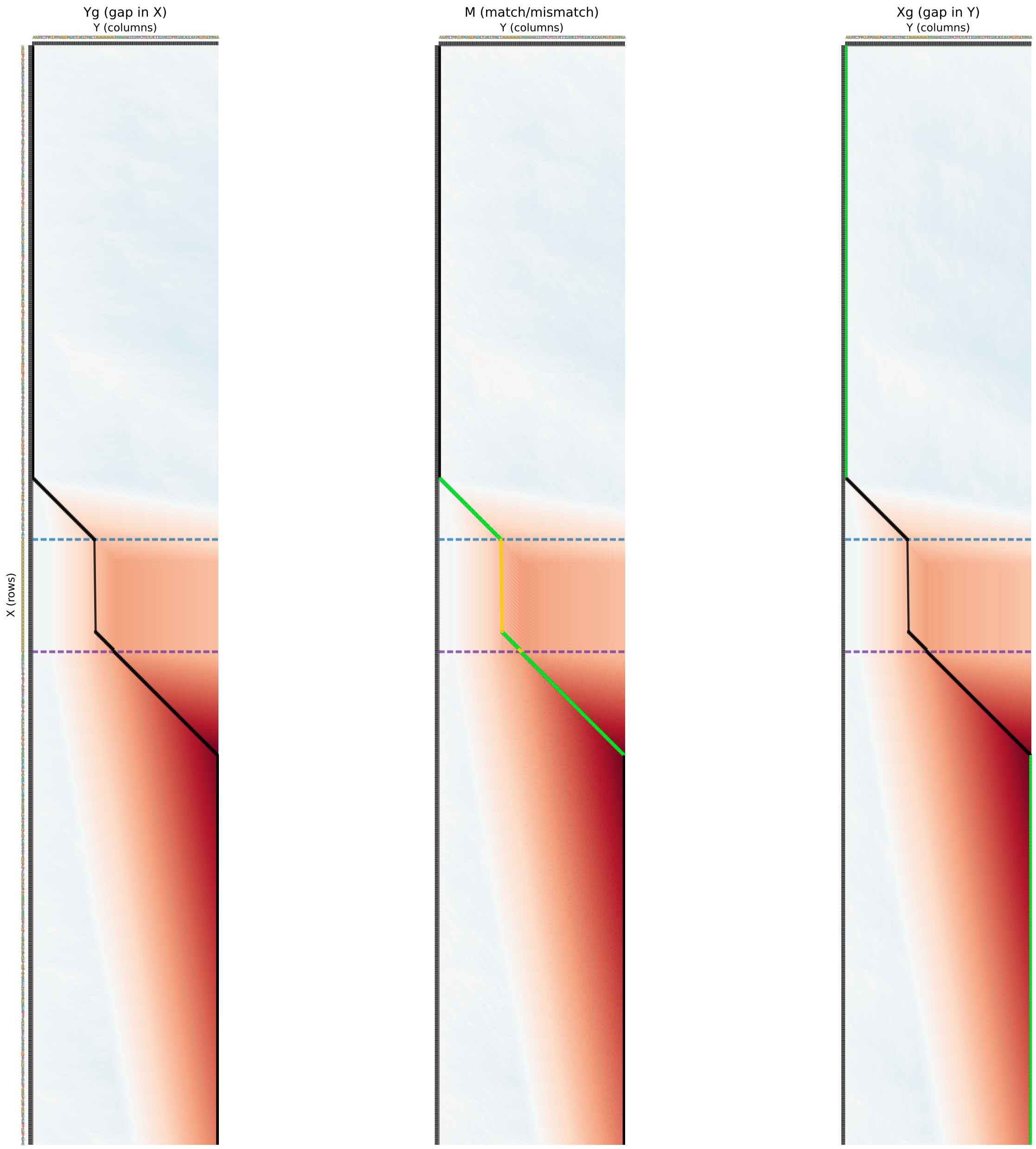

#### 6.6 Simulate and Align Multiple Reads (Pileup)

We now simulate a “pileup” of reads sampled from the full locus.

1. **Sample Reads**: Generate `n_reads` with random start positions from the full locus variant.
2. **Align**: Align each read to the mapping locus `X` using `run_flex_dp_fast`.
3. **Visualize**: Display the aligned sequences to observe how they match the reference, especially around the repeat region.

We reuse the `EP` pattern and scoring configuration for efficiency.

```python
# Simulation parameters
n_reads = 5000
M = 5
a = 1
b = 2

# Build variant and sample read
variant = locus_full.build_locus_variant(a, b, M)

# Generate random start positions and sort them
max_start = len(variant) - read_len
start_positions = rng.integers(0, max_start + 1, size=n_reads)
start_positions = np.sort(start_positions)

# Generate reads and align them
results = []
print(f"Aligning {n_reads} reads...")

for i, start_pos in enumerate(start_positions):
    # Extract read
    Y = variant[start_pos : start_pos + read_len]

    # Configure alignment (reusing X, EP, scoring)
    config = FlexInput(
        X=X,
        Y=Y,
        score_matrix=score_matrix,
        gap_open=gap_open,
        gap_extend=gap_extend,
        extra_predecessors=EP,
        alphabet_to_index=a2i,
        free_X=True,
        free_Y=True,
    )

    # Run fast alignment (no data needed for pileup)
    res = run_flex_dp_fast(config, return_data=False)

    # Expand alignment strings
    X_exp, Y_exp = expand_alignment_with_jumps(X, Y, res.path, res.jumps)

    results.append({
        "id": i,
        "start_pos": start_pos,
        "score": res.score,
        "X_exp": X_exp,
        "Y_exp": Y_exp,
```

```
        "jumps": res.jumps
    })

print("Done.")
```

```
Aligning 5000 reads...

Done.
```

```
from matplotlib.colors import ListedColormap
from matplotlib.patches import Patch

nt_color_map = {
    "A": "#80BCA3",
    "C": "#BDDEF7",
    "G": "#E6AC27",
    "T": "#BF4D28"
    }

base_array = -np.ones(shape=(len(results), len(locus.X)), dtype=int)
for ind, res in enumerate(results):
    ybases, xbases = get_aligned_bases(res["X_exp"], res["Y_exp"])
    base_array[ind] = [a2i.get(base, -1) for base in ybases]

# Create custom colormap
# Map -1 to background, and 0..3 to nucleotide colors
bg_color = "whitesmoke"
sorted_nts = sorted(a2i.keys(), key=lambda k: a2i[k])  # ['A', 'C', 'G', 'T']
colors = [bg_color] + [nt_color_map[nt] for nt in sorted_nts]
cmap = ListedColormap(colors)

# Plot with shifted values (-1 becomes index 0)
fig, ax = plt.subplots(figsize=(12, 6))
im = ax.imshow(base_array + 1, aspect='auto', cmap=cmap, interpolation='nearest')
ax.set_title("Aligned Read Bases at STR Locus")
ax.set_xlabel("Reference Position", fontsize=12)
ax.set_ylabel("Read Index", fontsize=12)

# Create legend with custom patches
legend_elements = []
for nt in sorted_nts:
    legend_elements.append(Patch(facecolor=nt_color_map[nt], edgecolor="black",␣
 ↪label=nt))
legend_elements.append( Patch(facecolor=bg_color, label='gap',␣
 ↪edgecolor="black"))
ax.axvline(x=locus.s - 0.5, color='red', linestyle='--', label='Repeat Start')
ax.axvline(x=locus.e - 0.5, color='blue', linestyle='--', label='Repeat End')
ax.legend(handles=legend_elements, loc='upper right', fontsize=12)
```

```
plt.tight_layout()
plt.show()
```

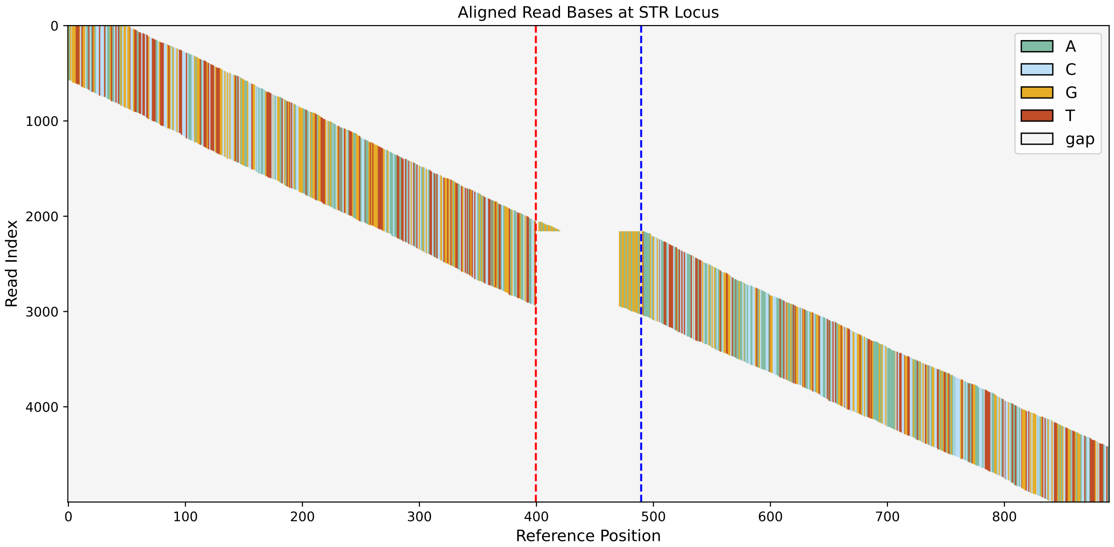

#### 6.7 Summary

In this notebook, we demonstrated how to apply NW-flex to a realistic STR alignment scenario:

1. **Locus Definition**: We distinguished between the **Full Locus** (sampling source) and **Mapping Locus** (alignment reference), creating a realistic semi-global alignment problem.
2. **Semi-Global Alignment**: By setting `free_X=True` and `free_Y=True`, we allowed reads to partially overlap the reference or extend beyond it without penalty at the ends.
3. **Phase-Aware Alignment**: Using `build_EP_STR_phase`, we enabled the aligner to handle repeat copy number differences by "jumping" between repeat units while preserving the motif phase.
4. **Cython Acceleration**: We used `run_flex_dp_fast` to efficiently process multiple reads, showing how the method scales to pileup simulations.

This workflow forms the basis for genotyping STRs from sequencing data, where we must align thousands of reads to a reference locus and infer the consensus repeat sequence.
